# Generative modeling reveals the connection between nuclear morphology and gene expression

**DOI:** 10.64898/2026.01.22.700673

**Authors:** Shuo Wen, Ramon Viñas Torné, Johannes Bues, Camille Lucie Lambert, Nadia Grenningloh, Timothée Ferrari, Elisa Bugani, Joern Pezoldt, Jillian Rose Love, Yinan Wan, Wouter Karthaus, Bart Deplancke, Maria Brbić

## Abstract

How transcriptional programs shape nuclear morphology, and how morphological features in turn reflect and influence cell identity and function, remains poorly understood. Here, we introduce COSMIC, a bidirectional generative framework that enables quantitative decomposition of transcriptional variance reflected in nuclear morphology and morphological variance explained by gene expression. Pretrained on over 21 million segmented nuclei, COSMIC models cross-modal relationships by using a multimodal dataset acquired with IRIS, a technology that captures high-resolution images and transcriptomes from the same single cells. COSMIC learns relationships between gene expression and nuclear morphology, enabling the identification of genes whose expression covaries with nuclear morphological variation. In prostate cancer cells, COSMIC distinguishes chemotherapy-responsive and –resistant cell states through coordinated morphological and transcriptional changes, revealing morphology-associated genes linked to tumour state. Applied to spatial transcriptomics of zebrafish embryogenesis, COSMIC reveals how local cellular neighborhoods influence morphology-expression relationships. These results demonstrate that generative modeling powered by paired single-cell measurements can quantify the information shared between nuclear morphology and gene expression, opening new avenues for mechanistic discovery in both basic and translational cell biology.

## Introduction

Deciphering the extent to which transcriptional programs encode cellular form, and, conversely, how morphological variation reflects underlying gene expression remains an open question in cell biology. Single-cell RNA sequencing (scRNA-seq) (1; 2) provides high-resolution molecular profiles, while imaging technologies (3; 4; 5; 6) capture the detailed morphological and structural organization of individual cells. Bridging these complementary modalities is essential for deciphering how transcriptional programs determine cellular form and, conversely, how morphological features affect cell identity, function, and responses to perturbations.

Multimodal computational frameworks could enable translation between transcriptomic and morphological information, revealing how gene-expression programs give rise to cellular phenotypes and how morphological variation reflects underlying molecular states. Over the past decade, substantial progress toward this vision has been driven by advances in both biotechnology and computational modeling. Large-scale programs such as GTEx (7) and TCGA (8) provide extensive molecular and imaging data that have powered methods for inferring transcriptomic information from histology images (9; 10; 11; 12; 13). At the single-cell level, prior work has established that nuclear and chromatin images contain information about transcriptional and cellular states, supporting the idea that morphology and gene expression share measurable biological signal (14; 15; 16). Imaging-based spatial transcriptomics (17; 18; 19; 20) and single-cell multimodal technologies, such as STAMP (21) and Patch-seq (22), enable coupling of images with molecular readouts at the single-cell resolution, but remain limited either in their sensitivity and gene coverage, or the throughput needed for training AI models. Using such data, computational methods (23) attempted to predict cellular morphology from transcriptomic profiles, but fell short of capturing fine-grained morphological differences between cells. Due to these limitations, quantifying the information shared between cellular morphology and gene expression remains challenging.

To quantify and model the information shared between nuclear morphology and gene expression at single-cell resolution, we developed COSMIC (**C**ross-m**O**dal generation between **S**ingle-cell RNA-seq and **MIC**roscopy images), a bidirectional generative framework that enables translation between single-cell transcriptomes and nuclear morphology. To pair modalities, we built a collection of 17,109 human and 9,039 mouse single cells using the IRIS platform (24), a microfluidic system that simultaneously acquires high-resolution microscopy images and high-quality transcriptomic profiles from the same individual cells.

We focus on nuclear morphology, which is known to be linked with transcriptional activity, chromatin organization, disease states, and can be used to identify pathologies (25). For example, Hutchinson-Gilford progeria syndrome is associated with disruption of the nuclear envelope and misshapen nuclei with nuclear blebs (26), while cellular senescence is accompanied by changes in nuclear morphology, chromatin organization, and heterochromatin structure (27; 28). These examples illustrate that nuclear morphology can reflect underlying cellular and transcriptional states. To capture distinct features of nuclear morphology, we pretrained an image encoder on 21.8 million segmented nuclear images to capture diverse morphological representations. COSMIC couples this nuclear morphology encoder with transcriptomics encoders (29; 30) to enable cross-modal generation using a diffusion model conditioned on the unimodal representations of transcriptome and nuclear morphology.

We show that discriminative morphological features can be generated from transcriptomic profiles with high accuracy, and that nuclear morphology, in turn, captures biologically meaningful variation in transcriptomic states. This bidirectional relationship spans multiple levels of biological information, capturing discrete differences between cell types as well as continuous variation associated with cell-cycle progression. COSMIC identifies genes and gene programs associated with nuclear morphology, revealing both established regulators and previously uncharacterized candidate genes whose expression is linked to morphological variation. Applied to the multi-modal IRIS dataset of prostate cancer cells treated with the chemotherapeutic agent Docetaxel, COSMIC identifies genes whose expression is associated with nuclear morphological differences between treatment-responsive and non-responsive cells. COSMIC is a general framework applicable to spatial transcriptomics technologies (17; 18) where it can help to reveal how local cellular neighborhoods influence morphology-expression relationship. We demonstrate it on the zebrafish embryo weMERFISH data (31) where COSMIC identified global tissue-level as well as local within-tissue nuclear morphology-associated genes.

## Results

### Overview of COSMIC

To model shared information between morphology and transcriptome, we developed COSMIC, a bidirectional generative framework that enables the prediction of gene expression profiles from microscopy images of nuclei, and, conversely, the reconstruction of nuclear images from transcriptomic data. To build a general-purpose encoder of nuclear morphology, we trained a vision encoder on microscopy images of cellular nuclei, referred to as the COSMIC nuclear encoder. We compiled a dataset of 21, 784, 309 nuclear images by segmenting and isolating individual nuclear crops from 50, 377 whole-well Hoechst-stained microscopy images from three studies, including JUMP-CP (32), Stojic-lncRNAs (33), and Pascual-Vargas-RhoGTPases (34) (Fig. 1a; Methods). We then used this large amount of individual nuclear images to pretrain an encoder of nuclear morphology. We trained the model in the encoder-decoder fashion using a masked autoencoder (MAE) (35) objective in which random patches of each input image were masked and the model learned to reconstruct them, encouraging the encoder to capture semantically rich features of nuclear structure (Fig. 1b; Methods).

**Figure 1.**
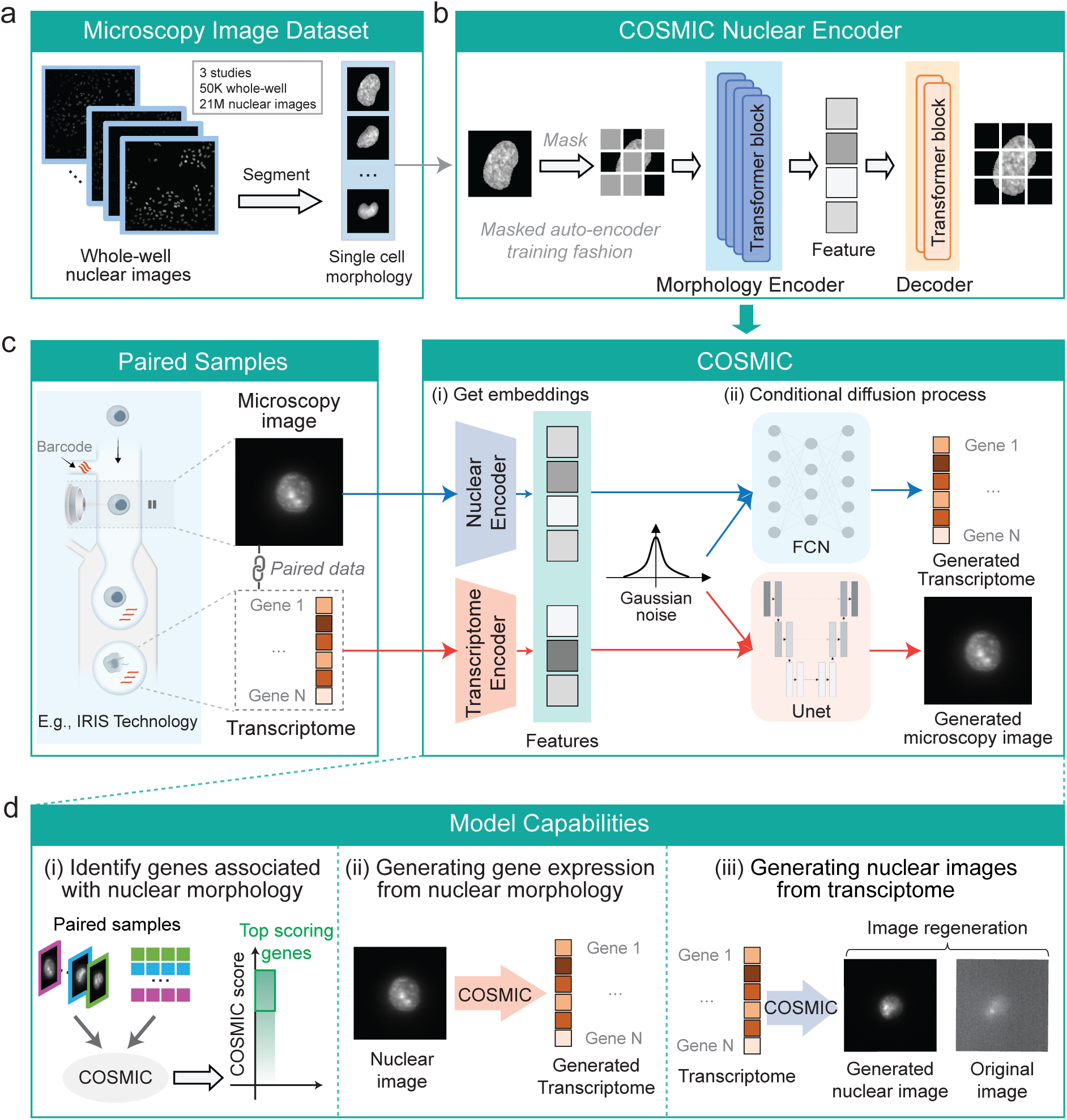
Overview of COSMIC, a generative model for bidirectional cross-modal generation between nuclear morphology and transcriptomic profiles. (a) Preparation of a dataset of single-cell nuclear images. We collected *∼*50.3K whole-well microscopy images stained with Hoechst dyes from three studies. Whole-well images were segmented to obtain single-cell images, resulting in a total of *∼*21.8M nuclear images. **(b)** To learn meaningful nuclear morphological representations of cells, we pretrained a masked auto-encoder (MAE) on collected nuclear images. During pretraining, random patches of the input images were masked, and the model was trained to reconstruct the missing regions. **(c)** Training pipeline of COSMIC. Conditioned on cell features extracted from microscopy images and single-cell transcriptomes obtained from pretrained unimodal models, COSMIC performs conditional generation using two conditional diffusion models. The diffusion models are trained on paired samples generated with IRIS technology to learn cross-modal relationships between nuclear morphology and the transcriptome. **(d)** COSMIC’s cross-modal capabilities support a range of downstream tasks, including *(i)* identifying genes associated with nuclear morphology, *(ii)* generating gene expression from nuclear morphology at the single-cell level, and *(iii)* generating nuclear images from the transcriptome profiles of single-cells, including the generation of biologically consistent nuclear images under noisy imaging conditions.

COSMIC builds on the nuclear encoder to capture information about nuclear morphology, and leverages existing embedding models to encode the transcriptomic profile of cells (29; 30; 36). These unimodal encoders provide conditioning representations to the conditional diffusion models (37; 38) trained on multimodal data of paired gene expression profiles and nuclear images (Fig. 1c). We leveraged paired multi-modal measurements using the IRIS platform (24), a microfluidic system that acquires high-resolution microscopy images and co-encapsulates each imaged cell in a droplet with a unique barcode for transcriptomic profiling, enabling morphological features to be directly linked to each cell’s transcriptome. Another approach for obtaining paired single-cell morphology–transcriptome measurements is spatial transcriptomics (ST) (17; 18) and COSMIC is applicable to both. The conditional diffusion model is trained on paired nuclear images and gene expression profiles, learning to capture correspondences between transcriptomic and morphological states. By conditioning generation on either modality, COSMIC synthesizes nuclear images from gene expression and predicts transcriptomic profiles from morphology, thereby enabling bidirectional translation across single-cell modalities and quantification of the strength of relationship between nuclear morphology and transcriptome (Methods). The unified design of COSMIC supports a range of model applications, including identifying morphology-associated genes, generating gene expression profile from nuclear morphology, and generating nuclear images from transcriptomes, with clean image recovery as an application (Fig. 1d).

### COSMIC quantifies the relationship between nuclear morphology and transcriptomic state

To determine how strongly nuclear morphology reflects transcriptional state and conversely how transcriptional state is reflected in nuclear morphology, we evaluated how well COSMIC generates each modality from the other. Image-to-transcriptome generation quantifies how much information nuclear morphology contains about transcriptional state, whereas transcriptome-to-image generation quantifies how predictive gene-expression profiles are for reconstructing biologically meaningful nuclear morphology.

We first used COSMIC to generate transcriptomic profiles from microscopy images of single cells (Fig. 2a). COSMIC encodes single-cell nuclear microscopy images using a nuclear encoder and uses them as a condition to predict the corresponding transcriptomic profiles. We trained COSMIC on 4, 520 paired nuclear image-transcriptome samples of mouse cells generated using IRIS (Supplementary Note 1) and evaluated it on an independent set of 4, 519 mouse cells. Specifically, we profiled mouse embryonic fibroblasts (fibroblasts, cell line 3T3), macrophage-like cells (macrophage, cell line RAW), CAR-engineered T cells derived from the A20 B-cell lymphoma model (B-cell lymphoma, cell line CAR A20), and primary naive CD8+ T cells (naive CD8). The dataset comprises batches generated in this work and batches from the companion manuscript (24) (Supplementary Table 1).

**Figure 2.**
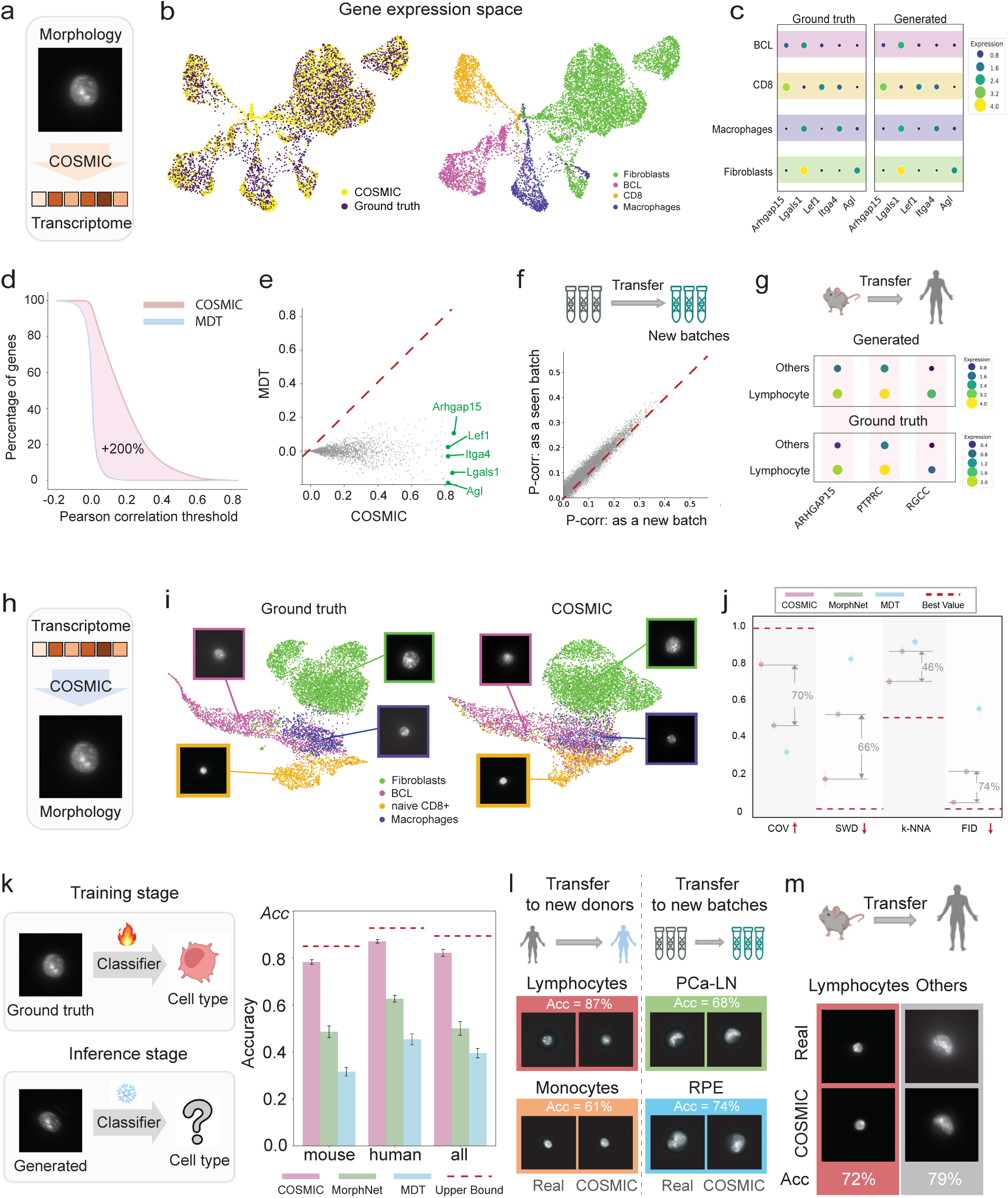
COSMIC generates biologically meaningful predictions. **(a)** COSMIC generates transcriptomic profiles from single-cell nuclear images obtained with the IRIS technology. **(b)** Comparison between ground truth and generated transcriptomes on (*n* = 4, 519) mouse cells in the UMAP space. Colors indicate ground truth and generated data (left), and different cell types (right). BCL refers to B-cell lymphoma. **(c)** Average gene expression of the five genes with the highest Pearson correlation across different cell types. Ground truth expression (left) and COSMIC predictions (right). **(d)** Cumulative percentage of genes exceeding a given Pearson correlation coefficient between generated and ground-truth transcriptomic profiles. COSMIC is compared to multi-domain translation (MDT). **(e)** Comparison of Pearson correlation coefficient between COSMIC and MDT for individual genes. Five genes with the highest Pearson correlation are highlighted in green. **(f)** COSMIC generalizes to unseen batches. We trained two conditional generation models on human samples: one model on all cell types and batches, and another one excluding one batch for the RPE1 cell line, and compared their performance. **(g)** COSMIC generalizes across species. We trained COSMIC on mouse cells and applied the model to human cells that have not been seen by the model. The overlapping cells between mouse and human are lymphocytes (with CD8 cells in mouse also classified as lymphocytes). The dot plot of average gene expression of lymphocyte differentially expressed genes shows that COSMIC can correctly extrapolate the expression of these genes from mouse to human lymphocytes. ‘Others’ refers to the remaining cell types, including PCa-Br and RPE. **(h)** Application of COSMIC to generate nuclear images given single-cell transcriptomic profile using IRIS data. **(i)** Embedding space comparison between ground truth images and COSMIC generated images on the (*n* = 4, 519) mouse cells. Embeddings are extracted using the pretrained nuclear encoder and visualized using UMAP. Colors indicate cell types. An example image for each cell type is shown in a box. **(j)** Quantitative evaluation of the quality of generated images of IRIS mouse cells. We assessed the similarity between ground-truth and generated distributions using coverage rate (COV), sliced Wasserstein distance (SWD), *k*-nearest neighbor accuracy (k-NNA), Fréchet inception distance (FID). Higher COV values indicate better coverage, lower SWD and FID values suggest higher fidelity, and k-NNA values closer to 0.5 reflect better sample diversity. Red dashes indicate the best value for each metric. The numbers represent the improvement achieved by COSMIC over the best alternative baseline. Error bars represent the standard deviation across five independent rounds of generation. **(k)** Evaluation of COSMIC and alternative methods on the cell type classification task using on the mouse and human datasets. We trained a classifier on ground-truth cell images and cell type annotations. We froze the classifier and applied it to the generated images of different methods without further training. Higher classification accuracy indicates better conditional generation quality. Error bars represent standard deviation across five independent rounds of generation. The red dashed line represents the upper bound obtained by evaluating classification accuracy on the ground-truth images. **(l)** COSMIC generates high-quality nuclear images for unseen batches. We trained COSMIC on human cells from four cell types, including lymphocytes, monocytes, prostate cancer cells from a lymph node metastasis (PCa-LN), and retinal pigment epithelial cells (RPE), holding out one batch per cell type. For lymphocytes and monocytes, held-out batches correspond to new donors; for PCa-LN and RPE, they correspond to independent experimental replicates. After training, we generated images for these unseen batches and evaluated the quality of the generated images by testing whether a classifier trained on ground-truth images could accurately distinguish the four cell types in the generated set. **(m)** COSMIC generates high-quality nuclear images across species, from mouse to human. We trained COSMIC on mouse data and evaluated its cross-species generalization on human samples. COSMIC accurately synthesized human lymphocytes cells, with 72% correctly classified as lymphocytes by the cell type classifier.

We first visualized the gene expression space of both generated and ground truth profiles. The resulting UMAP (39) projections revealed a substantial overlap between the COSMIC-generated transcriptomes and the real data in the gene expression space, and the clusters corresponding to different cell types were preserved (Fig. 2b). COSMIC produced biologically valid transcriptomic profiles that respected inter-cell type differences and the overall transcriptomic landscape. We observe similar results when applying COSMIC to human cells (8, 555 train samples and 8, 554 test samples) from the IRIS technology (Supplementary Fig. 1). The human data comprises peripheral blood mononuclear cells (PBMCs, including lymphocytes and monocytes) and established cell lines: RPE1 (retinal pigment epithelial cells; named as RPE), C4-2B (prostate cancer cells from a lymph node metastasis; named as PCa-LN), and DU145 (prostate cancer cells from a brain metastasis; named as PCa-Br). To understand the biological relevance of these predictions, we identified the top five most accurately predicted genes for each cell type and plotted their average expression across cell types (Fig. 2c). These gene signatures respected the expected enrichment patterns, confirming that COSMIC could recover cell type-specific expression programs for a subset of genes from nuclear morphology alone.

To evaluate gene-level fidelity, we computed the Pearson correlation coefficient between generated transcriptome profiles and their ground truth counterparts for all genes (Supplementary Note 5). We compared COSMIC to Multi-Domain Translation (MDT) (15), which aligns unpaired samples across modalities via representation space matching and does not require paired image-transcriptome measurements during training (Supplementary Note 2). Using COSMIC, 12.5% (*n* = 1, 814 out of 14, 455 genes) had a correlation coefficient higher than 0.4 (Fig. 2d). In contrast, none of the genes predicted by MDT exceeded this threshold. These correlations quantify the fraction of transcriptomic variation that can be predictably recovered from nuclear morphology. As expected, gene expression is only partially encoded in nuclear morphology, with additional variation arising from biological processes and measurement noise that are not observable from morphology alone. This analysis also revealed that many genes are well predicted from nuclear images (73.7% of the total genes had a BH-FDR adjusted p-value *<* 0.05). When comparing individual genes, COSMIC was particularly predictive for cell type-specific marker genes. For example, some marker genes, including *Lef1*, *Itga4*, *Lgals1*, and *Arhgap15*, displayed correlation coefficients nearing 0.8 across all cells, suggesting that morphology encodes precise expression patterns for a subset of genes with strong phenotypic associations (Fig. 2e). We additionally compared COSMIC with two simple baselines for image-to-transcriptome prediction: *(i)* a regression model trained on IRIS data using CellProfiler-derived morphology features (40) and *(ii)* an unpaired-data COSMIC baseline in which transcriptomes and nuclear images were randomly paired before cross-modal training (Supplementary Note 2). COSMIC outperformed both baselines, indicating that its performance is not explained by simple hand-engineered nuclear features or by dataset-level structure learned from unpaired samples (Supplementary Fig. 2).

We further assessed COSMIC’s ability to correctly model nuclear morphology-transcriptome relationships on unseen batches. In this experiment, we trained two separate models: one on all available human IRIS data, and another with one batch held out for the RPE1 cell line. For both models, we evaluated gene-wise correlation on the test set and found that the Pearson correlation patterns were highly consistent (Fig. 2f). This demonstrates that COSMIC did not overfit to individual batches and was capable of extracting generalizable morphological features predictive of transcriptional state. Besides, we tested cross-species generalization by training COSMIC exclusively on mouse data and applying it to generate transcriptomes from the morphology of human lymphocyte cells. Despite species differences, the generated human expression profiles preserved the expected differential gene expression patterns in lymphocyte cells, with accurate prediction of lineage-defining markers such as *ARHGAP15*, *PTPRC*, and *RGCC* (Fig. 2g). This result shows the opportunities to model evolutionarily conserved relationships between nuclear morphology and gene expression.

To dissect the factors contributing to COSMIC’s performance, we evaluated the effect of the pretrained COSMIC nuclear encoder by comparing it with different pretrained image encoders (ImageNet MAE (35), ImageNet ResNet, and OpenPhenom (41)), demonstrating that the nuclear encoder provides the strongest representation for predicting gene expression from nuclear morphology (Supplementary Fig. 3). Performance improved with more pretraining images but the gains were small at large scales (Supplementary Fig. 4). Evaluating the effect of smaller fractions of the IRIS training data showed that performance remained stable when using 60–100% of the total number of cells but decreased at smaller fractions (Supplementary Fig. 5).

We then quantified the ability of COSMIC to generate nuclear images from transcriptomic profiles of cells (Fig. 2h) using the same dataset and train-test split employed for generating single-cell transcriptomic profiles from nuclear images. As a transcriptomic encoder, we used scVI (29) and conditioned the diffusion model on its embeddings, guiding the denoising process to synthesize high-resolution nuclear images. To assess performance, we embedded both real and generated nuclear images using the nuclear encoder and visualized them in the 2D UMAP space. This visualization revealed strong agreement between the distributions of real and synthetic samples, with clear clustering by cell type (Fig. 2i). We compared COSMIC to two deep learning baselines: MorphNet (23), a generative adversarial network based model trained to synthesize images from gene expression, and Multi-Domain Translation (MDT) (15) (Supplementary Note 2). In contrast to COSMIC, MorphNet and MDT exhibited worse alignment between their embedding spaces and the ground truth (Supplementary Fig. 6).

Since UMAP visualization alone cannot establish biological relevance, we next quantified image-generation quality using multiple complementary metrics. To quantify the overall quality of the generated images, we use four metrics widely used in computer vision: coverage (COV) (42), sliced Wasserstein distance (SWD) (43), k-nearest neighbor accuracy (*k*-NNA) (44), and Fréchet inception distance (FID) (Supplementary Note 5). COV measures how well the generated samples span the real data distribution; SWD and FID assess statistical divergence between real and generated distributions; and k-NNA evaluates how well the real and generated samples are intermixed. Across these complementary metrics, COSMIC substantially outperformed both MorphNet and MDT (Fig. 2j), achieving 46% to 74% better performance compared to the best alternative method MorphNet. Similarly, we observed large improvements when applying COSMIC to human cells (8, 555 train samples and 8, 554 test samples) from the IRIS technology (Supplementary Fig. 7). COSMIC-generated images also preserved interpretable, feature-level morphology differences observed in real microscopy images: CellProfiler-derived nuclear morphology features from COSMIC-generated images recovered a high fraction of the CellProfiler features that were discriminative in real images (Supplementary Fig. 8). These results support that the generated images more closely match the real nuclear-image distribution than those produced by alternative methods and can be used to effectively quantify the strength of the morphology-transcriptome relationship.

We next evaluated whether COSMIC-generated images preserve cell type-specific information. To this end, we trained a convolutional neural network (CNN) classifier to predict cell types using real images and applied it on synthetic images generated by COSMIC, MDT, and MorphNet (Supplementary Note 2). The cell-type classifier has never seen generated images during training and has been solely trained on the training set of real images. On the IRIS mouse dataset, we found that a classifier applied to COSMIC’s generated images achieved an average classification accuracy of 78%, close to the upper bound of 85% observed on the test set of real data (Fig. 2k). In contrast, images generated by alternative methods MorphNet and MDT achieved significantly lower performance (48% and 32%, respectively). We observed comparable performance on a human dataset, where the model achieved 88% accuracy, with an upper bound of 93% on real images. We obtained similar performance gains using other metrics, and COSMIC produced class-confusion patterns closest to those observed for real images (Supplementary Fig. 9). This demonstrates that the COSMIC-generated images retained a strong conditional signal and preserved discriminative features of cell types.

We evaluated out-of-distribution of COSMIC performance with a held-out batch design, using the same CNN classifier trained on real images for evaluation. COSMIC was trained on mixed data from all human groups while withholding one batch from lymphocytes, monocytes, RPE, and PCa-LN; PCa-Br had only one batch and could not be held out. On these held-out batches, generated images enabled accurate cell type classification, achieving 87% accuracy for lymphocytes, 61% for monocytes, 68% for PCa-LN, and 74% for RPE. (Fig. 2l). This probes two generalization regimes: for PBMCs (lymphocytes and monocytes), each batch corresponds to a distinct donor, so COSMIC is evaluated on unseen donors; for cell lines such as RPE1 (RPE) and C4-2B (PCa-LN), the genetic background is fixed across batches, so COSMIC is evaluated on unseen experimental replicates. Beyond experimental replicates and donor variability, we further tested COSMIC’s ability for cross-species generalization. We trained COSMIC exclusively on mouse data and applied it to human gene expression profiles to generate microscopy images. Despite substantial shifts in both transcriptomic and imaging distributions, COSMIC successfully synthesized human lymphocyte images that were classified with 72% accuracy (Fig. 2m). By leveraging representations from single-cell foundation models to embed cells, COSMIC can generalize to new experimental replicates, donors, and species in a zero-shot fashion. Using UCE (30), a pretrained single-cell foundation model, as an encoder, we tested COSMIC in this setting. The results demonstrate COSMIC’s effectiveness even in this more difficult setting, as well as its compatibility with existing single-cell foundation models (Supplementary Fig. 10).

These results support that COSMIC quantifies the information shared between nuclear morphology and the transcriptome in both directions, revealing that one can infer substantial information about cell state from the nuclear morphology only. Image-to-transcriptome generation has an immediate practical value in settings where transcriptomic profiling measurements are costly or where only microscopy data is available, such as clinical pathology or large-scale morphological screens. Transcriptome-to-image generation has a practical value in recovering clean morphology from noisy image acquisitions. To demonstrate this, we applied COSMIC to restore morphology-derived feature differences that were suppressed by imaging noise and reduced noise-derived feature artifacts. Applied to a subset of IRIS CD8 cell images affected by substantial imaging noise, COSMIC enabled the recovery of morphological features supressed by noise (Supplementary Fig. 11).

### COSMIC learns continuous cell cycle dynamics from transcriptomes and morphology

While COSMIC effectively captures cell type identity through relationships between nuclear morphology and transcriptomic profiles, we next explored whether it can also encode biological information beyond discrete cell types and recover continuous biological processes.

We conducted a within-cell-type permutation to assess whether COSMIC uses morphology-expression correspondence beyond cell type identity (Fig. 3a): for each cell type, we randomly reassigned transcriptomic profiles to microscopy images, preserving the exact sets of images, expression profiles, and labels while breaking the true pairings, thereby removing any linkage between morphology and gene expression within cell types. If the model relied only on cell type information to make predictions, gene-level performance would be similar between original and permuted data, which we indeed observe for the marker genes (Fig. 3b). In contrast, we observed that the performance of a subset of genes is affected by this permutation, indicating that these genes were useful for characterizing nuclear morphological differences within cell types. The performance of some of these genes was affected across multiple cell types, indicating shared processes across different cell types. For example, such genes included cytoskeletal and contractile regulators (*Acta1*, *Acta2*, *Tubb4a*, *Tubb2b*, and *Arap2*), which modulate cytoskeletal tension and its transmission to the nucleus, thereby altering nuclear morphology (45). We further focused on cell type-specific genes, *i.e.*, genes affected within specific cell types. In fibroblasts, we observed that affected genes included several cell-cycle genes, such as *Top2a* and *Prc1* (Fig. 3c). Beyond cell cycle regulation, we identified affected genes that function as key modulators of the contractile cytoskeleton, like *Acta2* (smooth muscle actin) and *Cald1* (caldesmon). The increased expression of these genes suggests enhanced contractility and stress-fiber assembly, providing a plausible mechanism for the observed changes in nuclear morphology. Consistent with the fibroblast results, permutation reduced performance in other cell types, enabling characterization of genes that are most affected by the permutation (Supplementary Fig. 12).

**Figure 3.**
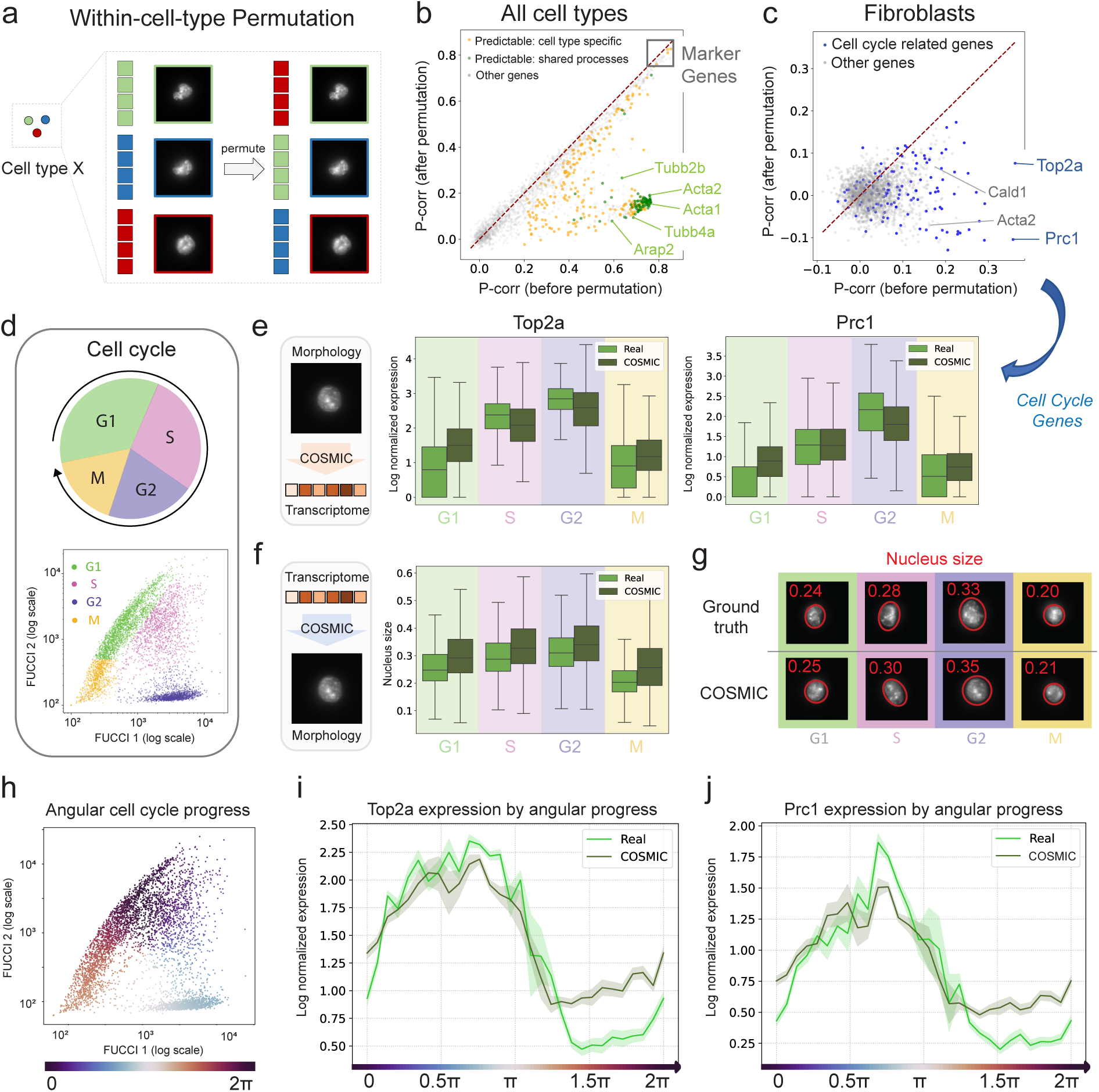
COSMIC accurately captures transcriptomic and morphological differences during cell cycle in mouse fibroblast cells. (a) To test whether COSMIC captures biological information beyond cell type identity, we permuted the pairings between gene expression profiles and nuclear images within each cell type. This procedure disrupts gene expression-morphology relationships while preserving the underlying cell type information, as permutations were restricted to within-cell-type groups. **(b)** Performance drop after within-cell-type permutation highlights biological signals beyond cell type. Across all cell types, genes with the strongest performance drop are primarily cell type specific predictable genes, while marker genes retain similar performance since the permutation does not remove cell type information. **(c)** Within fibroblasts, within-cell-type permutation leads to a performance drop for cell cycle related genes such as *Top2a* and *Prc1*, as well as for other genes like key regulators of the contractile cytoskeleton, including *Acta2* and *Cald1*. **(d)** Quantification of cell cycle progression using the FUCCI reporter. We simultaneously capture expression profiles, microscopy images, and FUCCI reporter intensities, and use the FUCCI signal to annotate cells into four cell cycle phases (G1, S, G2, M). **(e)** Transcriptomes predicted by COSMIC recapitulate cell cycle information. The box plot illustrates the distribution quartiles of *Top2a* expression levels across different cell cycle phases (G1: *n* = 1, 631 cells, S: *n* = 1, 036 cells, G2: *n* = 1, 366 cells, M: *n* = 748 cells.) Boxes depict distribution quartiles, with the center line corresponding to the median, and whiskers span 1.5 times the interquartile range. **(f)** Images generated by COSMIC recapitulate cell cycle information. Distribution of nucleus sizes, a morphological trait strongly linked to the cell cycle, for each cell cycle phase. Boxes depict distribution quartiles, with the center line corresponding to the median, and whiskers span 1.5 times the interquartile range. **(g)** Examples of ground truth (top) and images generated by COSMIC (bottom) for each phase. Ground truth examples are chosen based on median nucleus size, while COSMIC-generated examples are generated from the transcriptomes of the same cells. Images are annotated with relative nucleus sizes normalized to the largest nucleus observed in mouse cells. **(h)** Cell cycle can also be characterized using a continuous circular trajectory obtained from FUCCI intensities (Methods). The scatter plot visualizes cell cycle progression in the FUCCI space for fibroblast mouse data obtained via the IRIS technique. **(i, j)** The COSMIC-inferred expression profiles correctly recapitulate continuous expression changes of cell-cycle genes (i) *Top2a*, and (j)*Prc1*.

Focusing on mouse fibroblasts, we tested whether COSMIC can recover known biological dynamics without supervision, focusing on the cell cycle. The availability of Fluorescence Ubiquitin Cell Cycle Indicator (FUCCI) (46; 47) reporter signals in this dataset allowed us to explicitly validate the inferred cell cycle profiles (Fig. 3d).

We first analyzed COSMIC’s ability to recapitulate phase-dependent gene expression when inferring transcriptomes from nuclear images. As a representative marker, we examined the expression of *Top2a*, a gene encoding a DNA topoisomerase enzyme essential for chromosomal segregation (48). In the ground truth scRNA-seq data, *Top2a* showed the expected expression pattern, peaking in G2 phase and remaining low in G1 (Fig. 3e). COSMIC-generated transcriptomes preserved this phase-specific trend, with predicted expression levels closely mirroring the real data across all phases. Comparable results were obtained for *Prc1*, a cytokinesis regulator, where COSMIC accurately reproduced its elevated expression during G2/M and low expression during G1. Other genes, including *Cdk1*, *Ppp3ca*, *Kif20b*, and *Cenpf*, showed consistent phase-dependent patterns in the predictions (Supplementary Fig. 13). These results demonstrate that COSMIC can decode dynamic gene expression programs from nuclear morphology, likely by leveraging features such as chromatin condensation and nucleus size, which vary systematically throughout the cell cycle.

We next examined whether COSMIC could capture morphological changes associated with cell cycle progression when generating images from transcriptomes. Nuclear morphology is known to vary across phases, with G1-phase cells typically exhibiting smaller nuclei and G2-phase cells showing enlarged shapes due to DNA replication and preparation for mitosis. We quantified nuclear size in both real and COSMIC-generated images, stratified by FUCCI phase annotations. COSMIC reproduced the expected trend: the mean nuclear size increased from G1 to G2 and then declined slightly in M phase, closely matching the real measurements (Fig. 3f, g). These differences were not only statistically significant (Holm-corrected pairwise Welch’s t-test; p-value *<* 0.0001), but also visually apparent in representative generated images, which reflected consistent changes in nuclear size across the cell cycle.

Beyond the four discrete cell cycle phases, we further modeled the cell cycle as a continuous circular trajectory. To explore whether COSMIC preserved continuous biological gradients, we computed the angular cell cycle progression using the FUCCI intensity space and plotted expression of cell cycle genes across this inferred trajectory (Fig. 3h, Supplementary Note 5). In the ground truth data, *Top2a* exhibited a smooth, sinusoidal curve, increasing sharply after G1 phase and peaking during G2 phase. The COSMIC-generated expression profiles displayed a highly similar trend (Fig. 3i). We observe similar performance for *Prc1* (Fig. 3j) as well as other cell cycle genes including *Cdk1*, *Ppp3ca*, *Kif20b* and *Cenpf* (Supplementary Fig. 14), further validating the ability to model continuous, cyclic processes. Notably, this result emerged without explicit supervision on the angular trajectory, suggesting that COSMIC implicitly learned the progression structure from paired data. These results indicate the potential to reconstruct continuous biological processes from snapshot data in both imaging and sequencing modalities.

### COSMIC proposes shared and cell-type specific morphology-associated genes and gene programs

Given paired morphology and transcriptome measurements, COSMIC assigns each gene a morphology-association score that quantifies how strongly its expression distribution depends on nuclear morphology (Fig. 4a). The score compares image-conditioned expression distributions generated by COSMIC with a morphology-independent reference distribution aggregated across samples, capturing both shifts in expected expression and broader distributional changes. Specifically, mean shifts are measured using correlation-based scores, while broader distributional shifts are measured using hurdle mutual information (49), which accounts for zero-inflation (Methods). The resulting scores enable systematic prioritization of morphology-associated genes.

**Figure 4.**
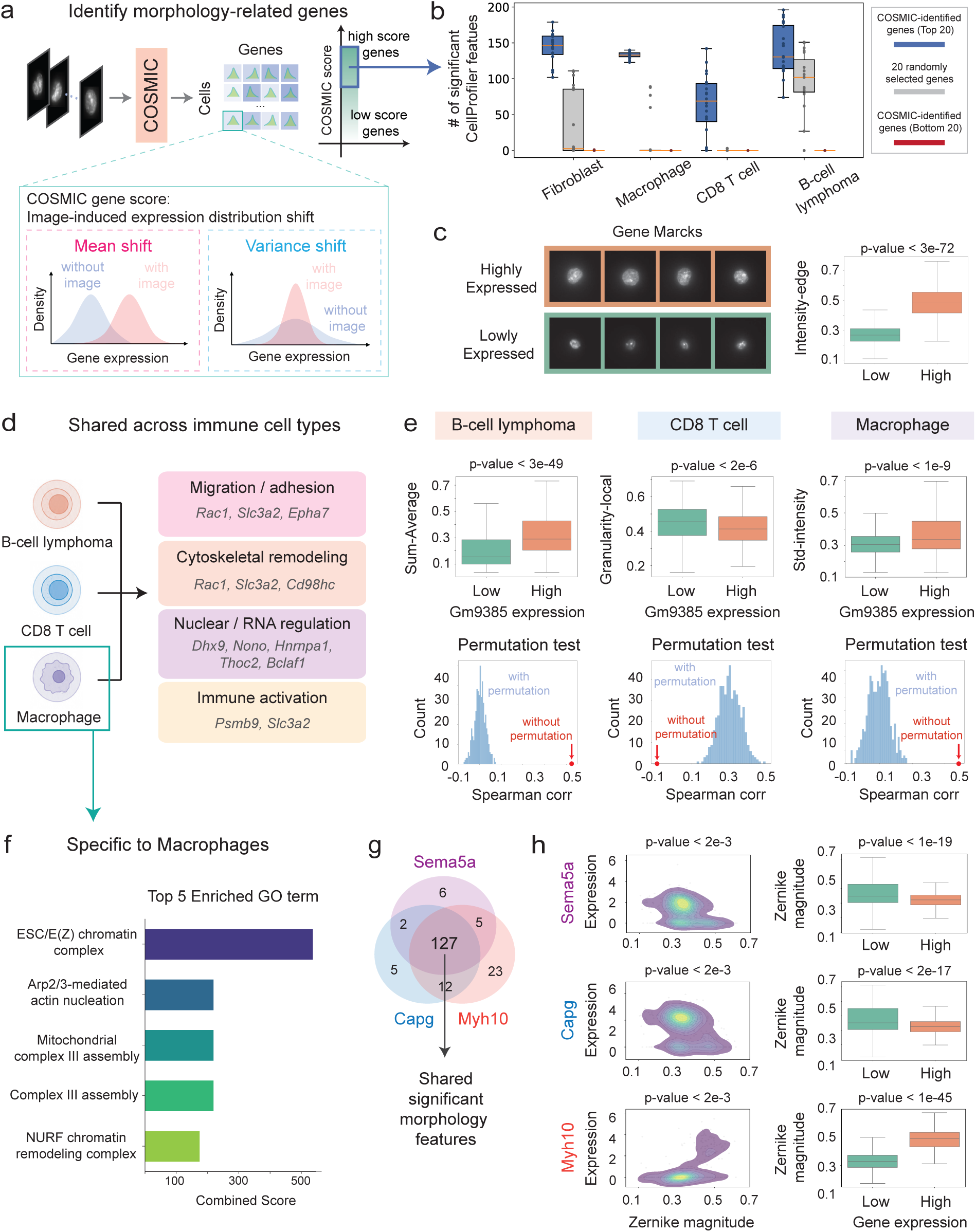
COSMIC proposes shared and cell-type specific morphology-associated genes and gene programs. **(a)** Given paired morphology and transcriptome samples, COSMIC generates gene-expression distributions by sampling with and without image-derived morphology information. Each gene is assigned a morphology-association score based on the resulting image-induced distribution shift, enabling nomination of genes whose expression distribution is associated with nuclear morphology (Methods). The score accounts for both mean shifts and variance or distributional shifts. **(b)** Validation of COSMIC-ranked genes using interpretable nuclear morphology features. Given a gene within a cell type, cells were divided into high– and low-expression groups using the gene’s mean expression. The numbers (out of 271 total) of significantly associated CellProfiler features (two-sided Kolmogorov-Smirnov test; FDR *<* 0.01) are compared between COSMIC’s highest-scoring genes, randomly selected genes, and COSMIC’slowest-scoring genes(*n* = 20). The lowest scoring genesidentified by COSMIC have zero associated CellProfiler features. **(c)** *Marcks*, a top morphology-associated gene nominated by COSMIC in macrophages, is shown as a representative example. (Left) Representative nuclear images illustrate cells with low and high *Marcks* expression. (Right) The corresponding boxplot quantitatively compares a CellProfiler-derived feature integrated edge intensity between the two groups (two-sided Kolmogorov-Smirnov test with Benjamini-Hochberg correction, FDR-adjusted p *<* 3 *×* 10*^−^*^72^). Feature values are normalized to the range 0–1 across the displayed cells. **(d)** Gene Ontology(GO)-based functional annotation of the top 30 morphology-associated genes (FDR-adjusted *p*-value *<* 10*^−^*^20^ for all these genes) shared across B-cell lymphoma cells, CD8 T cells, and macrophages. GO annotations were examined for each gene, and genes with related annotated functions were grouped into broader functional categories, including migration and adhesion, cytoskeletal remodeling, nuclear and RNA regulation, and immune activation. **(e)** COSMIC identifies *Gm9385* as a shared morphology-associated gene across immune cell types. *Gm9385* expression is associated with nuclear morphology in each cell type, and CellProfiler features validate this association. Examples of CellProfiler morphology feature associations, including intensity sum-average, local granularity, and standard deviation of intensity. (Top) Expression-stratified boxplots showing CellProfiler feature shifts between *Gm9385* low– and high-expression cells. (Bottom) The observed Spearman correlation between expression and each feature is compared with a null distribution obtained by permuting gene-expression values across cells; the permutation *p*-value quantifies how frequently a shuffled correlation is at least as extreme as the observed correlation. **(f)** Top five macrophage-specific morphology-associated gene programs identified by COSMIC. **(g)** Comparison of the number of shared significantly associated CellProfiler features between macrophage-specific candidate *Sema5a* with established morphology-associated cytoskeletal genes *Capg* and *Myh10*. **(h)** Association of *Sema5a*, *Capg*, and *Myh10* expression with a shared CellProfiler feature Zernike-magnitude that is significantly associated with all three genes. (Left) Joint distributions of gene expression and Zernike magnitude scores for *Sema5a*, *Capg*, and *Myh10* genes. Changes in the distribution of Zernike magnitude across gene-expression levels indicate statistical dependence between the two variables. (Right) Distribution of Zernike magnitude scores between cells with low and high gene expression of these genes.

To identify genes and gene programs associated with nuclear morphology, we applied COSMIC separately to fibroblasts, macrophages, naive CD8 T cells, and B-cell lymphoma cells in the mouse IRIS dataset. Within each cell type, genes were ranked according to their morphology-association scores. The resulting rankings spanned four categories: well-studied morphology-related genes, genes with morphology associations expected from their known functions, genes with known functions but non-obvious links to morphology, and poorly characterized genes (Supplementary Table 2, 3).

To interpret morphological features and link them to genes identified by COSMIC, we derived nuclear morphology features using CellProfiler (40). We first compared associations of COSMIC-prioritized genes with CellProfiler-derived features against those of randomly selected expressed genes and the lowest-scoring genes identified by COSMIC. Across all four cell types, COSMIC’s top 20 genes were associated with significantly more gene-associated morphology features than randomly selected genes (Fig. 4b; FDR-adjusted *p <* 0.001 across all cell types; one-sided Mann–Whitney U test). Notably, the bottom 20 genes ranked by COSMIC exhibited zero significantly associated morphology features across all evaluated cell types. This independent feature-based analysis supports that the COSMIC rankings capture genes whose expression is associated with measurable nuclear morphology variation. Additional analysis showed that these morphology associations were largely shared across cell types and were dominated by interpretable intensity, shape, and Haralick texture descriptors (Supplementary Fig. 15). Finally, a genome-wide comparison of all human IRIS cells showed that COSMIC recovered every gene flagged by a direct CellProfiler feature-association baseline while nominating many additional morphology-linked genes missed by handcrafted features (Supplementary Fig. 16).

To illustrate how the COSMIC morphology-association score connects to interpretable morphology measurements, we examined *Marcks*, one of the top high-scoring macrophage genes (Fig. 4c). We extracted nuclear features with CellProfiler and compared cells stratified by *Marcks* expression. Cells with high versus low *Marcks* expression showed visually distinct nuclear morphologies and a significant difference (two-sided Kolmogorov-Smirnov test; FDR-adjusted p *<* 3*e −* 72) in CellProfiler-derived integrated edge intensity, indicating that *Marcks* expression separates cells with different nuclear intensity patterns, as well as other features (Supplementary Table 4). This result is also biologically plausible because *Marcks* encodes an F-actin crosslinking protein, and actin cytoskeletal tension is expected to regulate nuclear architecture (50).

We next applied COSMIC to identify morphology-associated genes and gene programs shared across immune cell types. We first identified shared morphology-associated genes across B-cell lymphoma cells, macrophages, and CD8 T cells, and then examined their Gene Ontology (GO) (51) annotations and grouped related functions into broader functional categories. This GO-based functional annotation highlighted migration and adhesion, cytoskeletal remodeling, nuclear and RNA regulation, and immune activation as enriched in COSMIC nominated top-rankes genes (Fig. 4d). These results suggest that the immune-shared genes nominated by COSMIC converge on coherent biological programs that can plausibly influence nuclear morphology or reflect morphology-associated immune-cell states through cytoskeleton–nucleus coupling, nuclear deformation, chromatin remodeling, and immune-state-dependent changes in nuclear mechanics (52; 53; 54; 55). Among the shared candidates, COSMIC also nominated *Gm9385*, a poorly characterized gene with unknown function, as a morphology-associated gene across B-cell lymphoma cells, macrophages, and CD8 T cells. By examining CellProfiler features, we found significant associations (p *<* 1*e −* 10) with different morphology features, including intensity sum-average, standard deviation of intensity, and local granularity with local granularity features shared across all three immune cell types (Fig. 4e, Supplementary Fig. 17). Additional analysis of high-ranking immune-shared genes, including *Cox7a2l*, *Gm10132*, and *Nucks1*, showed that their expression was also significantly associated with nuclear morphology features across immune cell types (Supplementary Fig. 18).

Finally, we focused on the macrophage-specific morphology-associated programs (Fig. 4f). GO enrichment analysis of COSMIC prioritized genes identified terms related to ESC/E(Z) chromatin complex, Arp2/3-mediated actin nucleation, complex III assembly (mitochondrial complex III assembly and respiratory chain complex III assembly), and NURF chromatin remodeling complex. Representative genes driving these enrichments included *Rbbp4*, *Nckap1*, *Bcs1l*, *Uqcc1*, and *Bptf*. These annotations are consistent with prior studies linking *Rbbp4*/RbAp48 to PRC2 (ESC/E(Z))-related chromatin regulation (56), *Nckap1*/Nap1 to WAVE-regulatory-complex control of Arp2/3-dependent actin nucleation and lamellipodial protrusion (57; 58), *Bcs1l* and *Uqcc1* to mitochondrial respiratory-chain complex III assembly (59; 60), and *Bptf* to the NURF chromatin remodeling complex (61). These enriched terms suggest that the macrophage-specific genes proposed by COSMIC converge on coherent biological processes related to chromatin regulation, actin remodeling, and mitochondrial activity. Chromatin regulation and Arp2/3-dependent actin remodeling have established links to nuclear morphology and nuclear mechanics (62; 63). Mitochondrial activity, in contrast, may reflect broader macrophage metabolic and inflammatory states that are indirectly associated with nuclear morphology (64). Together with evidence that macrophage polarization involves nuclear-size, nuclear-stiffness, and chromatin-accessibility remodeling (55), these functions are plausible contributors to macrophage morphology-associated cell states.

Among genes with the highest COSMIC score was *Sema5a* which, unlike canonical cytoskeletal effectors, encodes a semaphorin-family signaling molecule rather than a direct actin-or myosin-based effector. Although semaphorin signaling converges on cytoskeletal remodeling, *Sema5a*’s relationship to nuclear morphology remains undefined (65). We therefore compared its morphology features with two established morphology-related cytoskeletal genes, the actin-capping protein *Capg* and the non-muscle myosin II heavy chain *Myh10* (66; 67), both of which were also highly ranked by COSMIC. For each gene, we stratified macrophages by expression level and identified CellProfiler-derived morphology features that differed significantly between high– and low-expression cells (two-sided Kolmogorov-Smirnov tests with Benjamini-Hochberg correction; FDR *<* 0.05). *Sema5a* shared a large fraction of its significant features with both *Capg* and *Myh10*, including 127 features common to all three genes (Fig. 4g, Supplementary Table 5), indicating that *Sema5a* occupies a macrophage image-feature space shared with known cytoskeletal morphology genes. Mutual-information analysis supported this overlap: for each gene, the density plot captured the dependence between expression and a representative CellProfiler-derived Zernike shape feature, and the expression-stratified boxplot revealed a significant shift in this feature between high– and low-expression cells (Fig. 4h). Notably, higher *Sema5a* and *Capg* expression were associated with lower Zernike magnitude, whereas higher *Myh10* expression was associated with higher values. This pattern is consistent with the opposing actions of these genes on the actin substrate: barbed-end capping and Rac1 down-modulation that limit filament assembly, versus actomyosin contractility that deforms the nucleus (67). The directional effect on Zernike magnitude itself, however, is an observation specific to our data. This analysis shows how COSMIC can be used to nominate genes beyond canonical cytoskeletal effectors and link them to interpretable nuclear image features. Additional top-ranked cell-type-specific genes showed significant associations with distinct CellProfiler-derived nuclear features, supporting that COSMIC identifies interpretable morphology-expression relationships specific to individual cell types (Supplementary Fig. 19).

### COSMIC identifies morphology-associated genes in prostate cancer cells

To evaluate whether COSMIC can uncover biologically meaningful morphology-transcriptome relationships in unstructured and clinically relevant contexts, we next applied it to prostate cancer cells. In contrast to the FUCCI model, where molecular and morphological variation follow a well-defined trajectory, tumor cells exhibit multiple, overlapping axes of heterogeneity, including proliferation, genomic instability, stress responses, and treatment effects. Whether cross-modal mappings remain detectable under such complexity is unclear. DU145 prostate cancer cells offer an ideal system to test this because nuclear morphology is a key histopathological hallmark in prostate cancer, where enlarged, irregular, and hyperchromatic nuclei correspond to high-grade disease and poor clinical outcome (68). DU145 cells, derived from a metastatic lesion, naturally display pronounced variation in nuclear size, shape, and texture that reflects this underlying genomic dysregulation.

We treated DU145 cells with the chemotherapeutic agent Docetaxel, a taxane that blocks mitotic progression, induces G2/M arrest, and ultimately triggers apoptosis (69), and profiled treated and untreated cells using IRIS (Fig. 5a). Embedding the nuclear images using the nuclear encoder revealed two major clusters corresponding primarily to treatment status, with a subpopulation of Docetaxel-treated cells clustering together with untreated cells, which suggests incomplete response or intrinsic resistance. Consistent with the expected phenotype, Docetaxel-responsive cells displayed enlarged, lobulated nuclei with increased eccentricity and reduced solidity, characteristic of G2/M arrest, whereas putative resistant cells likely continued cycling normally and, as a result, retained more regular nuclei (Fig. 5b).

**Figure 5.**
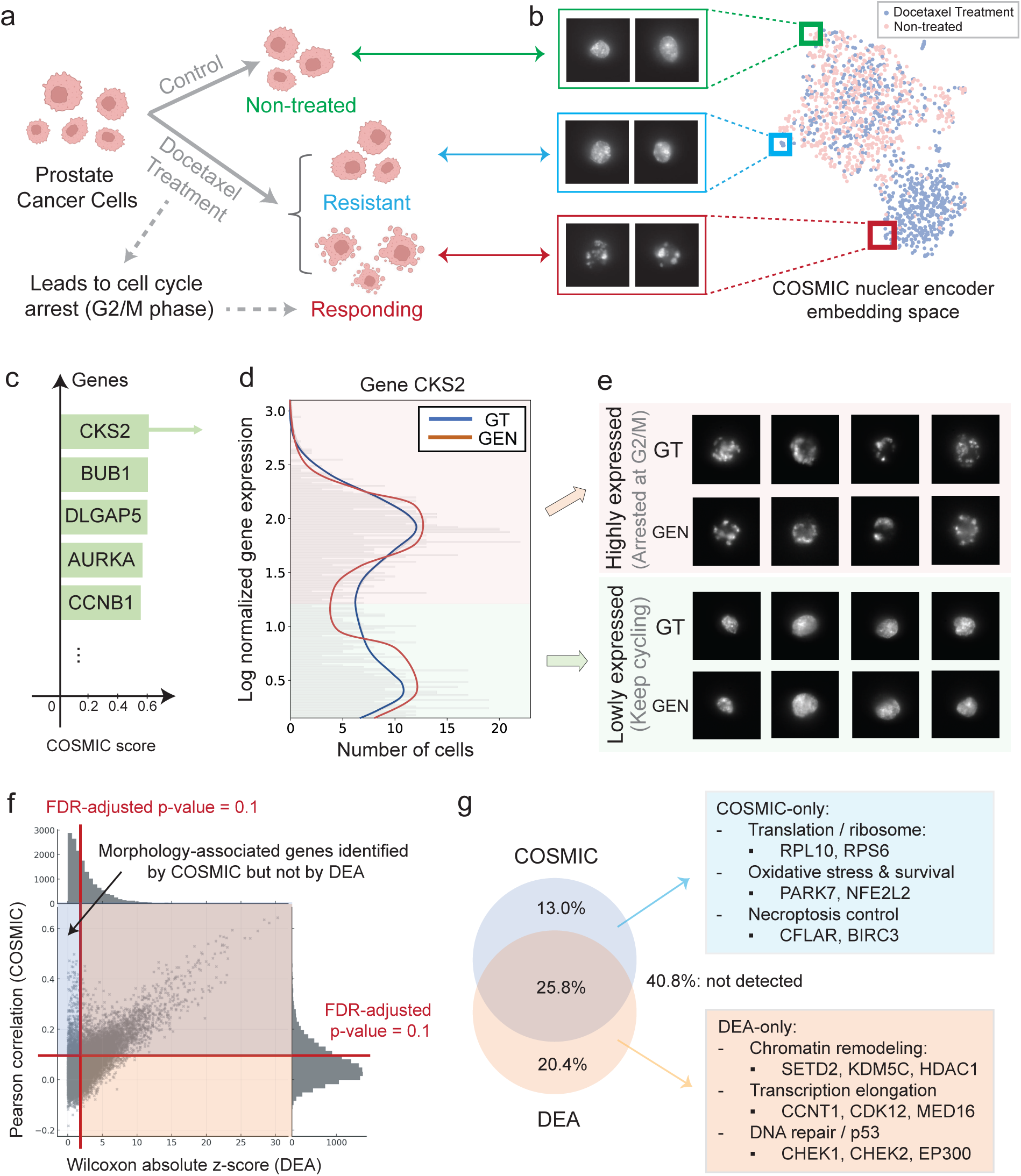
COSMIC identifies morphology-associated genes in prostate cancer cells. **(a)** Illustration of the data generated using IRIS. A subset of the prostate cancer cells (DU145 cell line) was treated with Docetaxel, a chemotherapeutic agent that inhibits cell division during the G2/M-phase, while other cells served as untreated controls. Among the treated cells, only a subset responded to the drug and arrested at the G2/M phase, while the rest remained unaffected. **(b)** UMAP of the COSMIC nuclear encoder embedding space of DU145 nuclear morphology images. The embedding reveals two distinct clusters, with color indicating Docetaxel treatment status (blue: treated, pink: untreated). Randomly selected samples illustrate three conditions: untreated, treated but non-responsive, and treated with a response, consistent with the illustration in panel a. **(c)** COSMIC identifies morphology-associated genes within the DU145 cell line. The top five genes associated with morphology in DU145 cells are *CKS2*, *BUB1*, *DLGAP5*, *AURKA*, and *CCNB1*. **(d)** The histogram of *CKS2* expression shows the number of DU145 cells (x-axis) as a function of their normalized *CKS2* expression level (y-axis). The bimodal distribution reveals two distinct sub-populations: cells with lower *CKS2* expression that continue cycling, and cells with high *CKS2* expression values. COSMIC generated expressions from nuclear morphology (red) agree well with the ground-truth expressions (blue). **(e)** Representative microscopy images for two states: *(i)* cells arrested at the G2/M phase (top), and *(ii)* cells that keep cycling (bottom). Comparison of ground truth nuclei with COSMIC-generated nuclei for both states. **(f)** Comparison of morphology-associated genes identified using COSMIC with genes obtained using differential expression analysis (DEA). A joint distribution plot comparing each gene’s nuclear morphology association (y-axis, Pearson correlation between ground truth and predicted expression, with significance cutoff at FDR-adjusted *p* = 0.1) against its differential expression analysis evidence (x-axis, composite final score representing the maximum score across clusters, with significance cutoff at FDR-adjusted *p* = 0.1). Genes strongly linked to morphology but not differentially expressed genes (upper-left quadrant). **(g)** The proportion of genes unique to COSMIC (blue), unique to differentially expressed genes (DEG) analysis (orange), and shared between the two sets. COSMIC-specific genes are enriched for pathways related to cellular architecture, biomass, and stress responses, whereas DEG-specific genes are enriched for chromatin remodeling, transcription elongation, and DNA damage pathways, which primarily modulate transcriptomic state without necessarily producing immediate changes in nuclear morphology.

COSMIC identified a set of nuclear morphology-associated genes such as *CKS2*, *BUB1*, *DLGAP5*, *AURKA*, and *CCNB1*, which are all well-known regulators of mitosis and drivers of tumor progression (70) (Fig. 5c). Their enrichment for mitotic spindle and chromosomal segregation pathways aligns with the distorted G2/M nuclear shapes observed in Docetaxel-responsive cells, consistent with COSMIC detecting cross-modal structure linked to mitotic instability. For *CKS2*, which was the top morphology-associated gene and a regulator frequently upregulated in aggressive prostate cancers (71), COSMIC recovered two clear expression subpopulations (Fig. 5d). These corresponded to the “responsive” (G2/M-arrested, enlarged nuclei) and “non-responsive” (regular nuclei) Docetaxel groups. One plausible biological interpretation is that *CKS2*-high cells progress rapidly into the vulnerable G2/M window where Docetaxel exerts its effect, while *CKS2*-low cells cycle more slowly and therefore appear less affected at the measured timepoint. Alternatively, the *CKS2*-high cells might represent cells closest to progressing to G2/M in the cell cycle, which would explain that these are the first cells to arrest in G2/M following Docetaxel treatment. We observed similar results for other genes with high correlations (Supplementary Fig. 20).

Additionally, COSMIC’s bidirectional structure allowed us to generate synthetic nuclear images conditioned on transcriptomes. These generated images recapitulated the expected morphological differences. Specifically, *CKS2*-high cells exhibited enlarged, lobulated nuclei, whereas *CKS2*-low cells showed compact, uniform nuclei (Fig. 5e). This demonstrates that COSMIC captures subtle but meaningful cross-modal structure and can resolve molecularly divergent subpopulations even within a homogeneous cancer line.

Notably, morphology-associated genes identified by COSMIC were not simply a subset of differentially expressed genes (DEGs). To compare COSMIC morphology-associated genes identification with standard differential expression analysis (DEA), we identified DEGs between cells arrested in G2/M and cells that continued cycling. We then plotted each gene according to its nuclear morphology association (y-axis; Pearson correlation between ground-truth and COSMIC-predicted expression, FDR-adjusted *p <* 0.1) and its DEA evidence (x-axis; the Wilcoxon absolute z-score, FDR-adjusted *p <* 0.1) (Fig. 5f). The upper-left quadrant of this plot contains genes that are associated with morphology yet not detected as DEGs, indicating that COSMIC recovers morphology-linked genes missed by DEA. Conversely, many DEGs showed little or no association with morphology, which may reflect roles confined to internal nuclear regulation. We speculate that this distinction arises because DEA depends on discrete cell clusters, which can dilute morphology effects, whereas COSMIC links gene expression directly to continuous variation in nuclear architecture.

To characterize the distinction between COSMIC-identified morphology-associated genes and conventional DEGs, we compared gene sets unique to each method (Fig. 5g). COSMIC-specific genes were enriched for pathways governing physical cellular architecture, biomass, and stress responses, including translation and ribosome components (*RPL10, RPS6*) (72; 73), oxidative stress regulators (*PARK7, NFE2L2*) (74), and survival/volume-control genes (*CFLAR, BIRC3*) (75). DEA-specific genes were enriched for chromatin remodeling (*SETD2, KDM5C, HDAC1*), transcription elongation (*CCNT1, CDK12, MED16*), and DNA damage pathways (*CHEK1, CHEK2, EP300*), which are processes that influence transcriptomic variance through enzymatic changes in the nucleus but without necessarily immediately affecting nuclear shape (76).

Together, these results show that COSMIC can robustly recover genes whose expression is tightly coupled to nuclear morphology in Docetaxel-treated prostate cancer cells and that these morphology-associated genes are systematically distinct from conventional DEGs. This highlights COSMIC’s ability to uncover cross-modal structure that links transcriptomic state to continuous variation in nuclear architecture.

### COSMIC identifies global and within-tissue-type morphology-associated genes in zebrafish early embryos

COSMIC is a general framework for linking transcriptome and nuclear morphology and it is also applicable to spatial transcriptomics (ST) data. To test this, we applied COSMIC to the zebrafish embryo development ST dataset (31) to evaluate whether the model can capture morphology-expression relationships in a native tissue context, complementing our analyses on suspension-based IRIS data. IRIS uniquely provides paired high-resolution nuclear images and full transcriptomic profiles from the same single cells, enabling transcriptome-wide quantification of morphology-expression relationships. In contrast, ST provides a tissue-embedded setting in which cells retain their native spatial organization, but the measured gene panel is more limited, and reliable single-cell segmentation remains a challenging problem (77). The zebrafish dataset combines weMERFISH imaging with integrated molecular profiles, providing spatially resolved single-cell morphology, cell-type annotations, and measured expression for 495 genes (Fig. 6a). We generated 48,203 single-cell paired samples by cropping individual cells and their adjacent local regions from the weMERFISH images (64 *×* 64 pixels per image; 0.432 *µ*m/pixel) and linking each cell to its spatially resolved molecular profile and annotated tissue type. These paired samples enabled COSMIC to perform bidirectional modeling between nuclear morphology and gene expression.

**Figure 6.**
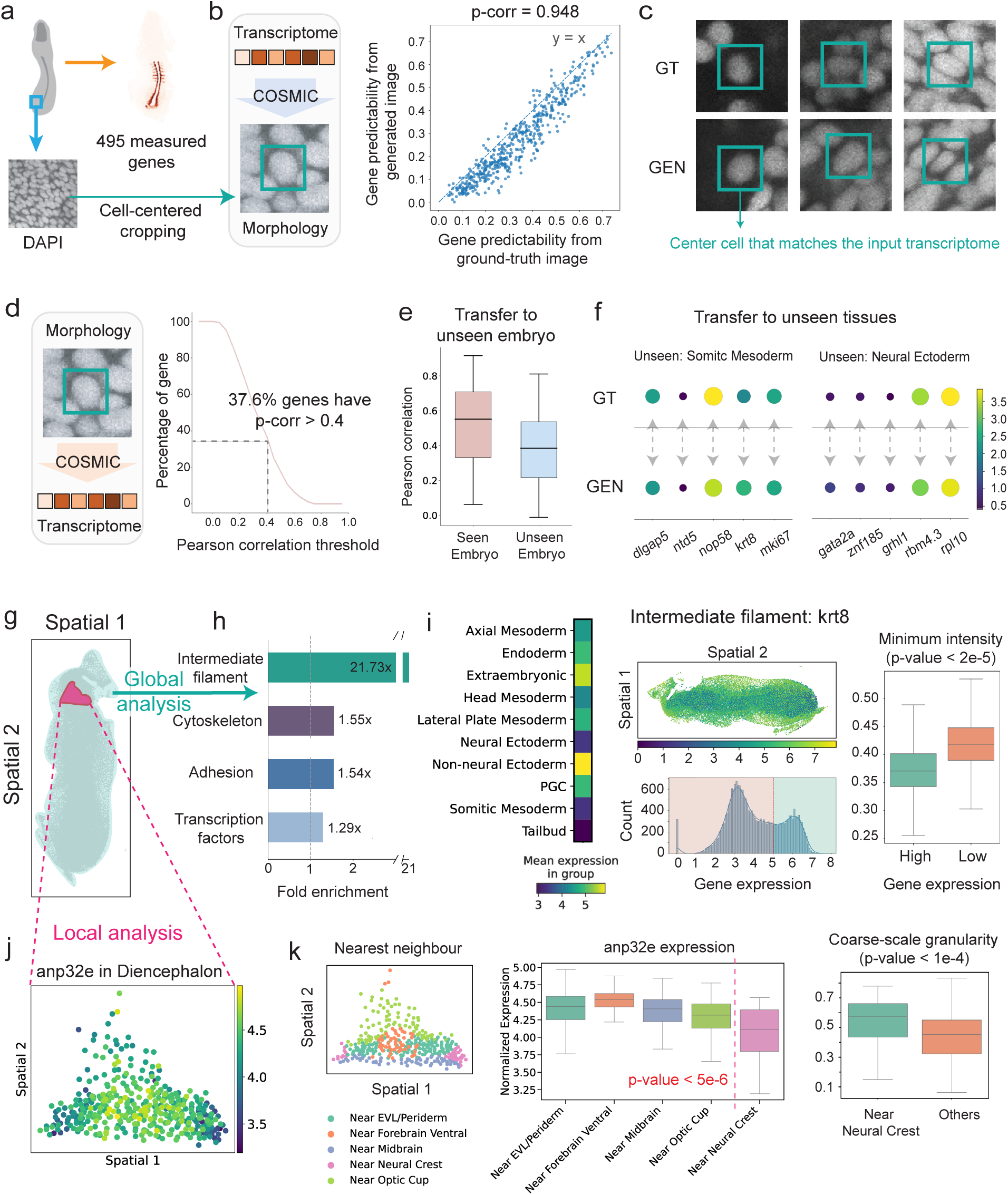
COSMIC identifies global and local morphology-associated genes in the zebrafish weMERFISH dataset. **(a)** Overview of the zebrafish embryo weMERFISH dataset (31). The dataset consists of 495 measured genes paired with DAPI images. We created single-cell paired samples by cropping individual cells and their local regions from DAPI images (64 × 64 pixels per image; 0.432 *µ*m*/*pixel) and linking them to their corresponding transcriptomes. **(b)** Validation of transcriptome-to-image generation. The box highlights the focused cell, which is matched to the input transcriptome. For each gene, morphology-associated predictability scores computed from COSMIC-generated images are compared with scores computed from ground-truth images. The strong agreement (Pearson corr = 0.948) indicates that genes predictable from ground-truth morphology remain predictable from COSMIC-generated morphology. **(c)** Representative matched examples of ground-truth (GT) and COSMIC-generated (GEN) image crops. **(d)** Image-to-expression predictability across genes. The x-axis shows the Pearson correlation threshold for morphology-based gene-expression prediction, and the y-axis shows the percentage of genes with Pearson correlation above that threshold. 37.6% of genes achieve Pearson correlation higher than 0.4. **(e)** Transfer to an unseen embryo. COSMIC is trained on the training samples from one embryo and evaluated on cells from either the test samples of the same embryo or an unseen embryo, testing whether morphology-based gene-expression prediction generalizes across biological replicates. **(f)** Transfer to unseen tissues. In each experiment, cells in one tissue are excluded during training, and COSMIC predicts their average expression profile from morphology at inference. Examples are shown for the best-predicted genes in somitic mesoderm and neural ectoderm, selected based on the highest Pearson correlation between predicted and observed expression across cells. **(g)** Schematic overview of global (*i.e.*, across whole embryo) and local (*i.e.*, tissue-specific) analysis of morphology-associated genes identified by COSMIC. **(h)** Global analysis reveals top enriched Gene Ontology terms among morphology-associated genes (top *n* = 100; BH-FDR adjusted p-value *<* 1 *×* 10*^−^*^30^) across the entire embryo. **(i)** Top-ranked morphology-associated gene example from the global analysis: intermediate filament gene *krt8*. (Left) Average *krt8* expression across different tissues. (Middle top) Spatial distribution of cells colored by *krt8* expression. (Middle bottom) Distribution of *krt8* expression and the threshold used to define high– and low-expression cells. (Right) Comparison of the CellProfiler feature minimum-intensity between the two expression groups; the p-value is from a two-sided Kolmogorov–Smirnov test. **(j)** Local analysis: Within diencephalon cells, COSMIC identifies *anp32e* as a top-ranked morphology associated gene. **(k)** (Left) Visualization of cells annotated as diencephalon cells according to their spatial locations. Cells are colored based on the annotation of their nearest-neighbor cells. (Middle) Distribution of *anp32e* expressions in diencephalon cells grouped by their nearest-neighbor annotation. (Right) Distribution of coarse-scale granularity CellProfiler feature scores between diencephalon cells that are closest to the neural crest and other cell types. P-values are from a two-sided Kolmogorov-Smirnov test.

We first evaluated whether COSMIC-generated images preserved gene-associated morphological information. For each gene, we compared morphology-associated predictability scores, including Pearson correlation and mutual information, computed from ground-truth images with the corresponding scores computed from generated images. These scores were highly concordant, indicating that genes predictable from ground-truth morphology remained predictable from COS-MIC-generated morphology and that COSMIC-generated outputs preserve biologically meaningful gene-level information (Fig. 6b). Representative matched examples further showed that generated image crops preserved local nuclear structures visible in the original image crops (Fig. 6c).

We further applied COSMIC to predict gene expression from nuclear images. Across all measured genes, 37.6% achieved a Pearson correlation greater than 0.4 (Fig. 6d), indicating that nuclear morphology contains sufficient information to recover gene expression variation contained in the gene panels. To test whether the learned morphology-expression relationship generalizes across biological replicates, we trained COSMIC on cells from one embryo and evaluated it on cells from either the same embryo distribution or an unseen embryo. Quantitatively, on the unseen embryo, COSMIC retained approximately 70% of its within-embryo morphology-based gene-expression prediction performance (Fig. 6e). We further evaluated whether COSMIC can generalize to unseen tissues by excluding cells from one tissue during training and then predicting their expression from morphology at inference. COSMIC was predictive across all unseen tissues (Supplementary Fig. 21), and even in the tissue types with lowest performance, somitic mesoderm and neural ectoderm, COSMIC recovered representative average expression profiles for genes such as *dlgap5*, *ntd5*, *nop58*, *krt8*, *mki67*, *gata2a*, *znf185*, *grhl1*, *rbm4.3*, and *rpl10* (Fig. 6f). These results support that COSMIC learns transferable morphology-expression structure that can generalize across embryos and to tissues not seen during training.

We next used COSMIC to nominate morphology-associated genes in the zebrafish embryo at both global (*i.e.*, across entire embryo) and local (*i.e.*, within a tissue) levels (Fig. 6g). At the global level, COSMIC prioritized genes with high morphology-association scores across spatially organized tissue types. Top morphology-associated genes (top *n* = 100; BH-FDR adjusted p-value *<* 1 *×* 10*^−^*^30^) showed enrichment for intermediate filament, cytoskeleton, adhesion, and transcription-factor genes, with intermediate filament genes showing the strongest enrichment (21.73-fold, Fig. 6h). Representative high-scoring structural and tissue-architecture genes included epithelial keratins, such as *krt8*, *krt18a.1*, and *krt4* (78; 79), adhesion and extracellular-matrix genes, including *cdh1*, *fn1a*, *fn1b*, and *nid2a* (80; 81; 82), and the epithelial transcription factor *grhl1* (83; 84; 85). These genes provide positive validation because their expression is expected to represent epithelial structure, adhesion, extracellular matrix organization, and surface tissue architecture. Examining a representative structural example identified by COSMIC, the intermediate filament gene *krt8* (78; 86) revealed that its expression varied across annotated tissue types and that cells stratified by *krt8* expression also showed spatial organization and differences in CellProfiler-derived minimum intensity features (Fig. 6i). These results indicate that *krt8* expression is associated with measurable image-feature variation across the embryo. Other global morphology-associated genes, including structural genes and developmental regulators, showed consistent signals from the analysis (Supplementary Fig. 22).

We then asked whether COSMIC could identify local, within-tissue morphology-expression variation that is affected by the cellular neighborhood. In diencephalon cells (cells located in the posterior lateral diencephalon), COSMIC identified *anp32e* as one of the top morphology-associated genes (Fig. 6j). To interpret this, we used the spatial context available in the ST data and grouped diencephalon cells according to their nearest-neighbor annotation. Diencephalon cells near neural crest cells showed lower *anp32e* expression compared with other diencephalon cells (Fig. 6k; left). This association may reflect shared transcriptional or developmental features between spatially adjacent diencephalon and neural crest cells, consistent with prior work showing that down-regulation of *anp32e* allows H2A.Z to accumulate at Sox-motif promoters and is required for normal neural crest development (87). The same neighborhood-defined groups also differed in image-derived morphology: CellProfiler features revealed a difference in coarse-scale granularity (granularity 15) between diencephalon cells near neural crest cells and other diencephalon cells (Fig. 6k; right). Beyond *anp32e*, COSMIC identified additional within-tissue morphology-associated genes across other annotated tissue types, further supporting its ability to nominate biologically interpretable morphology-expression relationships within spatially organized tissues (Supplementary Note 4, Supplementary Figure 24).

## Discussion

To model and quantify how transcriptional programs relate to nuclear morphology at single-cell resolution, we developed COSMIC, a bidirectional generative framework that quantifies and models the information shared between these two modalities. By training conditional diffusion models on paired transcriptomic and imaging data, COSMIC imposes strong biological priors on the mapping between modalities and enables generation in both directions: synthesizing realistic single-cell nuclear images from gene expression profiles and predicting transcriptomic states from single-cell nuclear images. Through this dual generative capability, COSMIC provides a way to characterize how molecular programs and nuclear morphology are coupled.

We applied COSMIC to datasets generated with IRIS technology, which uniquely provides paired samples of nuclear morphology and full transcriptomic profiles of cells at the single-cell resolution, and spatial transcriptomics data. We focused on the nuclear morphology, and we did not consider other aspects of cellular architecture, including cytoplasmic, organellar, and membrane morphology, but the COSMIC framework is in principle extendable to them. To apply COSMIC, one would need to select or retrain encoders that capture other aspects of cellular architecture.

The overarching aim is to build a detailed link between the two modalities that would enable association of a wide range of gene functions to distinct morphological features at single-cell resolution. At present, the main limitations arise from the scarcity and limited diversity of paired morphology-transcriptome data. Existing paired resources cover a restricted range of tissues and biological contexts, which constrains the spectrum of morphology-transcriptome relationships that the framework can learn. As general-purpose encoders improve and more heterogeneous paired datasets become available, for example through broader deployment of IRIS and related platforms, as well as the continued expansion of ST datasets, this data gap should narrow and COSMIC can be used as a unified generative framework that models the link and supports systematic discovery of morphology-informative genes. An additional challenge is that the relationship between morphology and molecular state is inherently non-unique. Similar nuclear phenotypes may arise from distinct transcriptional programs, while cells with related molecular states may be associated with multiple morphological configurations. COSMIC’s generative formulation provides a natural framework for representing such relationships as conditional distributions rather than deterministic mappings. Quantifying the uncertainty of these distributions will be important for defining when morphology supports reliable molecular inference and when direct molecular measurements remain necessary. Finally, existing data allow us to establish associations between nuclear morphology and the transcriptome, but paired perturbational datasets will be required to model causal relationships.

An open question for multimodal AI models in biology is what kind and scale of data are required to learn reliable cross-modal relationships. In this study, cross-modal relationships were learned from tens of thousands of paired samples, while the nuclear image encoder was pretrained on over 20 million segmented nuclear images. Pretraining on nuclear images substantially improved downstream morphology-to-transcriptome prediction, with performance continuing to improve with more pretraining images but with diminishing gains at larger scales. Cross-modal performance remained largely stable when the number of paired samples was reduced, but deteriorated in the low-data regime. Our results suggest that paired multimodal data are essential for learning the cross-modal mapping, while large-scale nuclear-image pretraining provides transferable visual representations that improve performance. As paired datasets expand to cover more tissues, cell states, and perturbations both larger-scale pretraining and more diverse paired data are expected to bring more benefits for learning cross-modal relationships that generalize across biological contexts.

Altogether, these results demonstrate that generative modeling enabled by paired single-cell measurements can capture and help to quantify the information shared between morphology and gene expression, offering new opportunities to decode the molecular basis of cellular form. Looking ahead, COSMIC sets the stage for a deeper integration of multimodal single-cell data as more paired transcriptomic and imaging datasets accumulate. This will open new avenues for identifying subtle morphological signatures of gene regulation, mapping continuous cellular trajectories, and uncovering cell-state transitions that are currently inaccessible through unimodal analyses.

## Methods

### Overview of COSMIC

COSMIC is a bidirectional generative model that links single-cell transcriptomic profiles to their corresponding nuclear morphologies, enabling prediction and inference in both directions. The architecture comprises two conditional generative modules: a transcriptome-to-image module (*seq2img*) that predicts nuclear morphology from gene-expression profiles, and an image-to-transcriptome module (*img2seq*) that infers transcriptomic profiles from nuclear images. Each module is implemented as a conditional diffusion model trained on paired single-cell transcriptomic and nuclear-morphology measurements generated using the IRIS platform. Conditioning is achieved through embeddings produced by pretrained encoders specific to each modality: a transcriptome encoder for the *seq2img* module and a nuclear-morphology encoder for the *img2seq* module. The training objective follows the standard denoising diffusion loss (37), which can be interpreted as a form of denoising score matching (88), enabling the model to learn biologically grounded mappings between transcriptomic and morphological representations.

We train COSMIC in two stages: *(i)* a forward diffusion process that progressively adds Gaussian noise to microscopy images or transcriptomic profiles, and *(ii)* a denoising process that reconstructs the original data from the corrupted inputs. During training, COSMIC learns to reverse the noise process through iterative refinement, gradually transforming random noise into meaningful nuclear images or gene expression profiles. By conditioning each denoising step on embeddings from the other modality, the model captures cross-modal relationships and learns to generate one modality from the other with high fidelity.

### Nuclear morphology encoder

To obtain robust single-cell nuclear image embeddings for conditioning the generative model, we train a large-scale model on 21,784,309 segmented nuclear images. Specifically, we collected 50,377 whole-well Hoechst-stained microscopy images from JUMP-CP (32), Stojic-lncRNAs (33), and Pascual-Vargas RhoGTPases (34). We segmented individual cells from the whole-well images by applying an adaptive thresholding procedure (Otsu thresholding (89)) followed by connected-component analysis to identify distinct cell regions. We then filtered out nuclei whose projected area lies outside the central range of the area distribution, discarding cells below the 5th percentile or above the 95th percentile in nuclear area, as well as nuclei that are severely occluded or truncated at image borders. After filtering, we crop each detected cell into a 128×128 patch centered on its nucleus. We mask out background pixels and neighboring cells to isolate single-cell morphology.

The model is trained in the encoder-decoder architecture. It uses a vision transformer (ViT-Large; 24 layers) as the backbone and consists of 307M parameters. We train the model in a masked autoencoder (MAE) (35) training fashion. We resize the input images to 224 *×* 224 and divide them into non-overlapping patches. In particular, during training, given an input image **x** from the segmented nuclear images, we apply a random binary mask **m** to produce **x**_masked_ = **m***⊙***x** with a 0.75 masked ratio per image, where *⊙* denotes Hadamard product. The encoder Enc_IMG_ extracts a representation from the masked patches, and the decoder *g_ϕ_* reconstructs the complete image, yielding **x^** = *g_ϕ_*(Enc_IMG_(**x**_masked_)). The training objective minimizes the reconstruction error on the masked regions:

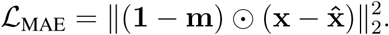

This approach encourages the model to learn biologically meaningful features of nuclear morphology. To leverage these representations for COSMIC, we remove the decoder part and use the learned image embeddings from the last layer of the encoder, **z**_IMG_ = Enc_IMG_(**x**) as conditioning inputs to the transcriptome generation model. The image embeddings are extracted from the full image without applying any masking during inference. We provide further details about training, hyperparameters and modeling choices in the Supplementary Note 3.

### Representations of transcriptomic profiles

To obtain embeddings of transcriptomic profiles for conditioning the generative model, COSMIC is compatible with existing models for batch effect removal (29; 90), as well as single-cell foundation models (30; 36; 91). In particular, we use scVI (29), a probabilistic framework based on variational inference, as a default model for our experiments. In this case, we train the scVI encoder on the transcriptomics profiles of cells. To test the model in a zero-shot fashion, we use the single-cell foundation model UCE (30), in which case we directly embed cells without any training. These embeddings, denoted by **z**_RNA_, are used to condition the diffusion models during generation. Full configuration details for scVI and UCE are given in Supplementary Note 3.

### Diffusion model conditioned on cell representations

Building on the learned representations of nuclear morphology and transcriptomic profiles introduced above, COSMIC employs conditional denoising diffusion probabilistic models (37; 38) to generate one modality from the other by iteratively refining noise-corrupted inputs. In this setting, a clean sample **x**_0_ corresponds to either a nuclear morphology image or a transcriptomic profile of a single cell. COSMIC operates in two phases: *(i)* a forward diffusion process that gradually adds Gaussian noise to a data sample, and *(ii)* a learned reverse process that reconstructs the original data by denoising. Formally, given a clean sample **x**_0_, the forward process generates a sequence **x**_1_, **x**_2_, …, **x***_T_* by adding noise at each timestep:

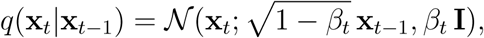

where *β_t_* is a fixed variance schedule. A closed-form expression allows sampling **x***_t_*directly from **x**_0_:

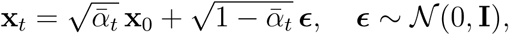

with 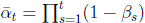.

The reverse process is parameterized by a neural network ***ɛ****_θ_*(**x_t_***, t,* **c**), which predicts the noise component ***ɛ*** given the current sample **x***_t_*, the timestep *t*, and a conditioning embedding **c** derived from the opposite modality. The model is trained to minimize the denoising objective:

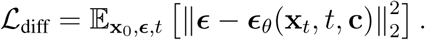

COSMIC includes two such conditional diffusion models. The first is the *seq2img* module with an Attention U-Net (92) architecture designed to synthesize nuclear morphology images from gene expression profiles. Here, the clean sample **x**_0_ is a nuclear morphology image and the conditioning input is the transcriptomic embedding **c** = **z**_RNA_. The U-Net operates on 256 *×* 256 images with multiple resolution levels, and integrates the conditioning embedding at several layers through feature-wise affine transformations and attention. The second is the *img2seq* module with an MLP-based conditional diffusion architecture designed to generate transcriptomic profiles from nuclear morphology images. Here, the clean sample **x**_0_ is a transcriptomic profile, and the conditioning input is the morphology embedding **c** = **z**_IMG_ extracted by the nuclear encoder. To limit the influence of intrinsic gene-expression noise on the diffusion model, a linear head is first applied to map **z**_IMG_ to a baseline prediction **x**_base_. Then, at each step, the diffusion MLP takes as input **x***_t_*, the image embedding **z**_IMG_, the baseline prediction **x**_base_, and the timestep *t*, and is trained to denoise the transcriptomic profile.

By conditioning on **z**_IMG_ or **z**_RNA_, the diffusion models learn to capture biologically meaningful relationships between morphology and gene expression. Through iterative refinement, COSMIC generates high-fidelity outputs that reflect the underlying structure of both modalities. Further details about hyperparameters and model selection are in Supplementary Note 3.

### Identification of morphology-associated genes

To identify genes whose expression is associated with nuclear morphology, we used COSMIC to predict transcriptomic profiles directly from single-cell nuclear images. Let *x_i,g_*denote the observed expression of gene *g* in cell *i*, and let *x*^*_i,g_* denote its expression predicted by COSMIC from the nuclear morphology of the same cell. The intuition is that, if the expression of a gene is associated with nuclear morphology, its morphology-derived prediction should also be associated with its observed expression across cells. We quantified this association using Pearson correlation and hurdle mutual information.

For each gene *g*, we first calculated the Pearson correlation between its predicted and observed expression across cells:

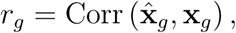

where **x**^*_g_* and **x***_g_* denote the vectors of predicted and observed expression values, respectively.

Part of *r_g_*can arise without any cell-specific image information, so we estimate and remove this baseline using a permutation-derived model. Specifically, we randomly permuted the correspondence between nuclear morphology images and transcriptomic profiles across all cells in our focused cell population (*e.g.*, within fibroblasts). We then trained a separate COSMIC model on the permuted morphology-transcriptome pairs using the model and the same train-test split, and calculated the correlation as:

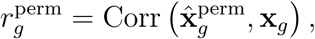

where 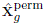 denotes the expression predicted by the model trained on the permuted pairs, therefore quantifying how well gene *g* is recovered without any image-expression correspondence. We used this quantity as a bias correction and defined the adjusted Pearson correlation as:

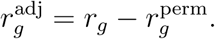

A larger 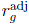 indicates that nuclear morphology captures cell-to-cell variation in the expected expression of the gene.

Pearson correlation primarily captures changes in expected expression and may not detect more general dependencies involving gene-detection probability, expression variance, or other distributional changes. We therefore also calculated the mutual information between the COSMIC-predicted and observed expression of each gene across cells. For notational simplicity, *I*(**a**; **b**) denotes the empirical mutual information estimated from the paired vectors **a** and **b**, and then we estimate the mutual information using

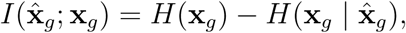

where *H*(**x***_g_*) denotes the entropy of the observed expression distribution across cells and *H*(**x***_g_|* **x**^*_g_*) denotes its conditional entropy given the morphology-derived COSMIC prediction. This quantity measures how much uncertainty about the observed expression of gene *g* is reduced by knowing its morphology-derived COSMIC prediction.

Given that single-cell gene expression is zero-inflated, we used hurdle mutual information (49) to quantify the dependence between the COSMIC-predicted and observed expression of each gene. The estimator separates this dependence into two components: (i) the ability of the prediction to capture whether the observed expression is zero or nonzero, and (ii) the association between predicted and observed gene expression levels among cells with nonzero expression.

To represent the zero-versus-nonzero component, for each cell *i*, we defined the scalar indicator

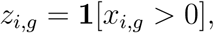

where *z_i,g_* = 1 indicates that gene *g* has nonzero observed expression in cell *i*, and *z_i,g_* = 0 otherwise. Collecting these indicators across the *N* evaluated cells gives the binary vector:

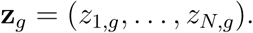

To represent the nonzero component, we first defined:

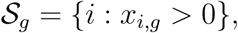

and then we defined:

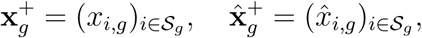

as the collection of nonzero observed expression values of gene *g* across cells. We also define:

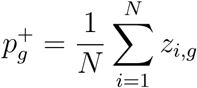

as the fraction of cells with nonzero observed expression. The hurdle mutual information was then estimated as:

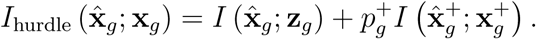

The first term measures whether COSMIC recovers the observed zero-versus-nonzero expression pattern across cells. The second term measures the association between predicted and observed expression levels among cells with nonzero expression, weighted by the fraction of such samples.

To get the top morphology-associated genes, we rank the genes using calibrated Pearson correlation and hurdle mutual information. Specifically, we first ranked genes separately in descending order of Pearson correlation and hurdle mutual information. Let 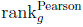 and 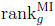 denote the corresponding ranks, with smaller values indicating stronger morphology association. The final COSMIC rank was defined as:

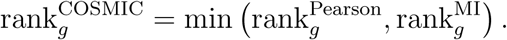

Thus, a gene receives a top COSMIC rank when either Pearson correlation identifies a shift in expected expression or hurdle mutual information identifies a broader change in its expression distribution.

To identify morphology-associated genes shared across multiple cell types, we first calculated 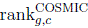 independently within each cell type *c*. For a set of cell types *C*, we then defined the shared rank as:

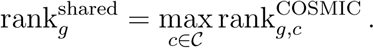

The shared rank is therefore determined by the cell type in which the gene has its lowest ranking. Consequently, a gene receives a top shared ranking only when it is top-ranked in every cell type under comparison. For the immune-shared analysis, this procedure was applied to the B-cell lymphoma cells, macrophages, and CD8 T cells, and we selected the top 30 shared genes.

While relative ranking identifies the top nuclear morphology-associated genes compared to the rest of the genes, it does not guarantee that these associations are statistically meaningful. To ensure biological relevance, we computed *p*-values for both metrics and controlled the false discovery rate (FDR) across all tested genes using the Benjamini-Hochberg procedure. For the Pearson correlation, significance was assessed using a standard two-sided *t*-test. For the mutual information, calculating exact *p*-values for the continuous expression levels is mathematically intractable. Therefore, we assessed its statistical significance based purely on the model’s ability to predict whether a gene was turned on or off (zero versus nonzero expression), using a likelihood-ratio test. A gene’s association with nuclear morphology was considered statistically significant if its adjusted *p*-value (FDR) for the corresponding metric was below 0.05. In our pipeline, any gene selected for downstream analysis must meet this strict significance threshold. The notation FDR-adjusted *p*-value *< α* indicates that the adjusted *p*-values for both the Pearson correlation and the hurdle mutual information are strictly less than *α*.

### Datasets and data pre-processing

For COSMIC training, we use paired single-cell nuclear images and transcriptomic profiles obtained from the IRIS platform, which enables simultaneous capture of scRNA-seq data and fluorescence microscopy images from the same individual cells. IRIS performs deterministic single-cell imaging before droplet encapsulation, using machine-vision-guided positioning of individual cells in a stopping zone. The platform acquires high-resolution images at approximately 500 nm lateral resolution using a 40*×* NA 0.9 objective. For each focal plane, the microscope acquires one brightfield channel and four epifluorescence channels, such as DAPI, FITC, TRITC, and Cy5 or similar channels. Focal planes are acquired as z-stacks across the channel height using a piezoelectric stage, enabling optimal focus selection and visualization of intracellular structures.

The paired COSMIC dataset includes both mouse and human samples. The mouse dataset contains 9,039 cells from 3T3, RAW, CAR A20, and naive CD8^+^ T cells, spanning 8 batches. The human dataset consists of 17,109 cells from DU145, RPE1, C4-2B, lymphocytes, and monocytes, spanning 14 batches. In the main mouse experiments, we use 4,520 cells for training and 4,519 cells for testing. In the human experiments, we use 8,555 cells for training and 8,554 cells for testing. For PBMC experiments, buffy coats and whole blood were obtained from fully anonymized healthy donors provided by ‘Transfusion Interregionale’ (Biopole, 1066 Epalinges, Project-Nr. P 285).

For image preprocessing, raw IRIS brightfield images of the stopping zone have a size of 512 *×* 2048 pixels. A YOLO-based object detection model is used to identify each cell’s location and optimal focal plane. Centered 250 *×* 250 pixel crops are then generated for each fluorescent channel. A DeepLabV3 segmentation model is subsequently used to compute cell masks for quality control and feature extraction. For COSMIC, the final nuclear image inputs are cropped around segmented nuclei, resized or represented as 256*×*256 pixel images, and intensity-normalized to the [0, 1] range. These final preprocessed nuclear morphology images serve as input to the COSMIC nuclear encoder and as targets for the sequence to image diffusion model.

We process transcriptomic data using standard single-cell RNA-seq workflows. Raw count matrices are filtered to remove low-quality cells, including cells with fewer than 500 detected genes or high mitochondrial gene content, and genes expressed in fewer than 5 cells. Counts are normalized using total-count scaling so that each cell has a total count equal to the median count across cells, followed by a log1p transformation.

The nuclear image pretrained masked autoencoder was trained separately from the paired IRIS data. For this pretraining, we construct a large-scale single-cell morphology dataset by aggregating Hoechst-stained nuclear images from three publicly available sources: the JUMP-Cell Painting dataset (32) with U2OS cells, the stojic-lncrnas dataset (33) with HeLa cells, and the pascualvargas-rhogtpases dataset (34) with LM2 cells. In total, this aggregation yielded approximately 50,377 whole-well microscopy images and 21.8 million segmented single-nucleus images. From each whole-well image, nuclei are automatically segmented using a deep learning-based pipeline, after which cropped image patches are extracted around each nucleus using a fixed-size bounding box. Quality control removes images with overlapping nuclei, debris, or segmentation errors, retaining only well-centered and morphologically intact nuclei. These curated image patches are used to train the COSMIC nuclear encoder from scratch with a masked autoencoder objective, enabling the model to learn rich and generalizable representations of nuclear morphology.

## Acknowledgements

We are grateful to Duygu Koldere Vilain for making the professional illustration of the IRIS technique. We are also grateful to Natasha Samson for the data on the cell line CAR A20. M.B. gratefully acknowledges the support of the Swiss National Science Foundation (SNSF) starting grant TMSGI2 226252/1, SNSF grant IC00I0 231922, SNSF grant 10.004.411, Swiss Cancer League (KFS-6340-02-2025-R), Kamprad Foundation (Cancer Immunotherapy Networking Program) and the Swiss AI Initiative (ab006, ab023). M.B. is a CIFAR Fellow in the Multiscale Human Program. We gratefully acknowledge the support of the Peter und Traudl Engelhorn Foundation to R.T. and the Swiss AI Fellowship to S.W. Figure elements, including icons of species, were created with BioRender.com.

## Author Contributions Statement

S.W., R.T., J.B., B.D. and M.B. designed the study. S.W., R.T. and M.B. contributed new analytical tools and designed the algorithm. S.W. performed the experiments and developed the software. S.W., R.T., J.B., C.L., J.P., Y.W., W.K., B.D., and M.B. contributed to data interpretation and analysis. J.B., C.L., N.G., T.F., E.B., J.P., J.L., B.D., and W.K. contributed to generating IRIS data. S.W., R.T., B.D., and M.B. wrote the manuscript with the input from J.B., C.L., Y.W. and W.K. Y.W contributed the raw spatial transcriptome images of nuclear morphology and helped with data interpretation. B.D. and M.B. supervised the IRIS data analysis research. M.B. supervised the research.

## Data availability

The JUMP-CP dataset is available at https://github.com/broadinstitute/cellpainting-gallery; the stojic-lncrnas dataset is available at https://idr.openmicroscopy.org/webclient/?show=screen-2301; the pascualvargas-rhogtpases is available at https://idr.openmicroscopy.org/webclient/?show=screen-1651. The IRIS datasets will be publicly available upon publication.

## Code availability

COSMIC was written in Python 3.9 using the PyTorch library. The source code is available on GitHub at https://github.com/mlbio-epfl/COSMIC.

