## Supplementary for "Generative modeling reveals the connection between nuclear morphology and gene expression"

[*]Shared second author.

[**]Shared senior author.

This PDF file includes:

Supplementary Notes 1 to 5
Supplementary Figures 1 to 23
Supplementary Table 1 to 5
Supplementary References

#### Supplementary Note 1 Datasets and preprocessing

**Human PBMC and T cell enrichment** Peripheral blood (PB) was diluted 1:1 with DPBS (Thermo Fisher Scientific, #14190169). PB:DPBS was over-layered on Ficoll-PaqueTM PLUS (cytiva, #10351856) enabling purification of PBMCs by density centrifugation at 800g for 30 min. The interphase containing PBMCs was washed once with DPBS and cell count was obtained with Neubauer (FAUST, #9161078) counting chamber with Trypan blue live–dead stain (Thermo Fisher Scientific, #T10283). When required, human CD8+ T cells were enriched from PBMCs using a column-based magnetic-activated cell sorting (MACS) Pan CD8 T cell Isolation negative selection kit (Miltenyi Biotec, #130-096-495) or for naive CD8+ T cells the Naive Pan T Cell negative selection isolation Kit (Miltenyi Biotec, #130-097-095) followed by CD8+ T Cell negative selection isolation kit (Miltenyi Biotec, #130-096-495), according to the manufacturer's specifications. For PBMC experiments performed by the laboratory of Prof. Bart Deplancke, buffy coats and whole blood were obtained from fully anonymized healthy donors provided by 'Transfusion Interregionale' (Biopole, 1066 Epalinges, Project-Nr. P_285). We added this in the revised manuscript (Datasets and data pre-processing).

**Murine T cell enrichment** Peripheral blood (PB) was diluted 1:1 with DPBS (Thermo Fisher Scientific, #14190169). PB:DPBS was over-layered on Ficoll-PaqueTM PLUS (cytiva, #10351856) enabling purification of PBMCs by density centrifugation at 800g for 30 min. The interphase containing PBMCs was washed once with DPBS and cell count was obtained with Neubauer (FAUST, #9161078) counting chamber with Trypan blue live–dead stain (Thermo Fisher Scientific, #T10283). Murine naive CD8+ T cells were enriched from PBMCs using STEMCELL EasySep™ Mouse Naïve CD8+ T Cell Isolation Kit (STEMCELL, #19858), according to the manufacturer's specifications.

**Cell lines** Human HEK 293T cells (American Type Culture Collection (ATCC), #SD-3515), mouse NIH/3T3 (ATCC, #CRL-1658), Fluorescent ubiquitination-based cell cycle indicator Fucci(SA)5 mouse 3T3 cells (provided by the D. Suter laboratory, EPFL Lausanne) and human RPE-1 cells (provided by the F. Naef laboratory, EPFL) were used. Drosophila S2 cells (provided by the B. Lemaitre laboratory, EPFL) were used. DU-145 and C4-2B (provided by the W. Karthaus laboratory, EPFL Lausanne) were used. Rela-GFP RAW 264.7 were used [1].

**Cell culture** Fucci(SA)5 3T3 cells were cultured in DMEM (Thermo Fisher Scientific, #41966029) with 10% v/v FBS (Thermo Fisher Scientific, #A5670701) and 1% v/v and penicillin–streptomycin (Thermo Fisher Scientific, #15140122). RPE-1 cells were cultured in DMEM GlutamaxTM (Thermo Fisher Scientific, #10566016) with 10% v/v FBS, 1x MEM Non-essential amino acids solution (Thermo Fisher Scientific, #11140050) and 1% v/v and penicillin–streptomycin. C4-2B were cultured in RPMI GlutamaxTM (Thermo Fisher Scientific, #61870036) with 10% v/v FBS and 1% v/v and penicillin–streptomycin. DU145 were cultured in DMEM GlutamaxTM (Thermo Fisher Scientific, #10566016) with 10% v/v FBS and 1% v/v and penicillin–streptomycin. Where stated DU145 cells were treated with 1 nM Docetaxel (Selleckchem, #S1148). S2 were cultured in Schneider's Drosophila Medium (Thermo Fisher Scientific, #21720024) with 10% v/v FBS and 1% v/v and penicillin–streptomycin. Rela-GFP RAW 264.7 were cultured in RPMI1640 (Thermo Fisher Scientific, #11875093) with 10% v/v FBS (Thermo Fisher Scientific, #A5670701), 1% v/v penicillin–streptomycin (Thermo Fisher Scientific, #15140122) and 2mM L-Glutamin (Thermo Fisher Scientific, #25030081).

Cells were cultured to 70% confluency. Prior to use, cells were washed with DPBS, dissociated with 0.25% Trypsin-EDTA (Thermo Fisher Scientific, #25200056), washed with DPBS and counted with Neubauer counting chamber (FAUST, #9161078) with Trypan blue live-dead stain (Thermo Fisher Scientific, #T10283). Preparation of murine anti-CD19 CAR-T cells Transduction agent was prepared with anti-CD19 CAR carrying plasmids (Addgene, #192115) and pCL-Eco packaging plasmid in Phoenix-ECO cells (ATCC, #CRL-3214) (Addgene, #12371). Virus-containing supernatants were collected at 48h post-transfection, passed through a 45 m filter, and kept at 4°C overnight. Virus-containing supernatants were centrifuged without cells at 2,000g for 2h at 32°C on plates protamine sulfate coated plates (10 ng/ml in PBS overnight, Thermo Fisher Scientific, #10341041). Cells were added (2 million T cells per ml of medium) and centrifuged for 30 min at 300g at 32°C. On day post transduction, the supernatant was replaced with medium supplemented with mouse IL-2 (10 ng/ml, Biolegend, #575408) and IL-7 (10 ng/ml, Biolegend, #577808). Cells were passaged every 24h at a ratio of 1:2 for two days prior to in vitro co-culture. CAR T cells were co-cultured with A20 (ATCC, #TIP-208), expressing CD19, at a ratio of 1:1 with a total cell number 200,000 cells per 96 well round bottom well for four hour.

**Live-cell organelle staining** Live cells were stained for 30 min at 37°C and 5% CO2. Nuclear stains were performed with 5 ng/ml Hoechst 34580 (Merck, #63493) in DPBS.

#### Supplementary Note 2 Related work and baselines

Existing methods have inferred transcriptomic profiles from histology images. PC-CHiP [2] associated CNN-derived image features with bulk RNA-seq profiles, while HE2RNA [3] used pre-trained ResNet-50 features to predict gene expression from H&E whole-slide images. More recent transformer-based methods, including tRNAsformer [4] and SEQUOIA [5], combine transformers with multiple-instance learning. However, these approaches operate at the bulk-tissue level and lack single-cell resolution.

MorphNet [6] and Multi-domain Translation (MDT) [7] model transcriptomic–morphological relationships at the cellular level. MorphNet uses adversarial training to generate microscopy images from gene-expression vectors, but is susceptible to mode collapse and often produces blurry images with limited morphological detail. MDT instead aligns unpaired transcriptomic and morphological data in a shared representation space. Although this removes the need for paired measurements, the absence of cell-level correspondences and explicit biological constraints limits its ability to recover precise, fine-grained, and cell-type-specific relationships.

All baselines were trained on the IRIS dataset (17,109 cells, 17,982 genes) so that every method sees the same cells, the same nuclear crops and the same transcriptome preprocessing. Raw counts were normalized per cell to a common library size and $\log(1+x)$-transformed. Nuclear crops were loaded as single-channel images scaled to $[0, 1]$ and resized to the input resolution required by each model. Because published implementations of MorphNet and MDT target different input modalities and dataset sizes, we kept their architectures and training objectives as described by the original authors and adapted only the input and output dimensions to IRIS; all deviations are stated explicitly below.

**MorphNet** MorphNet was trained as a conditional generative model mapping transcriptomic state to nuclear morphology. Rather than conditioning on the full $17{,}982$-dimensional expression vector, which is too high-dimensional to serve directly as a StyleGAN conditioning input, we conditioned the generator on a 64-dimensional transcriptome embedding computed for each cell, giving a conditioning matrix of size $17{,}109 \times 64$. Cells were paired with their own nuclear image, and the model was trained on the odd-indexed half of the dataset, with the held-out half reserved for evaluation.

The generator follows the StyleGAN2 design: an 8-layer mapping network with equalised-learning-rate linear layers maps the concatenation of a 256-dimensional noise vector $z$ and the 64-dimensional conditioning vector to a 512-dimensional style vector $w$, with $z$ pixel-normalised before concatenation. The synthesis network starts from a learned $4 \times 4$ constant and applies pairs of weight-modulated/demodulated $3\times3$ convolutions with per-layer noise injection, doubling resolution at each stage up to $256\times256$. Each stage carries a $1\times1$ modulated output head producing a single-channel image; these are combined with the skip-connection scheme of StyleGAN2. The discriminator is a mirrored residual stack of equalised $3 \times 3$ convolutions with $2 \times 2$ average pooling, and conditions on the continuous vector through a projection term: the pooled feature map is combined with a linear embedding of the conditioning vector by inner product and added to the unconditional logit. The channel schedule uses a base width of 8192 halved per stage with a floor of 16 channels.

Both networks were trained with the non-saturating logistic loss, without $R_1$ or path-length regularisation and without style mixing. We used Adam with a learning rate of $2 \times 10^{-4}$ and $\beta = (0.0, 0.99)$ for both generator and discriminator, a micro-batch of 4 with 4 gradient-accumulation steps (effective batch size 16), mixed-precision training and gradient checkpointing, for 100 epochs. Images for evaluation were generated by sampling a fresh $z$ per cell and conditioning on that cell's held-out transcriptome embedding.

**Multi-domain translation (MDT)** MDT aligns the two modalities in a shared latent space without requiring per-cell correspondence, and we implemented it in exactly that regime.

The image branch is a convolutional VAE. Nuclear crops were resized to $256 \times 256$ and replicated across three channels. The encoder applies four $4 \times 4$ stride-2 convolutions ($3 \rightarrow 64 \rightarrow 128 \rightarrow 256 \rightarrow 512$ channels) with ReLU activations, followed by linear layers producing the mean and log-variance of a 64-dimensional latent; the decoder mirrors this with four transposed convolutions and a sigmoid output. It was trained with the standard evidence lower bound (summed binary cross-entropy reconstruction term plus KL divergence) using Adam with a learning rate of $10^{-3}$ and batch size 256. After training, we extracted the 64-dimensional latent for every nucleus and held these fixed.

The transcriptome branch is a fully connected autoencoder operating on the 2,000 most highly variable genes, selected with `scanpy` using the experiment identifier as batch key. The encoder is $2000 \rightarrow 256 \rightarrow 256 \rightarrow 256 \rightarrow 64$ with LeakyReLU $(0.2)$ activations, and the decoder mirrors it back to 2,000 dimensions.

Alignment between the two 64-dimensional latent spaces is enforced adversarially. A discriminator ($64 \rightarrow 64 \rightarrow 16 \rightarrow 1$, LeakyReLU $(0.2)$, sigmoid output) is trained with binary cross-entropy to separate image latents from transcriptome latents, while the transcriptome encoder is trained to fool it. The discriminator therefore only ever compares the two *marginal* latent distributions and never observes a true cell-to-cell correspondence, which is precisely the unpaired setting MDT is designed for. The full objective is the mean squared reconstruction error of the transcriptome autoencoder plus the adversarial term weighted by $10$. Both the autoencoder and the discriminator were optimized with Adam (learning rates $10^{-4}$ and $10^{-3}$ respectively) at batch size 256 for 10 epochs.

To predict expression from an image at test time, the image VAE latent of a held-out nucleus is passed through the transcriptome decoder. Evaluation uses the true image–transcriptome correspondence, so that MDT is scored on exactly the same paired test cells as every other method; only its training is unpaired.

**Regression on CellProfiler nuclear morphology features** To test whether classical morphology descriptors suffice, we extracted CellProfiler features and regressed expression on them directly. Each nuclear crop was converted to grayscale, min–max rescaled to $[0, 1]$, and non-finite values set to zero. We then applied the core measurement set of the `cp_measure` implementation of CellProfiler (`get_core_measurements`), which yields **271** numeric descriptors per nucleus spanning size and shape, intensity, texture, granularity and zernike moments. Features were matched to transcriptomes by cell identifier.

Features were standardised using training-split means and standard deviations, with non-finite entries imputed by the training mean; expression targets were left on the $\log(1 + x)$ scale. The regressor is a multilayer perceptron with two hidden layers, ReLU activations and dropout $0.1$, and a softplus output to enforce non-negative predicted expression, trained with mean squared error using AdamW (learning rate $10^{-3}$, weight decay $10^{-4}$) at batch size 128 for 10 epochs.

Because a small number of interpretable descriptors might behave differently from the full set, we did not fix the feature count a priori. Features were first ranked into four semantic priority groups with size and shape, intensity and brightness, sharpness and texture, and all remaining features, and we then swept the number of retained features over $k = 271$, training an independent model for each $k$. Within each run, the epoch with the highest mean per-gene Pearson correlation on the validation split was selected, and that checkpoint alone was applied to the test split.

**Unpaired COSMIC** To isolate the contribution of paired supervision from that of the learned image representation, we retrained COSMIC with its architecture and optimizer held exactly fixed and only the correspondence destroyed. Before training, the vector of cell identifiers was permuted once globally, so that each transcriptome was associated with the nuclear image of a different, randomly chosen cell; the permutation was fixed for the whole run rather than resampled per epoch, so the model sees a consistent but incorrect pairing. The marginal distributions of both modalities, the number of training examples and the class balance are therefore identical to the paired run, and the only quantity removed is the cell-level correspondence.

As in the paired model, images were resized to $224 \times 224$ and replicated to three channels, and the backbone is a ViT-L/16 encoder initialised from masked autoencoder pretraining, with all blocks frozen except the final transformer block, followed by a linear head onto the gene dimension. Training used mean squared error with Adam at a learning rate of $10^{-4}$ and batch size 32 for 5 epochs. Evaluation was carried out on the true, unpermuted image–transcriptome pairs, so that the unpaired model is scored on the same test correspondence as the paired model.

#### Supplementary Note 3 Hyperparameters and Model Selection

**Encoder of Gene Expression.** In our transcriptomic encoder, we preprocess the dataset (mouse and human separately) by applying library-size normalization, log transformation, and selecting the top 1024 highly variable genes using `experiment_ID` as the batch key. The model is instantiated as an SCVI [8] variational autoencoder with a hidden dimension of 128 and a latent dimension of 64. Training is conducted for a maximum of 50 epochs while accounting for batch-specific effects through the experiment ID. The model is trained for a fixed budget of 50 epochs without early stopping or checkpoint-based selection. The final trained model is adopted directly, and its latent representation is used as the condition of the conditional diffusion model. When applying UCE [9] as the transcriptomic encoder, we directly use the pretrained 4-layer UCE model with its default configuration, without any further fine-tuning or retraining. For UCE, we perform inference only and use the resulting latent representation as an the conditioning signal for the diffusion model.

**Encoder of Cell Morphology.** In COSMIC, we pretrain the image encoder as a masked autoencoder built on a ViT-Large backbone with $16 \times 16$ patches and $224 \times 224$ inputs. We mask a fixed ratio of 0.75 tokens per image and reconstruct the corresponding pixel patches using an MAE decoder configured with a decoder embedding dimension of 512, a single transformer layer, 16 attention heads, an MLP ratio of 4.0, and no projection or attention dropout. Images are resized to 224 and augmented with independent horizontal and vertical flips (probability 0.5 each). Training uses mean-squared error between predicted and target masked patches, the AdamW optimizer with a learning rate of $1.5 \times 10^{-4}$, and mini-batches of size 256. We train the MAE for up to 10 epochs without early stopping or validation-based selection. We adopt the final-epoch checkpoint for downstream use.

**Decoder of Gene Expression.** In COSMIC, we map image-derived embeddings to gene-expression vectors in two training stages. Targets are constructed from the mouse dataset by applying library-size normalization followed by a $\log(1{+}x)$ transform. In Stage 1, a ViT-L/16 masked autoencoder backbone encodes each image into a 1024-dimensional feature vector (with only the last transformer block trainable), and a linear regressor maps this feature to the full gene-expression vector; we train this baseline with a mean squared error loss using Adam (learning rate $1 \times 10^{-4}$), a batch size of 32 for training (and 256 for testing), for 30 epochs. In Stage 2, we freeze the encoder and regressor and train a residual diffusion model on the prediction residuals $r_0 = x_0 - \hat{x}_{\text{base}}$, using a denoiser that is a 3-layer MLP (2048 of hidden dimension) conditioned on the 1024-dimensional image feature, We use a short diffusion horizon of $T = 10$ steps with a linear noise schedule from $5 \times 10^{-5}$ to $5 \times 10^{-4}$ and train the residual model with an MSE loss for $10$ epochs. We use a fixed train/test split implemented by alternating indices (odd for training, even for testing), evaluate on the test split after each epoch, and adopt the final checkpoint from Stage 2 for downstream use.

**Decoder of Cell Morphology.** In COSMIC, we train a conditional diffusion image decoder using the Imagen framework to generate $256 \times 256$ images from transcriptomic embeddings. The architecture comprises a single UNET with base dimension $32$, conditioning dimension $64$, and multi-scale widths given by $(1, 2, 4, 8)$. Each scale uses one ResNet block; self- and cross-attention are enabled only at the final scale. Conditioning vectors are $64$-dimensional with classifier-free guidance via a drop probability of $0.1$. The diffusion process spans $1000$ timesteps with prediction objective $x_{\text{start}}$. Inputs are resized to $256$ without additional augmentations. Training optimizes the Imagen loss with AdamW at a learning rate of $1.5 \times 10^{-4}$ and a mini-batch size of $16$. We train for up to $100$ epochs (iterating over mini-batches) without early stopping or validation-based checkpointing. We adopt the most recent checkpoint for downstream experiments.

#### Supplementary Note 4 Additional analysis in weMERFISH

Beyond *anp32e*, COSMIC nominated many genes whose expression was predictable from morphology within a single annotated tissue type. We therefore performed the same post hoc spatial analysis for additional genes. Specifically, for cells from each annotated tissue, we grouped cells by the majority identity of their 100 nearest spatial neighbors, restricted to tissues adjacent to the target tissue, and then tested whether these neighborhood-defined groups differed in both gene expression and DAPI-derived nuclear morphology. To reduce the possibility that these signals reflected annotation error, we also checked each cell's transcriptome-wide identity and required the neighborhood-distinguished cells to retain the identity of their annotated tissue rather than that of the neighboring tissue.

In every case, neighborhood identity was not provided as an input to COSMIC. Instead, neighborhood serves as an unmodeled biological correlate that helps interpret the morphology-expression relationships discovered by COSMIC. Together with anp32e, these examples support that COSMIC can nominate biologically interpretable morphology-associated genes even within a single annotated tissue type.

**Gene *msx1b* within trigeminal placode.** COSMIC identified *msx1b* as a gene whose expression was predictable from morphology within annotated trigeminal placode cells (Supplementary Fig.24a). Grouping these cells by their nearest-neighbor tissue type, trigeminal placode cells near Neural Crest cells had markedly higher *msx1b* expression than other Trigeminal placode cells (mean normalized expression 3.50 vs. 1.74; $p < 3 \times 10^{-21}$). This is consistent with prior developmental studies showing that Msx genes act at the neural plate border, where cranial placodes and neural crest arise. In particular, Bmp-dependent regulation of Msx expression specifies Msx activity in the neural fold region, and *msx1* has been shown to act upstream of early neural crest markers such as *snail*, *slug*, and *foxd3* [10]. In zebrafish, Msx-family genes are expressed in partially overlapping neural crest and preplacodal ectoderm domains, and their disruption affects both neural crest differentiation and cranial placode development [11]. These findings make it plausible that Trigeminal placode cells adjacent to Neural Crest cells retain higher *msx1b* expression, consistent with the neighborhood-associated expression pattern identified by COSMIC. Importantly, the same neighborhood-defined groups also differed in image-derived morphology, most prominently in nuclear texture (CellProfiler feature Granularity, $p < 5 \times 10^{-10}$). Thus, *msx1b* represents a morphology-associated gene whose expression marks a local neural-crest-adjacent developmental state that is also reflected in nuclear texture within Trigeminal placode cells.

**Gene *foxb1a* within ventral forebrain.** COSMIC identified *foxb1a* as morphology-predictable within annotated forebrain ventral cells (Supplementary Fig.24b). Forebrain-ventral cells near Diencephalon cells had higher *foxb1a* expression than other forebrain-ventral cells (mean normalized expression 2.66 vs. 0.81; $p < 5 \times 10^{-31}$). This is consistent with prior work showing that *Foxb1* expression, after broad early expression in the developing neural tube, becomes restricted to the ventral and caudal diencephalon and marks neuroepithelial domains that contribute to regionalized diencephalic nuclei [12]. Thus, higher *foxb1a* expression in ventral forebrain cells neighboring Diencephalon cells is biologically plausible, as these cells lie near a regional boundary associated with diencephalic identity. The same neighborhood-defined groups also differed in nuclear shape (CellProfiler feature Central moment, $p < 2 \times 10^{-4}$). Thus, *foxb1a* links a local diencephalon-adjacent transcriptional state to measurable variation in nuclear shape within the annotated ventral forebrain.

**Gene *sox2* within lens placode.** COSMIC identified *sox2* as morphology-predictable within annotated placode lens cells (Supplementary Fig.24c). Lens-placode cells near optic cup cells had higher *sox2* expression than other lens-placode cells (mean normalized expression 3.09 vs. 1.73; $p < 1 \times 10^{-31}$). This is consistent with classical lens-induction studies showing that, after close association with the optic vesicle, *Sox2* and *Sox3* are activated specifically in the vesicle-facing head ectoderm, and that these *Sox2/3*-positive, *Pax6*-positive ectodermal cells subsequently give rise to the lens placode [13]. Removal of the prospective retina region eliminates *Sox2/3* expression in the corresponding head ectoderm, supporting the idea that optic tissue provides an inductive signal for *Sox2* activation [13]. Therefore, the higher *sox2* expression observed in lens-placode cells neighboring optic cup cells is consistent with a local optic-cup-associated lens induction program. The same neighborhood-defined groups also differed in nuclear edge intensity (CellProfiler feature Edge intensity, $p < 8 \times 10^{-15}$). Thus, *sox2* is not only spatially consistent with optic-cup-mediated lens induction, but is also associated with a nuclear morphology difference within lens placode cells.

**Gene *cdx4* within posterior hindbrain.** COSMIC identified *cdx4* as morphology-predictable within annotated Hindbrain R7 cells (Supplementary Fig.24d). Hindbrain-R7 cells near formed somites had higher *cdx4* expression than other Hindbrain-R7 cells (mean normalized expression 2.56 vs. 0.40; $p < 3 \times 10^{-48}$). This is consistent with prior work establishing Cdx factors as posteriorizing regulators of embryonic patterning. *cdx4* regulates *hox* gene expression during zebrafish development [14], and Cdx factors are required in the neural ectoderm to specify spinal cord identity at the expense of hindbrain identity. Loss of *cdx1a* and *cdx4* causes posterior expansion of hindbrain character and reduction of spinal cord identity, whereas *cdx4* overexpression in the hindbrain disrupts rhombomere patterning and induces spinal cord-specific genes [15]. These observations make elevated *cdx4* expression in posterior Hindbrain R7 cells near formed somites biologically plausible, as this region lies close to the hindbrain-spinal cord transition. The same neighborhood-defined groups also differed in nuclear edge intensity (CellProfiler feature Edge intensity, $p < 4 \times 10^{-25}$). Thus, *cdx4* connects a posterior boundary-associated transcriptional program with measurable nuclear morphology variation within Hindbrain R7 cells.

Across these examples, the neighborhood analysis provides a biological interpretation of the genes nominated by COSMIC. Each gene marks a local developmental context supported by prior literature, and in each case the same neighborhood-defined cells also show a measurable difference in DAPI-derived nuclear morphology. Thus, COSMIC does not simply recover spatially patterned genes; it identifies genes whose context-dependent expression is accompanied by nuclear morphology differences within a single annotated tissue type. Together with *anp32e*, these results support COSMIC as a framework for nominating biologically interpretable morphology-associated genes in tissue-embedded single-cell data.

#### Supplementary Note 5 Evaluation metrics

We evaluated COSMIC using different metrics, including Coverage (COV) [16], Sliced Wasserstein Distance (SWD) [17], $k$-Nearest Neighbor Accuracy ($k$-NNA) [18], gene-level correlation, and classification accuracy. Unless stated otherwise, all distances are computed in the embedding spaces defined by the nuclear encoder or the transcriptomic encoder using the Euclidean distance.

**Distributional fidelity and diversity.** To assess the distributional fidelity and diversity of COSMIC's generated outputs relative to real data, we computed COV, SWD, and $k$-NNA. Coverage (COV) [16] measures the proportion of real samples that are the nearest neighbour of at least one generated sample. Formally, given real samples $\mathbf{X}_{\text{real}} = \{\mathbf{x}_i\}_{i=1}^N$ and generated samples $\mathbf{X}_{\text{gen}} = \{\hat{\mathbf{x}}_j\}_{j=1}^M$, we define

$$\text{COV}_d(\mathbf{X}_{\text{real}}, \mathbf{X}_{\text{gen}}) = \frac{1}{N} \sum_{i=1}^{N} \mathbf{1}[\,\exists\, j \in \{1, \dots, M\} \text{ s.t. } i = \hat{\mathbf{x}}_j \mathbf{X}_{\text{real}}]\,.$$

where $\hat{\mathbf{x}}_j \mathbf{X}_{\text{real}} = \arg\min_{\mathbf{x} \in \mathbf{X}_{\text{real}}} d(\hat{\mathbf{x}}_j, x)$ is the (index of the) nearest real point to $\hat{\mathbf{x}}_j$. In all our experiments, we set $M = 10N$, sampling the model more densely than the data to obtain a more stable empirical estimate of coverage, in line with prior practice of using many more generated than real samples for evaluation [19].

We measure fidelity with the Sliced Wasserstein distance (SWD) [17]. Intuitively, SWD compares two point sets by repeatedly *(i)* projecting both sets onto a random one-dimensional direction, *(ii)* sorting the projected points, and *(iii)* averaging the absolute gaps between the two sorted lists. Doing this over many random directions and averaging the results yields a stable distance: a small SWD means that the generated embeddings align well with the real ones.

Formally, let $\{\boldsymbol{\theta}_k\}_{k=1}^K$ be unit vectors drawn at random from the unit sphere in the embedding space. For each direction $\boldsymbol{\theta}_k$, we project the real and generated samples to obtain one-dimensional point sets

$$u^{(k)} = \{\langle \boldsymbol{\theta}_k, \mathbf{x}_i \rangle\}_{i=1}^N, \qquad v^{(k)} = \{\langle \boldsymbol{\theta}_k, \hat{\mathbf{x}}_j \rangle\}_{j=1}^M.$$

We then compute the one-dimensional Wasserstein–1 distance $W_1(u^{(k)}, v^{(k)})$ by sorting and pairing the projected points. The sliced Wasserstein distance between the two distributions is defined as the expectation of this quantity over random directions and is approximated in practice by the empirical average

$$\text{SWD}(\mathbf{X}_{\text{real}}, \mathbf{X}_{\text{gen}}) = \frac{1}{K} \sum_{k=1}^{K} W_1\big(u^{(k)}, v^{(k)}\big).$$

To normalize the scores to $[0, 1]$, we divide all SWD values by $\mathrm{SWD}(\mathbf{X}_{\text{real}}, \mathbf{X}_{\text{gauss}})$, where $\mathbf{X}_{\text{gauss}}$ is sampled from a Gaussian distribution with the same mean and variance as $\mathbf{X}_{\text{real}}$.

We also use Fréchet Inception Distance (FID) [20] to quantify the global discrepancy between real and generated image distributions. Unlike the original FID formulation, which uses an Inception network trained on natural images, we computed FID in the embedding space of our pre-trained nuclear encoder. This choice provides a domain-specific evaluation because the encoder was trained on large-scale nuclear microscopy images and therefore captures biologically relevant variation in nuclear morphology. Given feature embeddings from real images and generated images, we approximate each distribution as a multivariate Gaussian with means $\boldsymbol{\mu}_{\text{real}}$ and $\boldsymbol{\mu}_{\text{gen}}$ and covariance matrices $\boldsymbol{\Sigma}_{\text{real}}$ and $\boldsymbol{\Sigma}_{\text{gen}}$. FID is then defined as

$$\mathrm{FID}(\mathbf{X}_{\text{real}}, \mathbf{X}_{\text{gen}}) = \|\boldsymbol{\mu}_{\text{real}} - \boldsymbol{\mu}_{\text{gen}}\|_2^2 + \mathrm{Tr}\left(\boldsymbol{\Sigma}_{\text{real}} + \boldsymbol{\Sigma}_{\text{gen}} - 2\left(\boldsymbol{\Sigma}_{\text{real}}\boldsymbol{\Sigma}_{\text{gen}}\right)^{1/2}\right).$$

Lower FID values indicate that the generated images more closely match the real image distribution in the pretrained nuclear morphology feature space.

To quantify how well real and generated samples mix in feature space, we use a $k$-nearest neighbor accuracy (k-NNA) metric extended from 1-NNA [18]. Let $\mathbf{X} = \mathbf{X}_{\text{real}} \cup \mathbf{X}_{\text{gen}}$ denote the union of real and generated samples, and let $y(x) \in \{0, 1\}$ be the domain label (real or generated). For each sample $\mathbf{x} \in \mathbf{X}$, we consider its $k$ nearest neighbors $\{\mathbf{x}_{(i)}\}_{i=1}^{k}$ in $\mathbf{X} \setminus \{\mathbf{x}\}$ and compute the fraction of neighbors that come from the same domain. The k-NNA score is defined as

$$\text{k-NNA}_k = \mathbb{E}_{\mathbf{x}\in\mathbf{X}}\left[\frac{1}{k}\sum_{i=1}^{k}\mathbf{1}\left[y(\mathbf{x}_{(i)}) = y(\mathbf{x})\right]\right]. \tag{1}$$

For balanced real and generated sets with matching cardinality (that is, $|\mathbf{X}_{\text{real}}| = |\mathbf{X}_{\text{gen}}|$), an ideal generator yields k-NNA$_k \approx 0.5$, which indicates that local neighborhoods are well mixed across domains. Larger values of k-NNA$_k$ indicate that neighborhoods are dominated by a single domain and that the two distributions are more easily separable. In all experiments, we set $k = 0.05 \times (|\mathbf{X}_{\text{real}}| + |\mathbf{X}_{\text{gen}}|)$, that is, we use the $5\%$ nearest neighbors of the pooled dataset.

**Gene-level correlation.** To evaluate transcriptome generation from images, we computed the Pearson correlation between predicted and ground-truth gene expression vectors. When evaluating across cell lines, for each gene $g$, we define

$$\text{P-corr}_g = \mathrm{PearsonCorr}(\mathbf{x}_g^{\text{RNA}}, \hat{\mathbf{x}}_g^{\text{RNA}}),$$

where $\mathbf{x}_g^{\mathrm{RNA}}$ and $\hat{\mathbf{x}}_g^{\mathrm{RNA}}$ are the real and generated expression values for gene $g$ across cells. The resulting distribution of correlation scores reflects gene-level accuracy and informs the extent to which morphological features constrain transcriptional variability.

When evaluating within each cell line, we calculate the Pearson correlation separately for each experimental batch and then average the correlation scores across batches. This reduces the influence of batch effects on the evaluation scores. Where appropriate, we assess statistical significance using the standard two-sided $t$-test for the Pearson correlation coefficient, with multiple-hypothesis correction applied to the resulting adjusted $p$-values.

**Cell-type classification.** To assess how well generated images capture cell type information, we trained a cell type classifier on real data and evaluated its performance on generated data. Specifically, we train an image cell type classifier (a four-layer convolutional backbone followed by a two-layer fully connected head) on ground-truth nuclear images and tested it on generated images to determine whether cell type discriminative visual features are preserved. Similarly, for transcriptome prediction, we train a two-layer multilayer perceptron (MLP) as the transcriptome cell type classifier and test it on generated transcriptomes to evaluate whether cell type discriminative molecular signatures are preserved.

**Cell cycle angular score calculation.** Following [21], we assign each cell a continuous cell cycle phase on the interval $[0, 2\pi)$. We first apply a log1p transformation to FUCCI intensities, then project them onto the orthonormal basis given by the first two principal components (PC1 and PC2). For each cell, we compute the angle

$$\theta = \mathrm{arctan2}(\mathrm{PC2}, \mathrm{PC1}),$$

which yields a continuous angular score along the cell cycle trajectory. The orientation of the angle is chosen such that phases assigned from FUCCI gating (for example G1, S, G2, M) align with the expected order as described in [21].

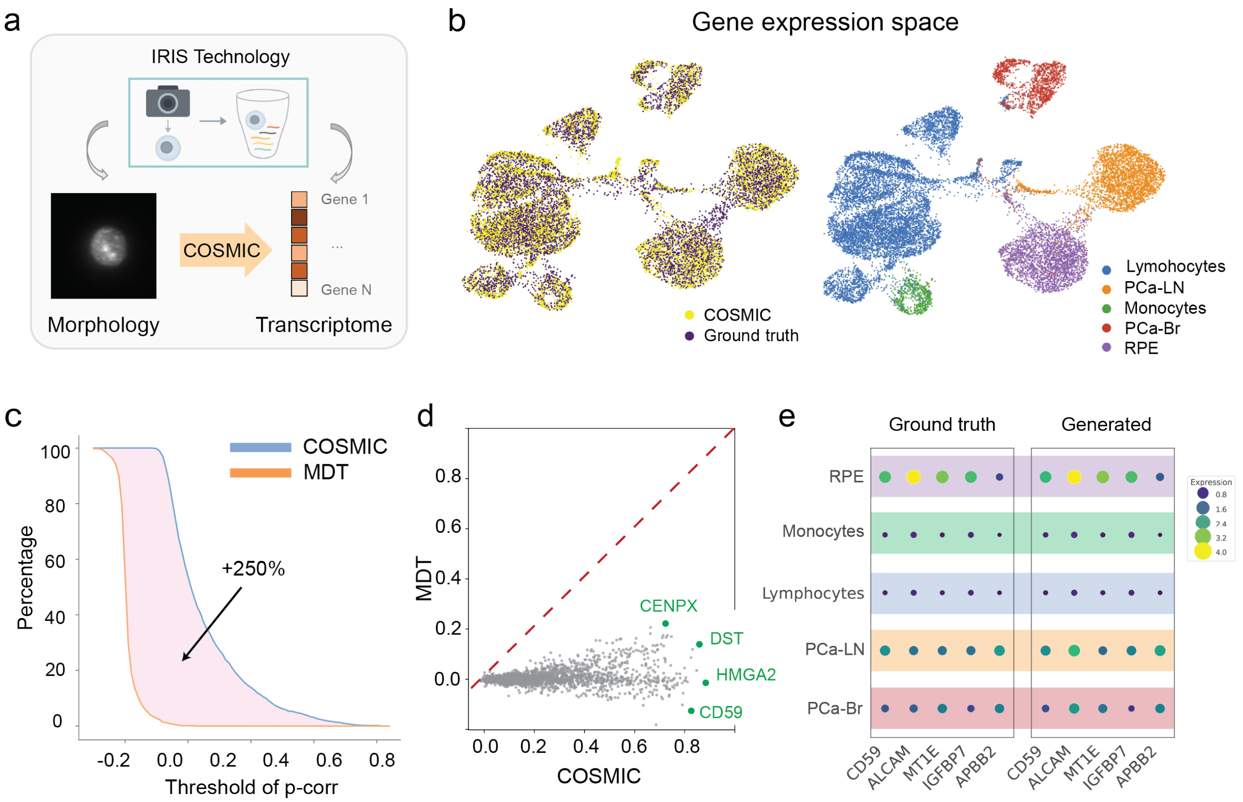


**Supplementary Figure 1:** **(a)** Results of generating transcriptomic profiles of cells from nuclear images on human data. **(b)** UMAP of human data showing overlap between COSMIC-generated expression (yellow) and ground-truth RNA profiles (purple), and the same embedding colored by cell type (right: lymphocytes, PCa-LN, monocytes, PCa-Br, RPE). **(c)** Cumulative distribution of per-gene Pearson correlations on human data comparing COSMIC to the MDT baseline; COSMIC yields a substantially larger fraction of positively correlated genes, corresponding to a $250\%$ accumulated improvement from a correlation threshold of $0$. **(d)** Per-gene performance comparison (MDT vs COSMIC); points below the diagonal indicate COSMIC outperforms MDT. Example genes are annotated (*CENPX*, *DST*, *HMGA2*, *CD59*). **(e)** Cell-type–level heat map of top predictable genes, showing that COSMIC preserves characteristic expression patterns relative to ground truth, consistent with accurate recovery of cell-type signatures.

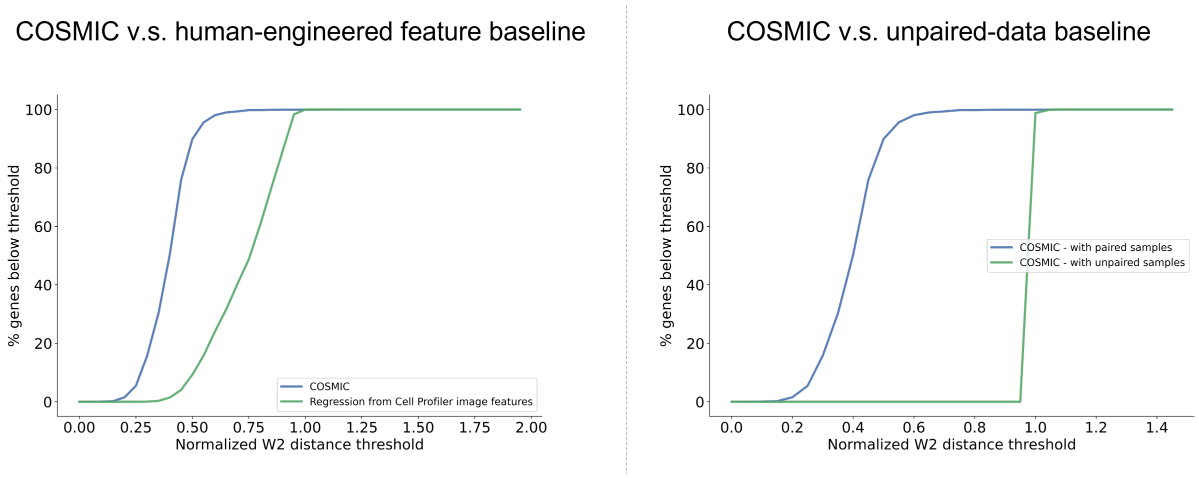


**Supplementary Figure 2:** We evaluated two additional baselines for the task of generating transcriptomes from nuclear images on IRIS human cells. First, we applied a regression model to predict transcriptomes from CellProfiler features. Specifically, we extracted 271 morphological features, ranging from commonly used descriptors such as Area and Compactness to less frequently used measures such as PerimeterCrofton. Second, we tested a random-matching baseline in which transcriptomes and nuclear images were paired at random before training COSMIC. The curves above summarize gene-level predictive performance across normalized Wasserstein-2 distance thresholds. For any threshold on the x-axis, the y-axis shows the fraction of genes with normalized Wasserstein-2 distance below that value, so curves that rise more quickly indicate better performance. COSMIC consistently outperforms both simple baselines, highlighting the benefit of the COSMIC architecture and paired multimodal data.

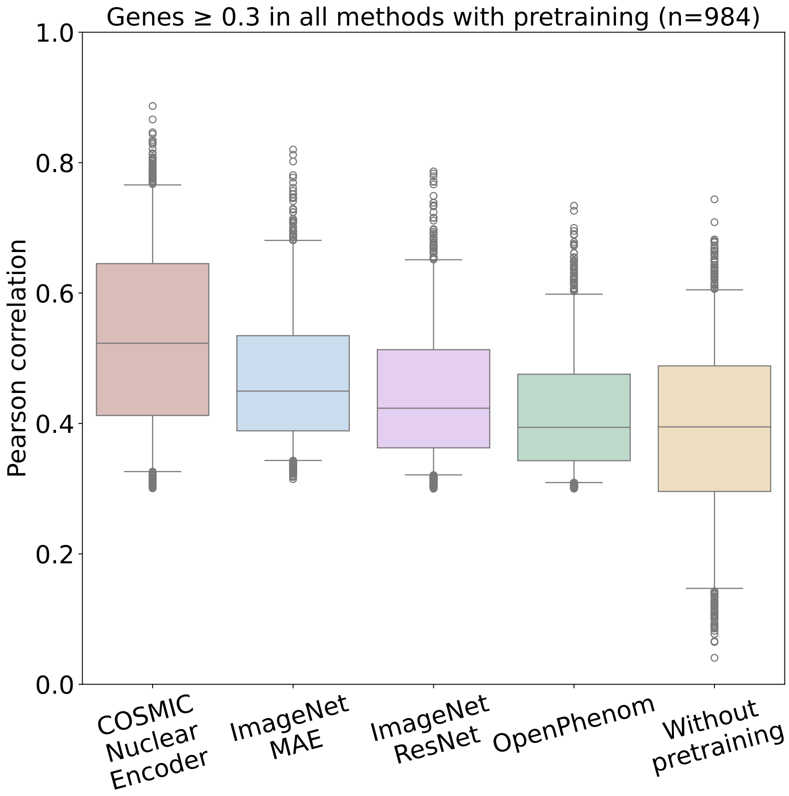


**Supplementary Figure 3:** Comparison of four image encoders for generating nuclear images from transcriptomic profiles: COSMIC Nuclear Encoder, ImageNet MAE [22], ImageNet ResNet [23], and OpenPhenom [24], and the MAE without pretraining. The boxplot shows gene-wise Pearson correlations for genes with Pearson correlation $\geq 0.3$ across all methods with pretraining. (The without-pretraining model serves as a weak baseline and would filter out many genuinely morphology-associated genes simply due to limited representation capacity. This choice allows us to evaluate whether each encoder can recover genes with measurable image-associated signals, while still including the without-pretraining baseline as a comparison.) COSMIC Nuclear Encoder achieves the highest distribution of correlations, indicating that nuclear morphology-focused pretraining provides an advantage over generic natural-image pretraining and existing morphology encoders. These results suggest that the pretrained encoder does not transfer gene-expression labels from unrelated cell lines, but instead provides reusable nuclear image features that can be adapted to IRIS ground-truth paired image-transcriptome data. Each box indicates the interquartile range (25th to 75th percentiles) with the central line marking the median. Whiskers extend to the 5th and 95th percentiles.

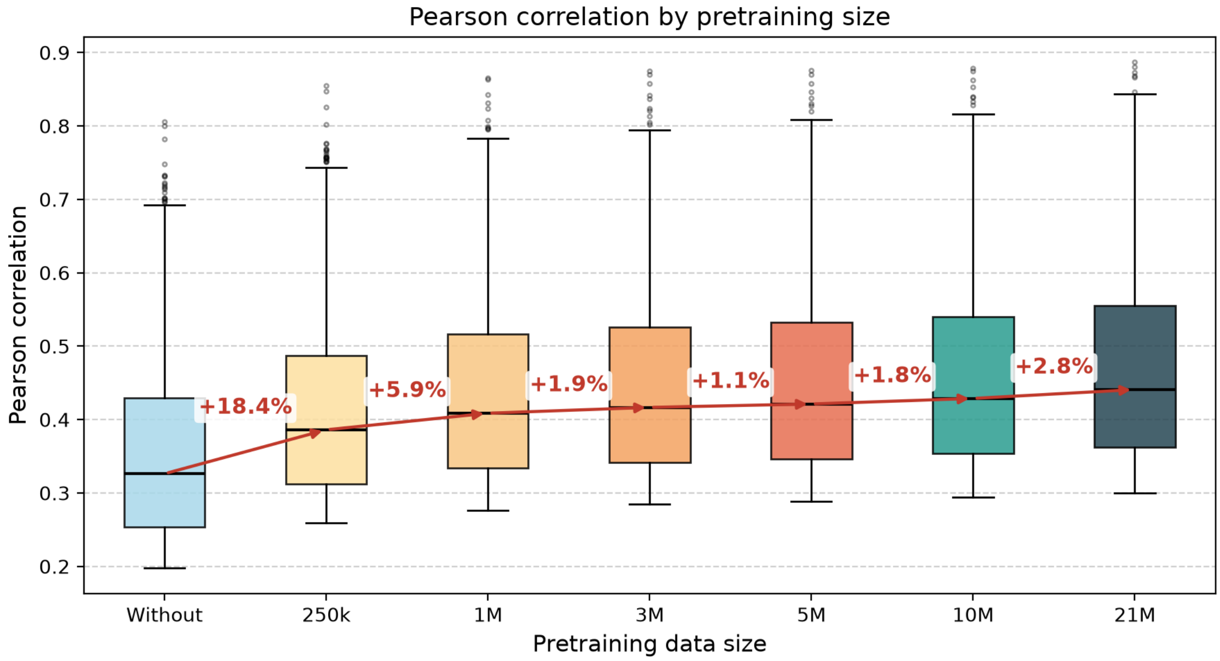


**Supplementary Figure 4:** Effect of nuclear image pretraining dataset size on downstream morphology-to-transcriptome prediction. We evaluated how the number of nuclear images used for pretraining affects downstream image-to-transcriptome prediction performance. COSMIC was trained with nuclear image encoders pretrained using progressively larger subsets of the available pretraining data, including models trained without nuclear-image pretraining and models pretrained with 250K, 1M, 3M, 5M, 10M, and 21M segmented nuclear images. Boxplots show the distribution of gene-wise Pearson correlations on the downstream prediction task across training fractions, using a fixed gene set defined as genes with Pearson correlation $\geq 0.3$ in the 21M training condition. The red arrows and percentage annotations indicate the relative increase in median Pearson correlation between consecutive conditions.

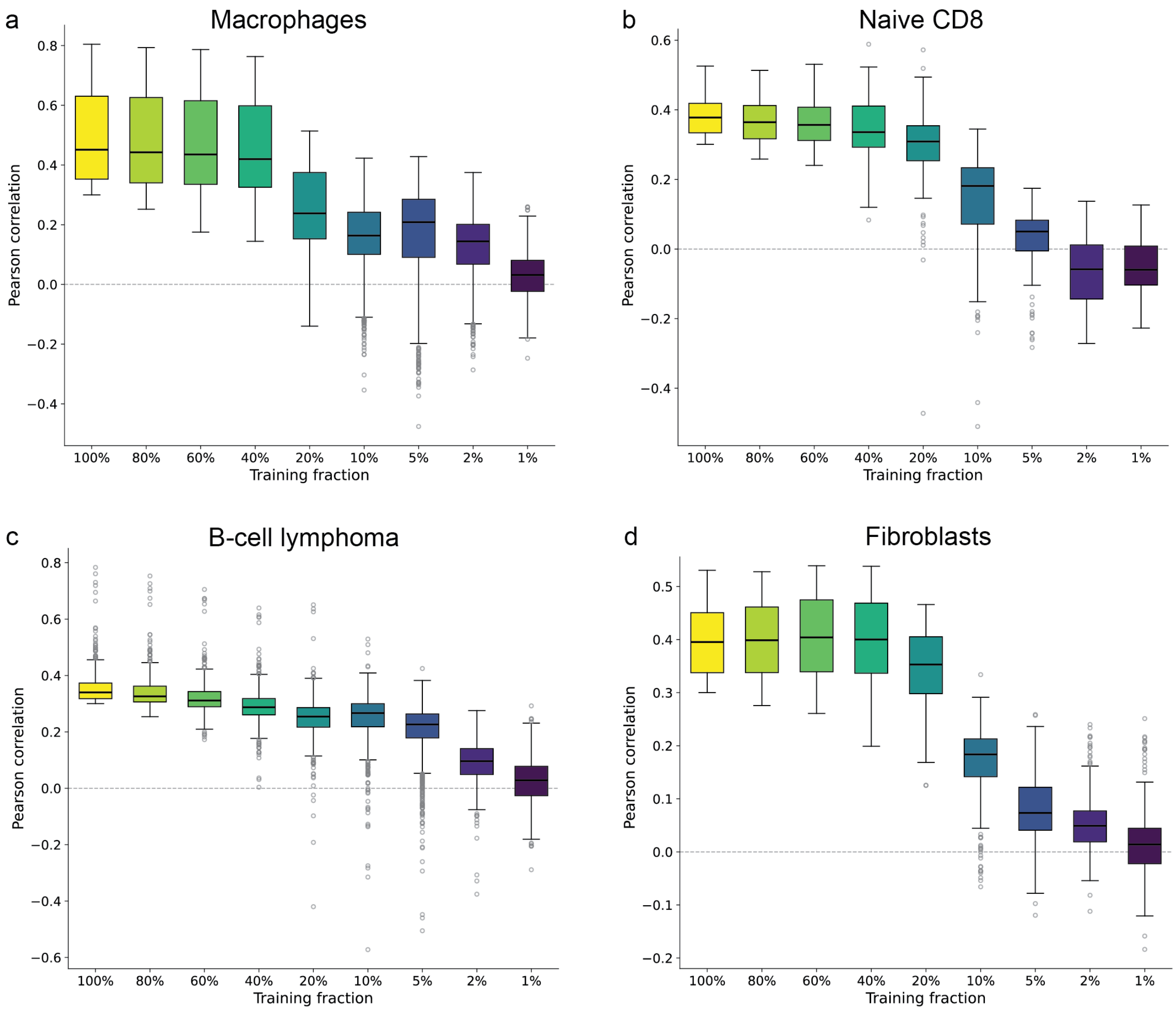


**Supplementary Figure 5:** Data requirements for cross-modal training. We evaluated how the number of paired IRIS cells used for cross-modal training affects image-to-transcriptome prediction performance within each cell type: **(a)** macrophages, **(b)** naive CD8 T cells, **(c)** B-cell lymphoma, and **(d)** fibroblasts. COSMIC was trained using progressively smaller fractions of the IRIS training set, ranging from 100% to 1%, while keeping the evaluation set and training protocol fixed. The boxplots show gene-wise Pearson correlations across training fractions within each cell type, for a fixed gene set defined as genes with Pearson correlation $\geq 0.3$ in the 100% training fraction for that cell type. Prediction performance remains broadly stable when the training set is reduced from 100% to 60%, indicating that the IRIS training data is sufficient for the evaluated cross-modal alignment task. Performance decreases more noticeably when small fractions of the data are used, especially when less than 10%, indicating that sufficient paired training data are required to maintain cross-modal prediction performance.

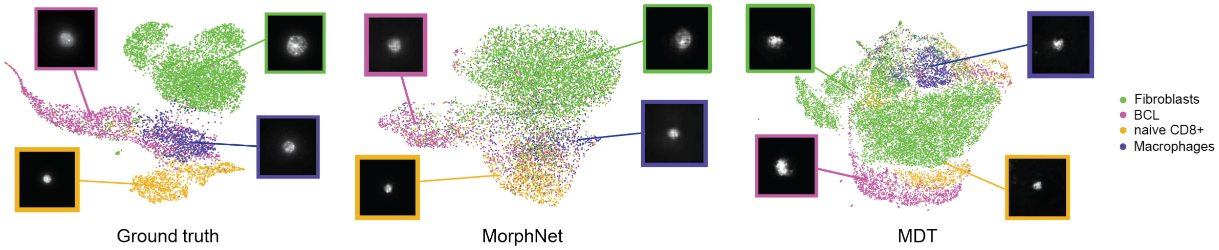


**Supplementary Figure 6:** Embedding space comparison between ground truth images and generated images (from MorphNet and MDT) on the 4,519 mouse cells (test set) obtained with the IRIS technique. Embeddings are extracted using the pretrained COSMIC nuclear encoder and visualized using UMAP. Each point represents a cell, and colors indicate cell types. An example image for each cell type is shown in a box. BCL stands for the B-cell lymphoma.

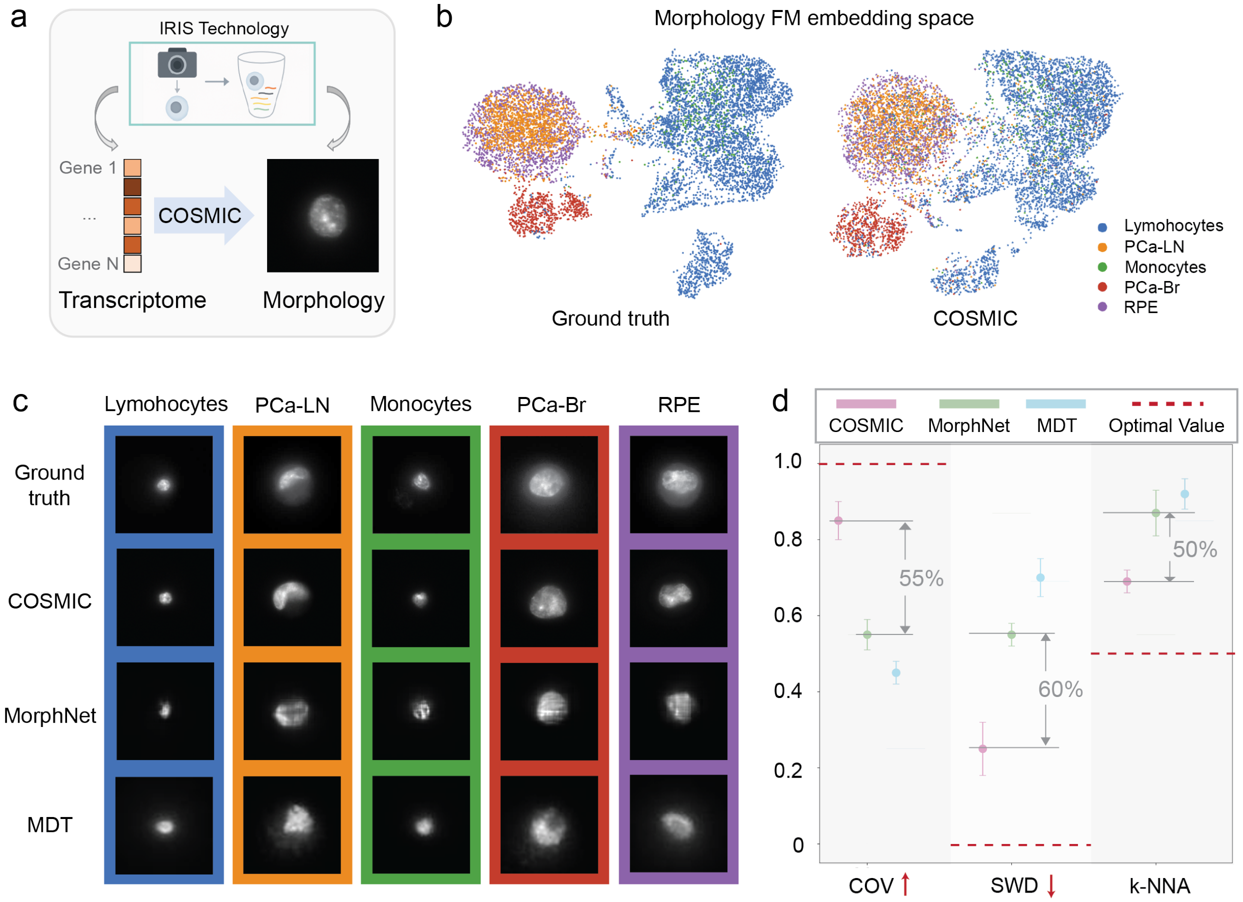


**Supplementary Figure 7:** **(a)** Results of generating nuclear images from transcriptomic profiles on human data. **(b)** UMAP visualizations for the human dataset, showing cell-type structure in the COSMIC nuclear encoder embedding space (left: ground truth; right: COSMIC). Colors denote cell types: lymphocytes (blue), PCa-LN (orange), monocytes (green), PCa-Br (red), and RPE (purple). **(c)** Representative examples of image generation. **(d)** Quantitative evaluation of the quality of generated images of IRIS human cells. We assessed the similarity between ground-truth and generated distributions using coverage rate (COV), sliced Wasserstein distance (SWD), and $k$-nearest neighbor accuracy ($k$-NNA). Higher COV values indicate better coverage, lower SWD values suggest higher fidelity, and k-NNA values closer to 0.5 reflect better mixing between real and generated samples. Red dashes indicate the optimal value for each metric. The numbers represent the improvement achieved by COSMIC over the best alternative baseline. Error bars represent standard deviation across five independent rounds of generation.

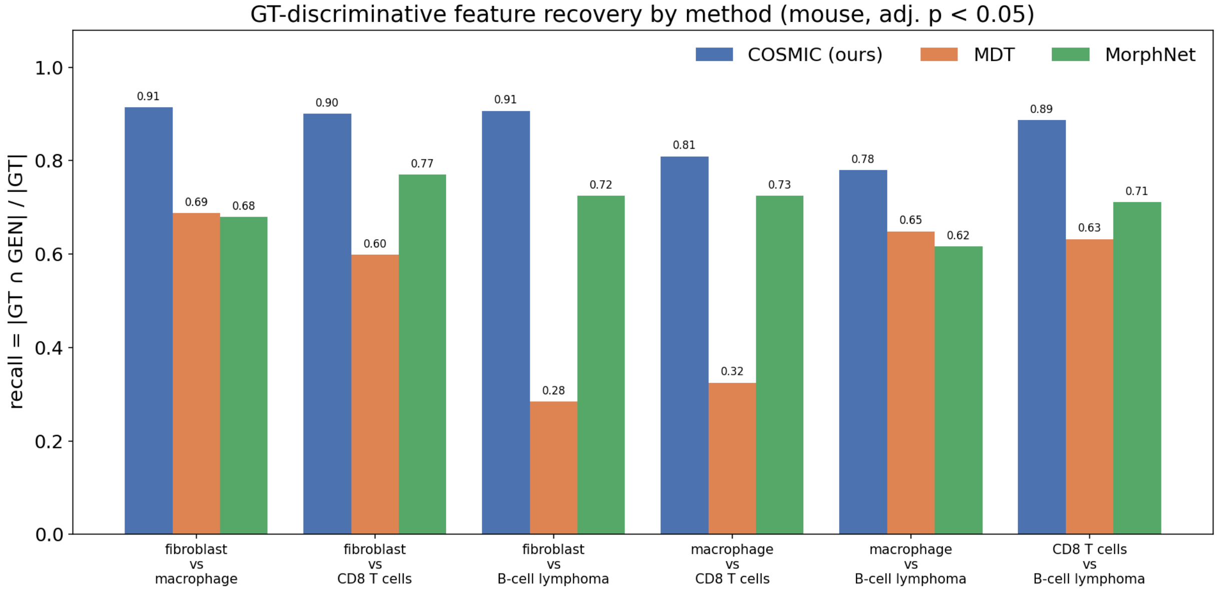


**Supplementary Figure 8: Generated images recover the features that discriminate different cell types.** For each pair of mouse cell types, CellProfiler morphology features were tested for between-cell-type differences using a two-sided Mann–Whitney U test, computed independently for ground-truth (GT) and generated (GEN) images. We considered 271 core features from `get_core_measurements`, of which 264 non-constant features were tested. Three generation methods, COSMIC, MDT, and MorphNet, were evaluated against the same GT reference. A GT-discriminative feature was counted as recovered only if it was significant in both GT and GEN after Benjamini–Hochberg FDR correction (adjusted $p < 0.05$) and the direction of the difference was the same in GT and GEN. Bars show $\mathrm{Recall} = |\mathrm{GT} \cap \mathrm{GEN}|/|\mathrm{GT}|$, representing the fraction of features that significantly distinguish two cell types in real images and also distinguish them, in the same direction, in generated images. Using adjusted $p$-values controls multiple-testing inflation arising from differences in cell numbers, while requiring consistent direction prevents a method from receiving credit for a feature whose cell-type ordering is reversed.

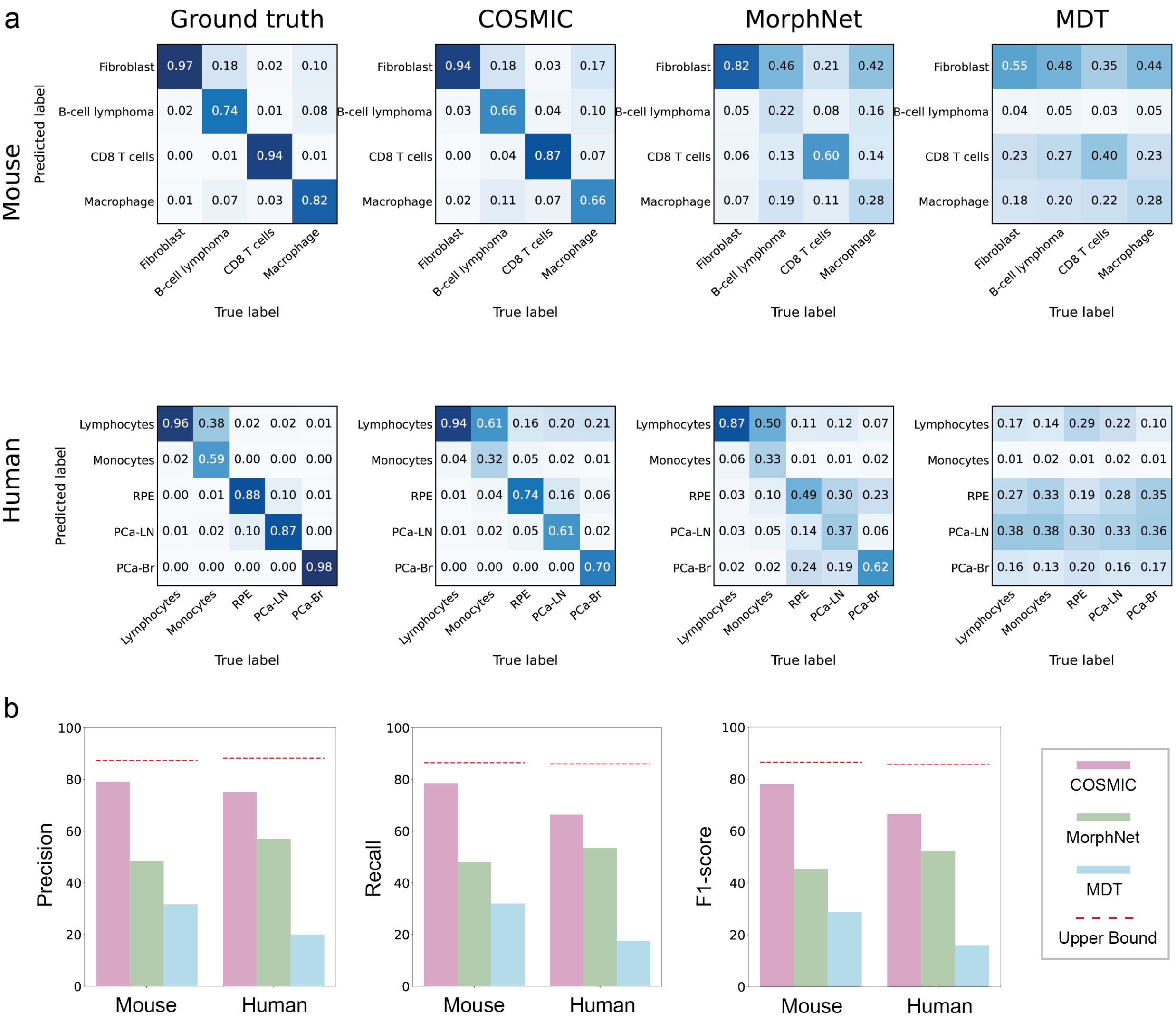


**Supplementary Figure 9:** Comparison of cell-type classification performance using real and generated microscopy images. **(a)** Confusion matrices of a CNN classifier trained on real microscopy images and evaluated on real images (Ground truth) or images generated by COSMIC, MorphNet, and MDT for the mouse (top) and human (bottom) datasets. Rows denote predicted cell types and columns denote ground-truth cell types. **(b)** Macro-averaged precision, recall, and F1-score for cell-type classification using images generated by COSMIC, MorphNet, and MDT on the mouse and human datasets. Colors indicate the generation method, and red dashed lines indicate the corresponding performance on real microscopy images.

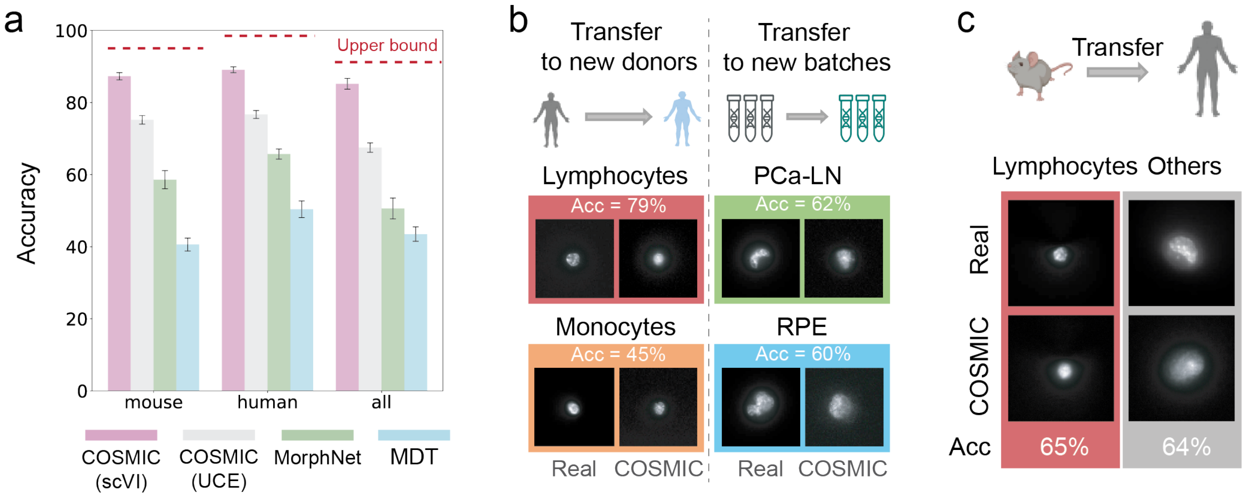


**Supplementary Figure 10:** Evaluation of COSMIC when using the single-cell foundation model UCE [9] as the gene expression encoder. **(a)** Evaluation of COSMIC and alternative methods on the cell type classification task using the IRIS mouse and human datasets. We trained a classifier on ground-truth cell images and cell type annotations. Higher classification accuracy indicates better conditional generation quality. Error bars represent standard deviation across five independent rounds of generation. The red dashed line represents the upper bound obtained by evaluating classification accuracy on the ground-truth images. **(b)** COSMIC generates high-quality nuclear images for unseen batches. We trained COSMIC on IRIS-derived human samples from four cell types, including lymphocytes, monocytes, prostate cancer cells from a lymph node metastasis (PCa-LN), and retinal pigment epithelial cells (RPE), holding out one batch per cell type. For lymphocytes and monocytes, held-out batches correspond to new donors; for PCa-LN and RPE, they correspond to independent experimental replicates. After training, we generated images for these unseen batches and evaluated the quality of the generated images by testing whether a classifier trained on ground-truth images could accurately distinguish the four cell types in the generated set. **(c)** COSMIC generates high-quality nuclear images across species, from mouse to human. We trained COSMIC on mouse data and evaluated its cross-species generalization on human samples. The overlapping cell line between IRIS mouse and human datasets is Lymphocytes (CD8 in mouse cells). COSMIC accurately synthesized human Lymphocytes, with $65\%$ correctly classified as Lymphocyte by our cell type classifier.

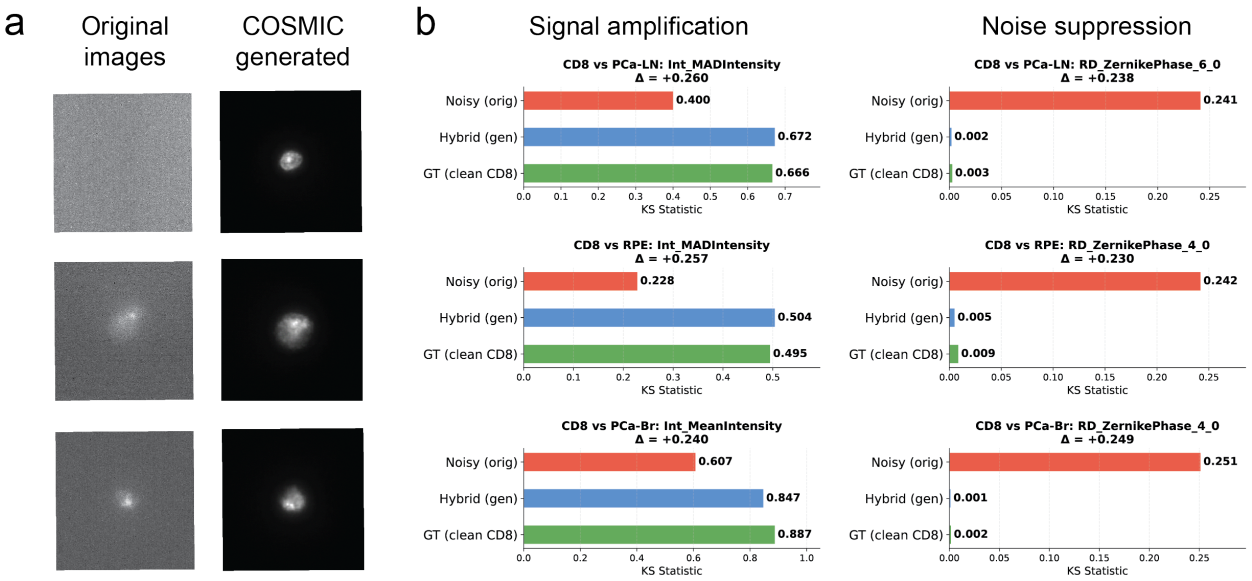


**Supplementary Figure 11:** COSMIC-generated images recover clean-sample feature distributions. **(a)** Representative noisy acquisitions (column 1) and corresponding COSMIC-generated images (column 2) for the three comparison lines (top to bottom: PCa-LN, RPE, PCa-Br). **(b)** Kolmogorov–Smirnov (KS) statistics for the distributional difference of representative morphological features between CD8 cells and each comparison line, comparing original noisy images (red), COSMIC-generated images (blue), and clean CD8 reference (green); $\Delta$ is the change versus the noisy baseline. For intensity features (middle), COSMIC restores the true between-line separation that noise had suppressed, with generated KS values closely matching the clean reference (e.g., 0.672 vs. 0.666). For Zernike phase features (right), the noisy images show a large spurious difference absent in clean data, which COSMIC drives back to near zero to match the reference. Thus COSMIC can replace noisy acquisitions with images whose feature distributions track the clean samples—amplifying meaningful signal while removing noise-derived artifacts.

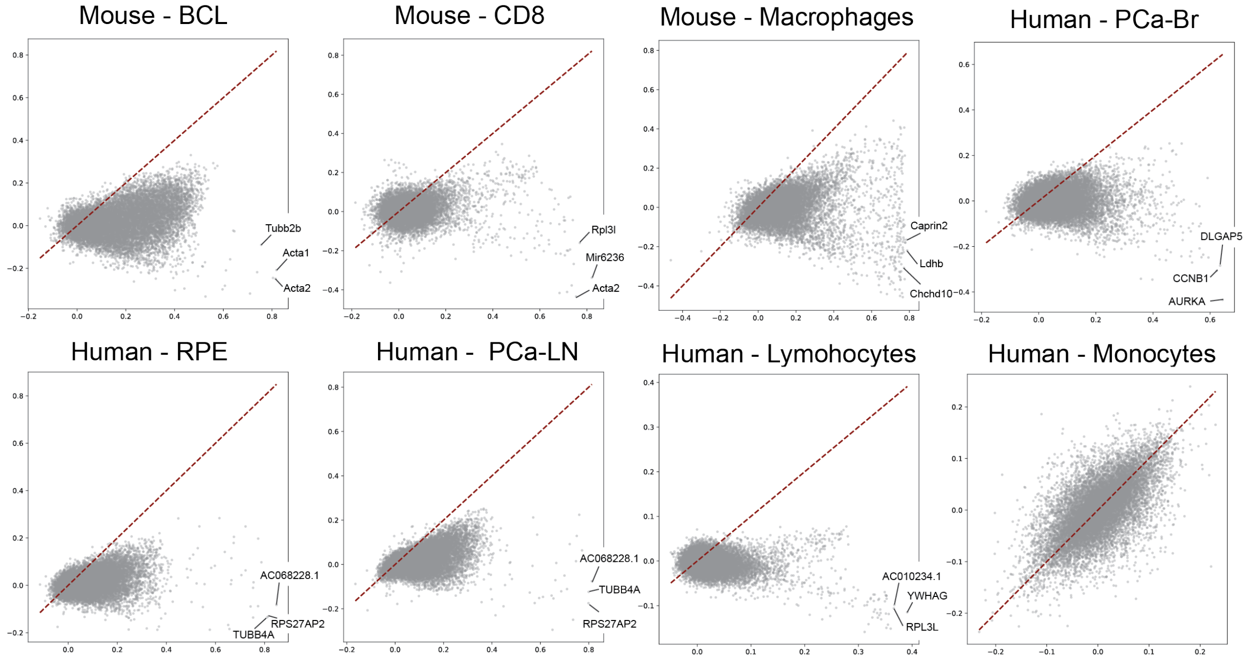


**Supplementary Figure 12:** Performance drop after within–cell-type permutation highlights biological signals beyond cell type. Each panel shows gene-wise performance before (x-axis) vs after permutation (y-axis); the red dashed line is $y = x$. Points below the diagonal indicate losses when true image–expression pairings are broken within a cell type.

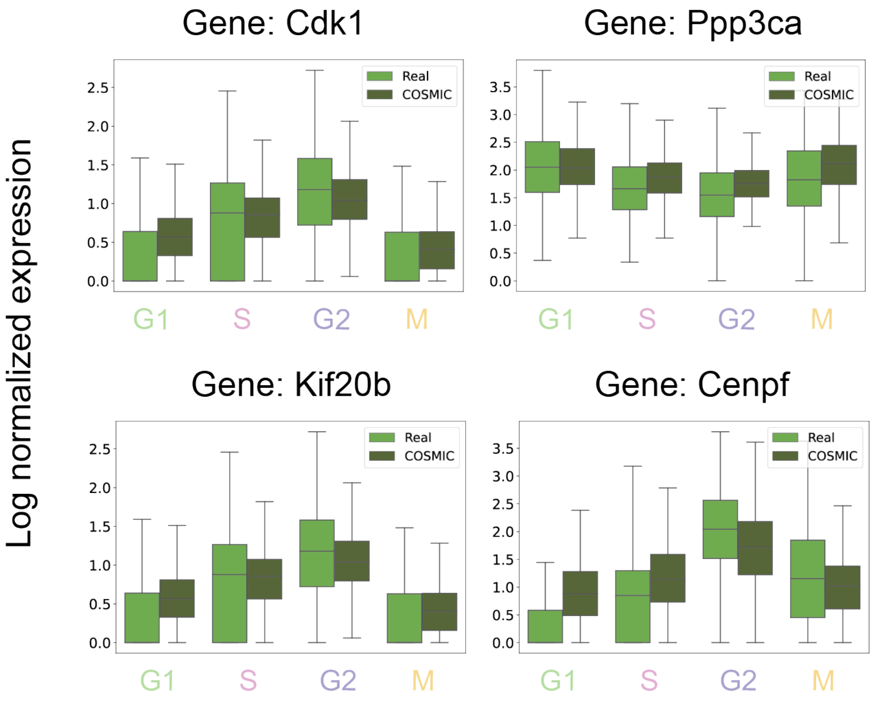


**Supplementary Figure 13:** Transcriptomes predicted by COSMIC across different cell cycle phases. The results include the genes of *Cdk1*, *Ppp3ca*, *Kif20b*, and *Cenpf*. Each box plot illustrates the distribution quartiles of the expression level of one gene across different cell cycle phases (G1: $n = 1,631$ cells, S: $n = 1,036$ cells, G2: $n = 1,366$ cells, M: $n = 748$ cells.) Boxes depict distribution quartiles, with the center line corresponding to the median, and whiskers span 1.5 times the interquartile range.

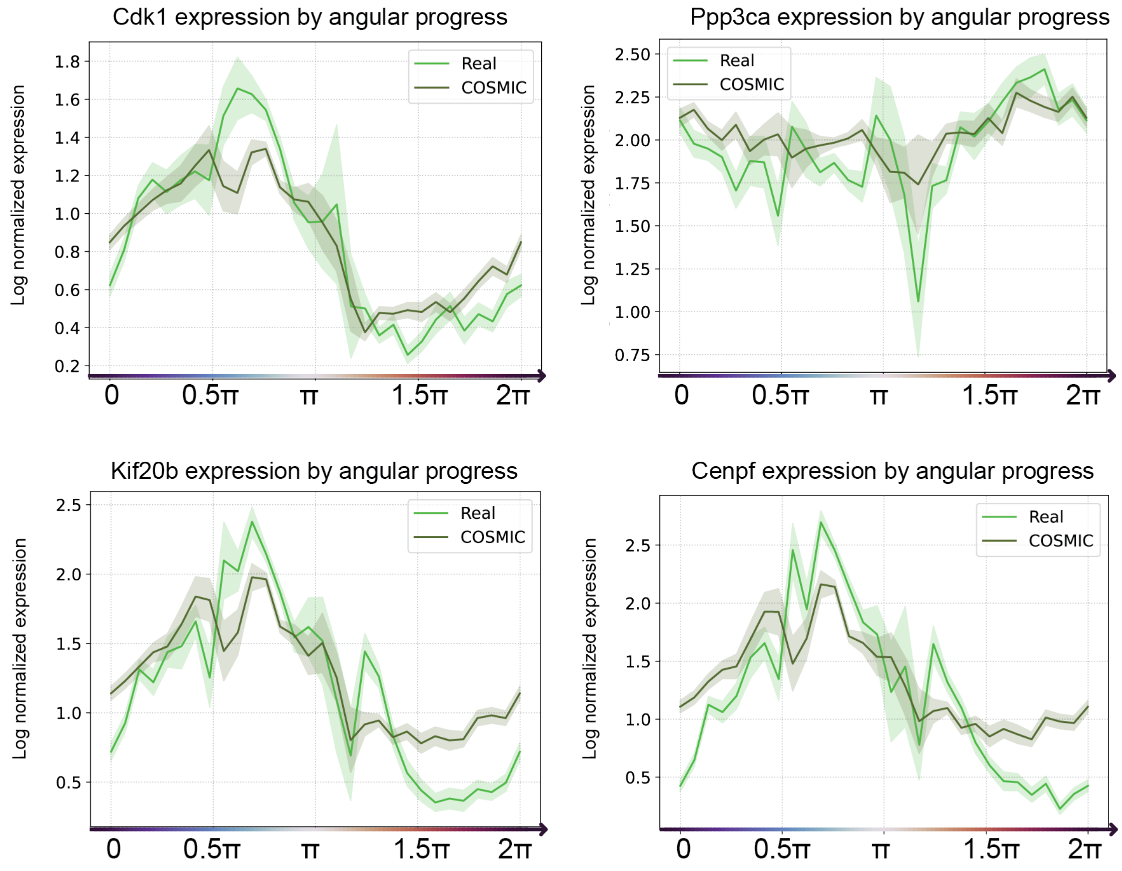


**Supplementary Figure 14:** Cell cycle modeled as a continuous circular trajectory using FUCCI intensities (same as in Fig. 4h). The COSMIC-inferred expression profiles correctly recapitulate continuous expression changes of cell-cycle genes *Cdk1*, *Ppp3ca*, *Kif20b*, and *Cenpf*.

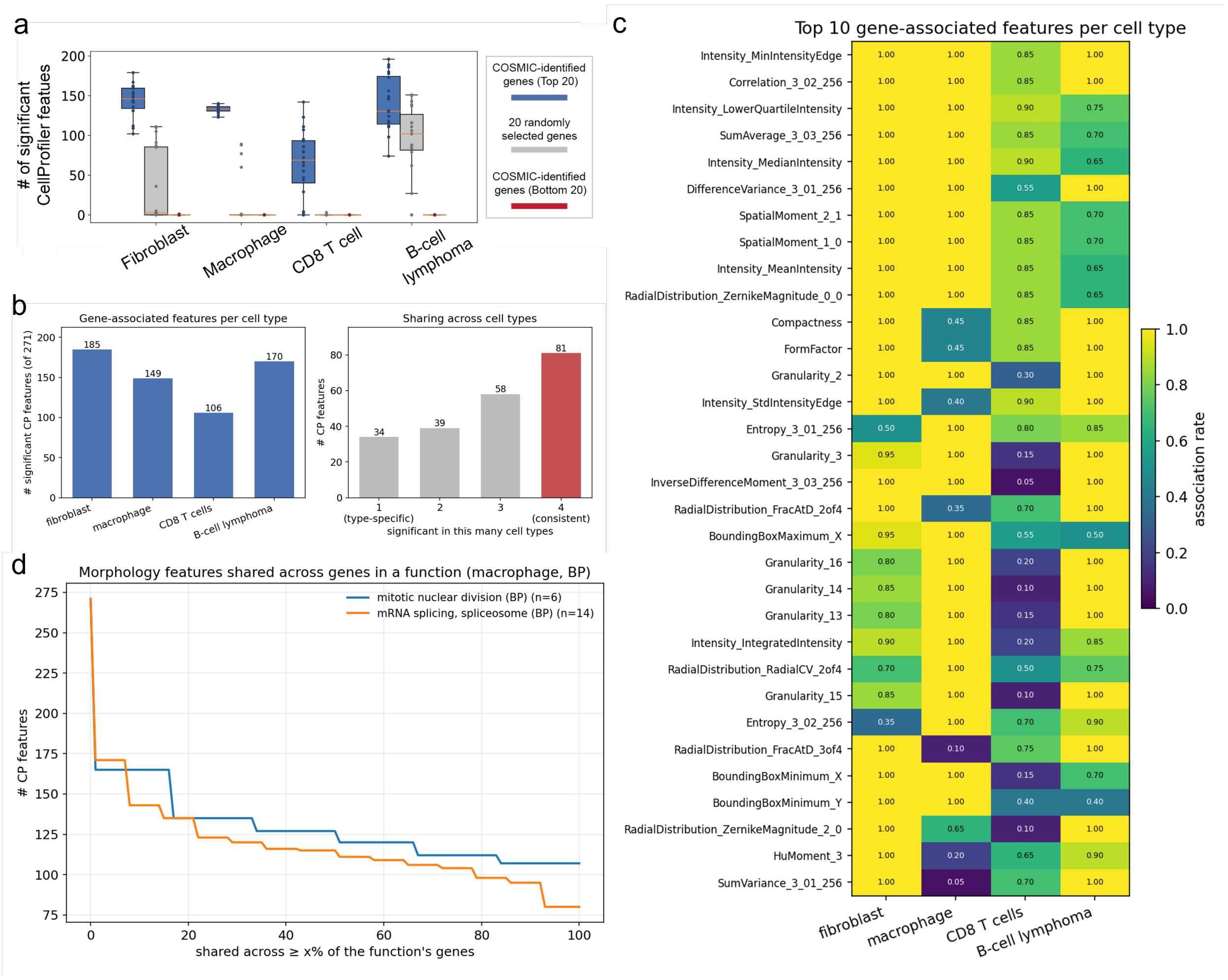
a
### of significant CellProfiler featues
COSMIC-identified genes (Top 20)
20 randomly selected genes
COSMIC-identified genes (Bottom 20)
Fibroblast
Macrophage
CD8 T cell
B-cell lymphoma
b
Gene-associated features per cell type
### significant CP features (of 271)
185
149
106
170
fibroblast
macrophage
CD8 T cells
B-cell lymphoma
Sharing across cell types
### CP features
34
39
58
81
1 (type-specific)
2
3
4 (consistent)
significant in this many cell types
c
Top 10 gene-associated features per cell type
Intensity_MinIntensityEdge
Correlation_3_02_256
Intensity_LowerQuartileIntensity
SumAverage_3_03_256
Intensity_MedianIntensity
DifferenceVariance_3_01_256
SpatialMoment_2_1
SpatialMoment_1_0
Intensity_MeanIntensity
RadialDistribution_ZernikeMagnitude_0_0
Compactness
FormFactor
Granularity_2
Intensity_StdIntensityEdge
Entropy_3_01_256
Granularity_3
InverseDifferenceMoment_3_03_256
RadialDistribution_FracAtD_2of4
BoundingBoxMaximum_X
Granularity_16
Granularity_14
Granularity_13
Intensity_IntegratedIntensity
RadialDistribution_RadialCV_2of4
Granularity_15
Entropy_3_02_256
RadialDistribution_FracAtD_3of4
BoundingBoxMinimum_X
BoundingBoxMinimum_Y
RadialDistribution_ZernikeMagnitude_2_0
HuMoment_3
SumVariance_3_01_256
fibroblast
macrophage
CD8 T cells
B-cell lymphoma
association rate
d
Morphology features shared across genes in a function (macrophage, BP)
mitotic nuclear division (BP) (n=6)
mRNA splicing, spliceosome (BP) (n=14)
### CP features
shared across ≥ x% of the function's genes

**Supplementary Figure 15: Analysis of COSMIC-nominated genes and explainable CellProfiler nuclear-morphology features.** For every gene, cells of a given type were split into high- and low-expression groups at the gene's mean expression, and each of 271 CellProfiler features derived from segmented nuclear masks, including intensity, texture, granularity, shape, and Zernike descriptors, was compared between groups using a two-sided Kolmogorov-Smirnov test with Benjamini-Hochberg correction (FDR $< 0.01$). **(a)** Validation of COSMIC-ranked genes using interpretable nuclear morphology features. Given a gene within a cell type, cells were divided into high- and low-expression groups using the gene's mean expression. The numbers (out of 271 total) of significantly associated CellProfiler features (two-sided Kolmogorov-Smirnov test; FDR $<$ 0.01) are compared between COSMIC's 20 highest-scoring genes, 20 randomly selected genes, and COSMIC's 20 lowest-scoring genes. Notably, the bottom 20 genes identified by COSMIC across all cell types exhibit zero associated CellProfiler features. **(b)** Left: Number of features associated with the top-ranked genes in each cell type. A feature was counted when it was significant for at least $50\%$ of that cell type's top 20 genes. Right: Number of features significant in exactly $n$ of the four cell types. In total, 81 features were consistently associated across all four cell types (red), whereas only 34 were restricted to a single cell type. **(c)** Association rate across cell types, defined as the fraction of the top 20 genes for which each CellProfiler feature was significant. The displayed features represent the union of the ten features with the highest association rates in each cell type. Consistently associated features include intensity descriptors, such as edge and minimum intensity; shape descriptors, including perimeter, solidity, and Feret diameter; and Haralick texture descriptors, including contrast, correlation, and information measure. **(d)** CellProfiler features shared across macrophage genes belonging to Gene Ontology biological processes plausibly related to nuclear morphology, including mitotic nuclear division (GO:0140014; $n = 6$ genes) and mRNA splicing via the spliceosome (GO:0000398; $n = 14$ genes). Curves show the number of features shared across at least $x\%$ of the genes assigned to each function. Between 95 and 107 features were shared across more than $90\%$ of the genes in each function and were dominated by texture, granularity, and nuclear-size descriptors. The functions therefore converge on a common morphology signature rather than distinct function-specific feature sets.

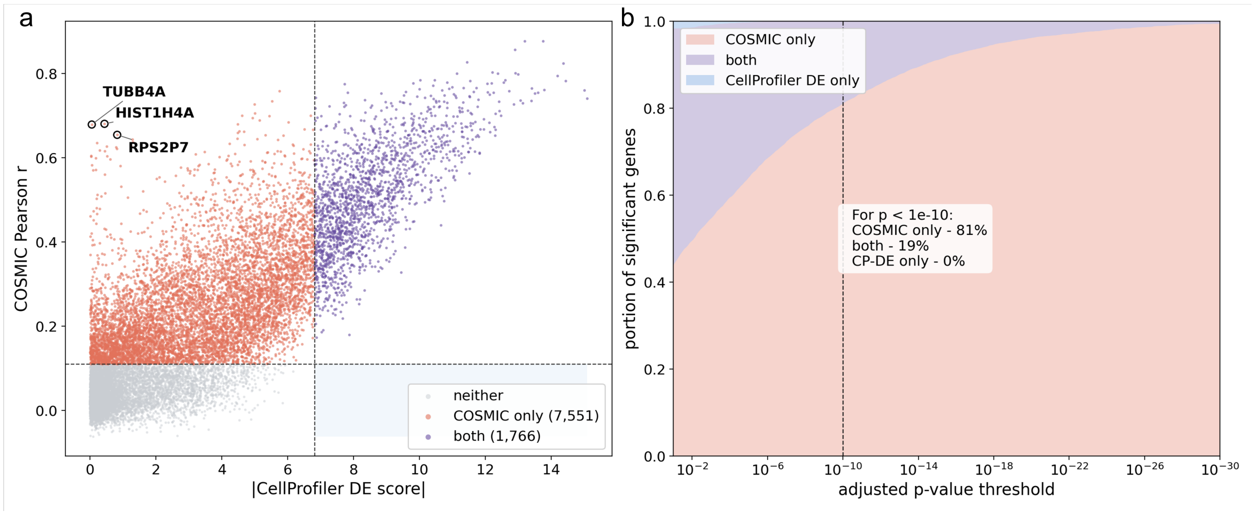


**Supplementary Figure 16:** Comparison of COSMIC-selected morphology-associated genes with genes identified using direct statistical association between gene expression and CellProfiler-derived nuclear morphology features on human data from IRIS. **(a)** Each point represents one of 17,982 genes, positioned according to its CellProfiler differential-expression score ($|z_{\text{Wilcoxon}}|$ for that gene's expression between the two morphological groups defined by $k$-means clustering of CellProfiler features) and its COSMIC Pearson $r$ (the uncalibrated correlation between image-predicted and measured expression). Dashed lines indicate a Benjamini–Hochberg-adjusted $p < 10^{-10}$ threshold applied identically to both methods. This analysis evaluates whether genes prioritized by COSMIC reflect measurable morphology–expression relationships rather than only internal cycle consistency. Genes identified by the direct feature-association baseline provide an interpretable reference set based on handcrafted nuclear morphology descriptors, whereas COSMIC uses learned high-dimensional image representations. All 1,766 genes identified by the feature-association baseline were also recovered by COSMIC (purple), leaving the lower-right region empty and supporting the biological relevance of the COSMIC-nominated gene set. COSMIC additionally identified 7,551 candidates not recovered by the simple feature-based approach (orange), including *TUBB4A*, *HIST1H4A*, and *RPS2P7* (circled). **(b)** Proportions of significant genes identified by COSMIC only, by both methods, or by the CellProfiler DE baseline only across increasingly stringent adjusted $p$-value thresholds. At adjusted $p < 10^{-10}$, 81% of significant genes were identified by COSMIC only and 19% by both methods, with no genes identified by the CellProfiler baseline alone.

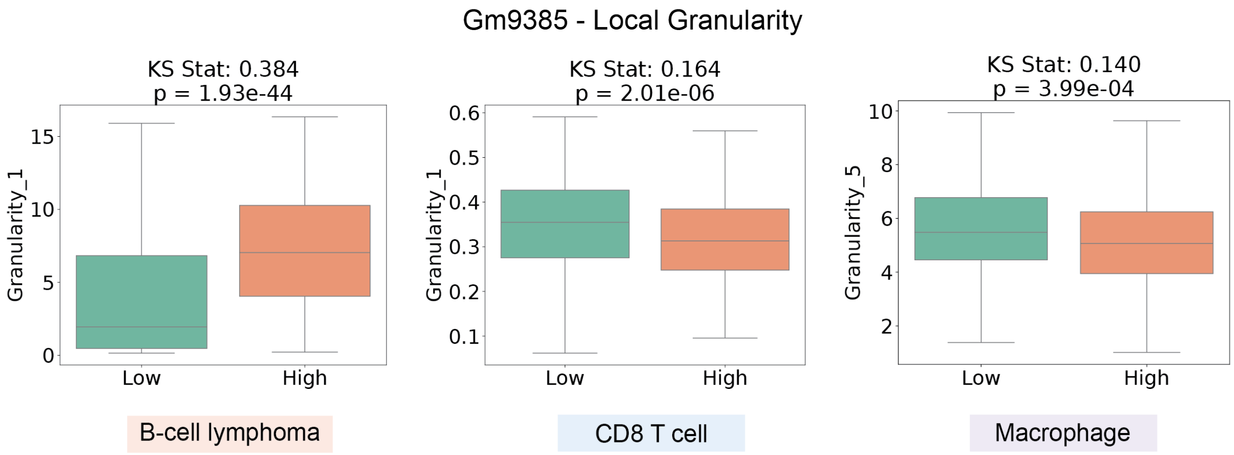


**Supplementary Figure 17:** Cells grouped by low and high expression of the shared morphology-associated gene show consistent differences in local granularity features across immune cell types. In B-cell lymphoma cells, CD8 T cells, and macrophages, CellProfiler-derived local granularity features differ significantly between low- and high-expression groups, indicating that local nuclear texture/granularity is a shared morphology-associated feature across these three immune cell types. KS statistics and p-values are shown above each comparison.

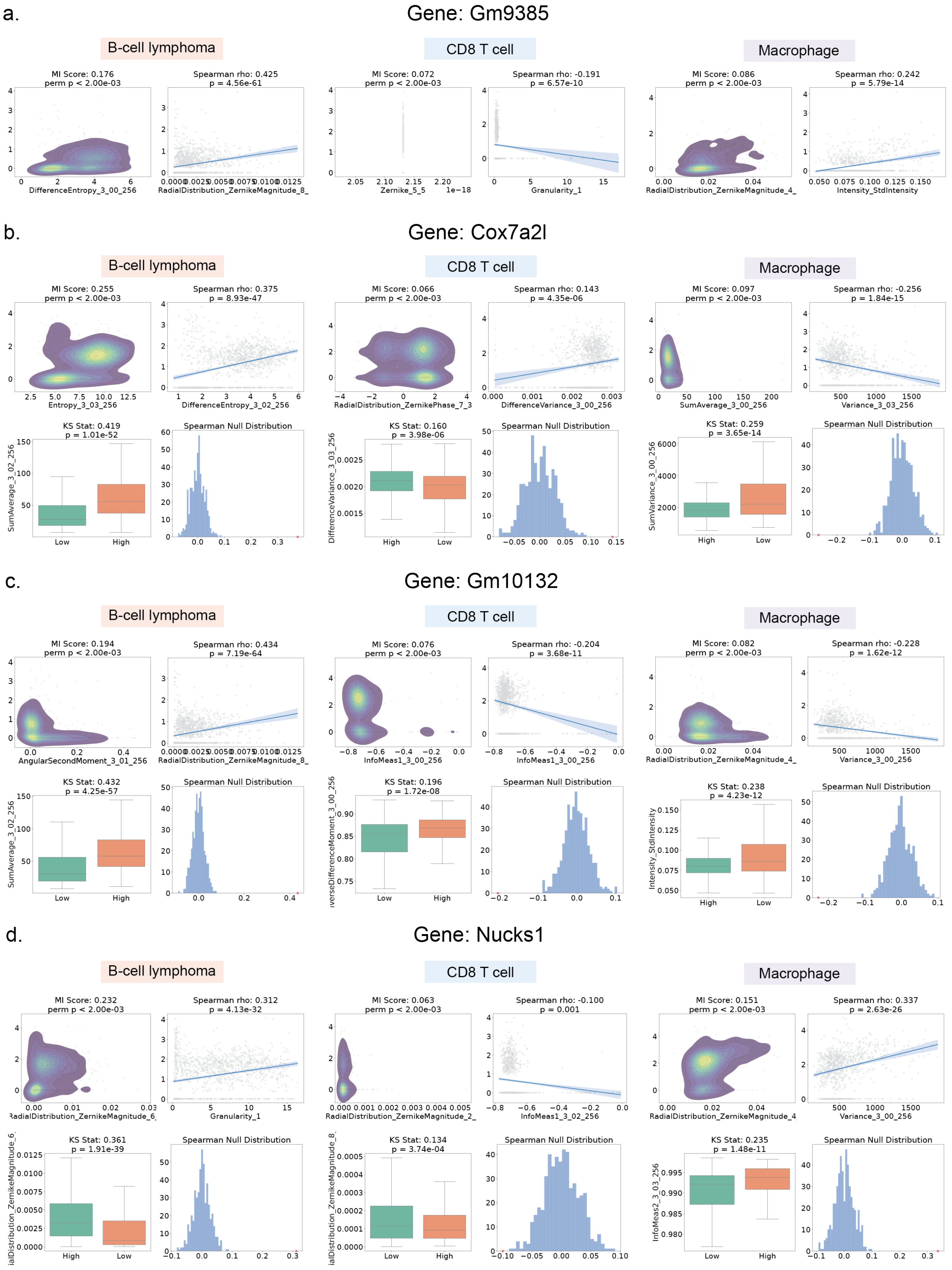

a.
Gene: Gm9385
B-cell lymphoma
CD8 T cell
Macrophage
MI Score: 0.176
perm p < 2.00e-03
DifferenceEntropy_3_00_256
Spearman rho: 0.425
p = 4.56e-61
RadialDistribution_ZernikeMagnitude_8_
MI Score: 0.072
perm p < 2.00e-03
Zernike_5_5
1e−18
Spearman rho: -0.191
p = 6.57e-10
Granularity_1
MI Score: 0.086
perm p < 2.00e-03
RadialDistribution_ZernikeMagnitude_4_
Spearman rho: 0.242
p = 5.79e-14
Intensity_StdIntensity
b.
Gene: Cox7a2l
B-cell lymphoma
CD8 T cell
Macrophage
MI Score: 0.255
perm p < 2.00e-03
Entropy_3_03_256
Spearman rho: 0.375
p = 8.93e-47
DifferenceEntropy_3_02_256
MI Score: 0.066
perm p < 2.00e-03
RadialDistribution_ZernikePhase_7_3
Spearman rho: 0.143
p = 4.35e-06
DifferenceVariance_3_00_256
MI Score: 0.097
perm p < 2.00e-03
SumAverage_3_00_256
Spearman rho: -0.256
p = 1.84e-15
Variance_3_03_256
KS Stat: 0.419
p = 1.01e-52
SumAverage_3_02_256
Low
High
Spearman Null Distribution
KS Stat: 0.160
p = 3.98e-06
DifferenceVariance_3_03_256
High
Low
Spearman Null Distribution
KS Stat: 0.259
p = 3.65e-14
SumVariance_3_00_256
High
Low
Spearman Null Distribution
c.
Gene: Gm10132
B-cell lymphoma
CD8 T cell
Macrophage
MI Score: 0.194
perm p < 2.00e-03
AngularSecondMoment_3_01_256
Spearman rho: 0.434
p = 7.19e-64
RadialDistribution_ZernikeMagnitude_8_
MI Score: 0.076
perm p < 2.00e-03
InfoMeas1_3_00_256
Spearman rho: -0.204
p = 3.68e-11
InfoMeas1_3_00_256
MI Score: 0.082
perm p < 2.00e-03
RadialDistribution_ZernikeMagnitude_4_
Spearman rho: -0.228
p = 1.62e-12
Variance_3_00_256
KS Stat: 0.432
p = 4.25e-57
SumAverage_3_02_256
Low
High
Spearman Null Distribution
KS Stat: 0.196
p = 1.72e-08
nverseDifferenceMoment_3_00_256
Low
High
Spearman Null Distribution
KS Stat: 0.238
p = 4.23e-12
Intensity_StdIntensity
High
Low
Spearman Null Distribution
d.
Gene: Nucks1
B-cell lymphoma
CD8 T cell
Macrophage
MI Score: 0.232
perm p < 2.00e-03
RadialDistribution_ZernikeMagnitude_6_
Spearman rho: 0.312
p = 4.13e-32
Granularity_1
MI Score: 0.063
perm p < 2.00e-03
RadialDistribution_ZernikeMagnitude_2_
Spearman rho: -0.100
p = 0.001
InfoMeas1_3_02_256
MI Score: 0.151
perm p < 2.00e-03
RadialDistribution_ZernikeMagnitude_4
Spearman rho: 0.337
p = 2.63e-26
Variance_3_00_256
KS Stat: 0.361
p = 1.91e-39
ialDistribution_ZernikeMagnitude_6_
High
Low
Spearman Null Distribution
KS Stat: 0.134
p = 3.74e-04
ialDistribution_ZernikeMagnitude_8_
Low
High
Spearman Null Distribution
KS Stat: 0.235
p = 1.48e-11
InfoMeas2_3_03_256
Low
High
Spearman Null Distribution

**Supplementary Figure 18:** Additional examples of COSMIC-nominated genes shared across B-cell lymphoma cells, CD8+ T cells, and macrophages, including **(a)** *Gm9385*, **(b)** *Cox7a2l*, **(c)** *Gm10132*, and **(d)***Nucks1*. For each gene and cell type, the mutual-information density plot shows the relationship between gene expression and the selected CellProfiler-derived morphology feature, where each gray point represents one cell and colored contours indicate cell-density distributions in feature-expression space. Expression-stratified boxplots compare the morphology feature between low- and high-expression cells. Spearman correlation panels quantify the association between gene expression and the morphology feature, with the blue line indicating the fitted regression trend and the shaded region representing the confidence interval. Permutation tests compare the observed Spearman correlation against a null distribution generated by randomly permuting gene-expression values across cells. These examples support that immune-shared COSMIC-nominated genes are associated with interpretable nuclear morphology features across multiple immune cell types.

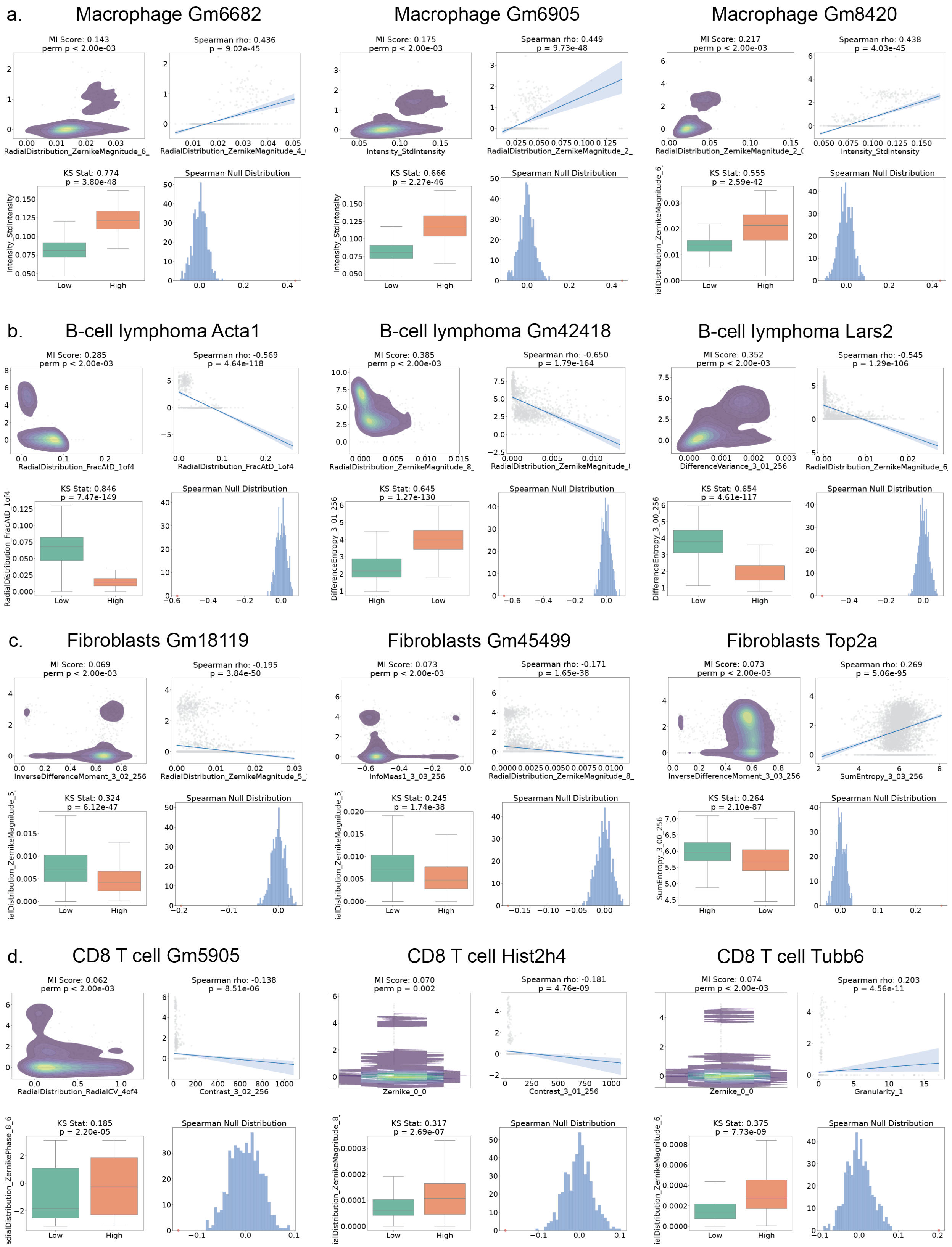
a.
Macrophage Gm6682
Macrophage Gm6905
Macrophage Gm8420
MI Score: 0.143
perm p < 2.00e-03
Spearman rho: 0.436
p = 9.02e-45
KS Stat: 0.774
p = 3.80e-48
Spearman Null Distribution
MI Score: 0.175
perm p < 2.00e-03
Spearman rho: 0.449
p = 9.73e-48
KS Stat: 0.666
p = 2.27e-46
MI Score: 0.217
perm p < 2.00e-03
Spearman rho: 0.438
p = 4.03e-45
KS Stat: 0.555
p = 2.59e-42
b.
B-cell lymphoma Acta1
B-cell lymphoma Gm42418
B-cell lymphoma Lars2
MI Score: 0.285
perm p < 2.00e-03
Spearman rho: -0.569
p = 4.64e-118
KS Stat: 0.846
p = 7.47e-149
MI Score: 0.385
perm p < 2.00e-03
Spearman rho: -0.650
p = 1.79e-164
KS Stat: 0.645
p = 1.27e-130
MI Score: 0.352
perm p < 2.00e-03
Spearman rho: -0.545
p = 1.29e-106
KS Stat: 0.654
p = 4.61e-117
c.
Fibroblasts Gm18119
Fibroblasts Gm45499
Fibroblasts Top2a
MI Score: 0.069
perm p < 2.00e-03
Spearman rho: -0.195
p = 3.84e-50
KS Stat: 0.324
p = 6.12e-47
MI Score: 0.073
perm p < 2.00e-03
Spearman rho: -0.171
p = 1.65e-38
KS Stat: 0.245
p = 1.74e-38
MI Score: 0.073
perm p < 2.00e-03
Spearman rho: 0.269
p = 5.06e-95
KS Stat: 0.264
p = 2.10e-87
d.
CD8 T cell Gm5905
CD8 T cell Hist2h4
CD8 T cell Tubb6
MI Score: 0.062
perm p < 2.00e-03
Spearman rho: -0.138
p = 8.51e-06
KS Stat: 0.185
p = 2.20e-05
MI Score: 0.070
perm p = 0.002
Spearman rho: -0.181
p = 4.76e-09
KS Stat: 0.317
p = 2.69e-07
MI Score: 0.074
perm p < 2.00e-03
Spearman rho: 0.203
p = 4.56e-11
KS Stat: 0.375
p = 7.73e-09
Intensity_StdIntensity
RadialDistribution_FracAtD_1of4
DifferenceVariance_3_01_256
InverseDifferenceMoment_3_02_256
InfoMeas1_3_03_256
InverseDifferenceMoment_3_03_256
SumEntropy_3_03_256
RadialDistribution_RadialCV_4of4
Contrast_3_02_256
Contrast_3_01_256
Zernike_0_0
Granularity_1
Low
High

**Supplementary Figure 19:** Examples of top COSMIC-proposed morphology-related genes within each cell type: **(a)** macrophages, **(b)** B-cell lympphoma, **(c)** fibroblasts, and **(d)** CD8+ T-cells. Representative examples of genes are ranked among the top morphology-associated genes identified by COSMIC in each of the cell types. For each gene, the mutual-information density plot illustrates the relationship between gene expression and the selected CellProfiler-derived morphology feature, where each gray point represents a single cell and colored contours indicate cell-density distributions in feature-expression space. Expression-stratified boxplots compare the morphology feature between low- and high-expression cells. Spearman correlation panels quantify the association between gene expression and the morphology feature, with the blue line indicating the fitted regression trend and the shaded region representing the confidence interval. Permutation tests compare the observed Spearman correlation against a null distribution generated by randomly permuting gene-expression values across cells. These examples highlight top COSMIC-proposed morphology-related genes and demonstrate their associations with specific nuclear morphology features across different cell types.

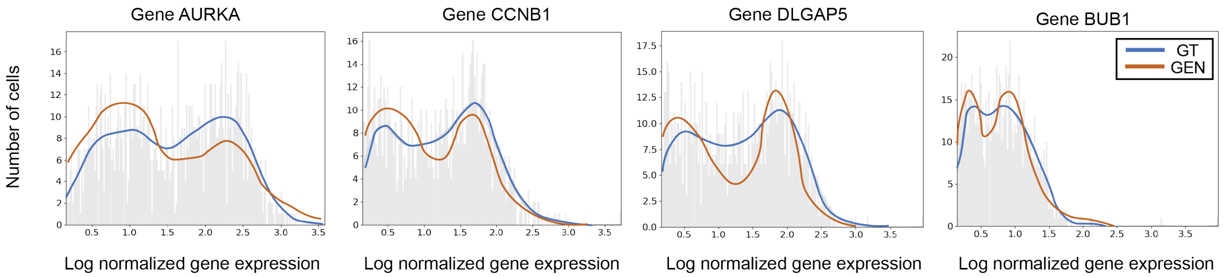


**Supplementary Figure 20:** Expression histograms for additional top morphology-associated genes in DU145 cells, including *AURKA*, *CCNB1*, *DLGAP5*, and *BUB1*. For each gene, the histogram shows its normalized expression level (y-axis) as a function of the number of DU145 cells (x-axis), comparing COSMIC-generated expressions from nuclear morphology (red) with ground-truth expressions (blue).

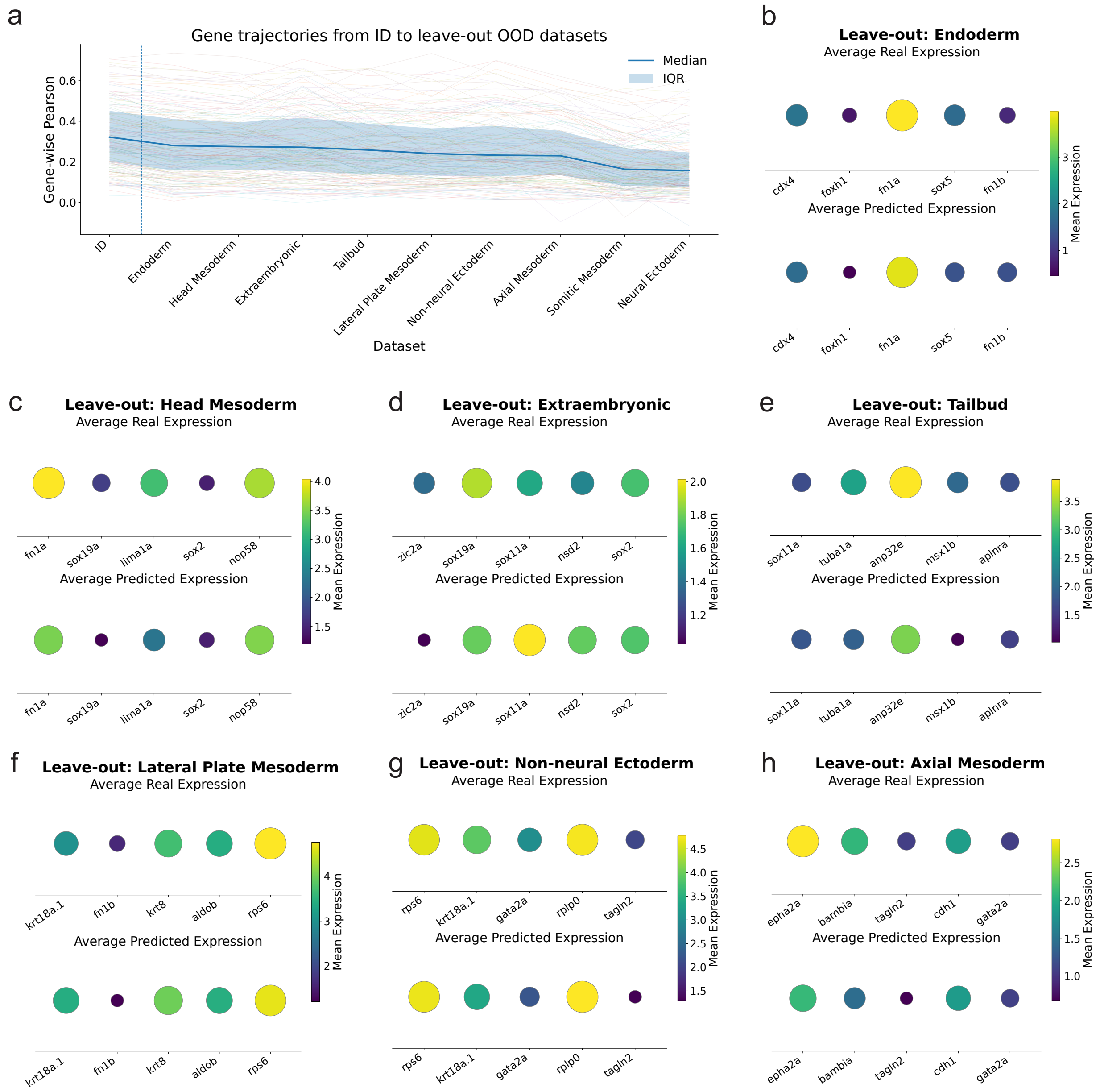
a
Gene trajectories from ID to leave-out OOD datasets
Median
IQR
Gene-wise Pearson
0.6
0.4
0.2
0.0
ID
Endoderm
Head Mesoderm
Extraembryonic
Tailbud
Lateral Plate Mesoderm
Non-neural Ectoderm
Axial Mesoderm
Somitic Mesoderm
Neural Ectoderm
Dataset
b
Leave-out: Endoderm
Average Real Expression
Average Predicted Expression
cdx4
foxh1
fn1a
sox5
fn1b
Mean Expression
c
Leave-out: Head Mesoderm
Average Real Expression
Average Predicted Expression
fn1a
sox19a
lima1a
sox2
nop58
Mean Expression
d
Leave-out: Extraembryonic
Average Real Expression
Average Predicted Expression
zic2a
sox19a
sox11a
nsd2
sox2
Mean Expression
e
Leave-out: Tailbud
Average Real Expression
Average Predicted Expression
sox11a
tuba1a
anp32e
msx1b
aplnra
Mean Expression
f
Leave-out: Lateral Plate Mesoderm
Average Real Expression
Average Predicted Expression
krt18a.1
fn1b
krt8
aldob
rps6
Mean Expression
g
Leave-out: Non-neural Ectoderm
Average Real Expression
Average Predicted Expression
rps6
krt18a.1
gata2a
rplp0
tagln2
Mean Expression
h
Leave-out: Axial Mesoderm
Average Real Expression
Average Predicted Expression
epha2a
bambia
tagln2
cdh1
gata2a
Mean Expression

**Supplementary Figure 21:** Additional held-out-tissue experiments COSMIC's ability to generalize to unseen tissue types in the zebrafish embryo development weMERFISH data. We consider 11 tissue types annotated in the data including endoderm, head mesoderm, axial mesoderm, tailbud, lateral plate mesoderm, extraembryonic, non-neural ectoderm, somitic mesoderm, and neural ectoderm. **(a)** Gene-wise Pearson correlations across in-distribution (ID) and leave-one-tissue-out settings. **(b)-(h)** Comparison between real and COSMIC-predicted average expression for representative genes in held-out tissue types. COSMIC recovers characteristic expression patterns for these genes despite having no training examples from the held-out tissue type, supporting that the model learns transferable morphology-expression structure rather than simply memorizing training data patterns.

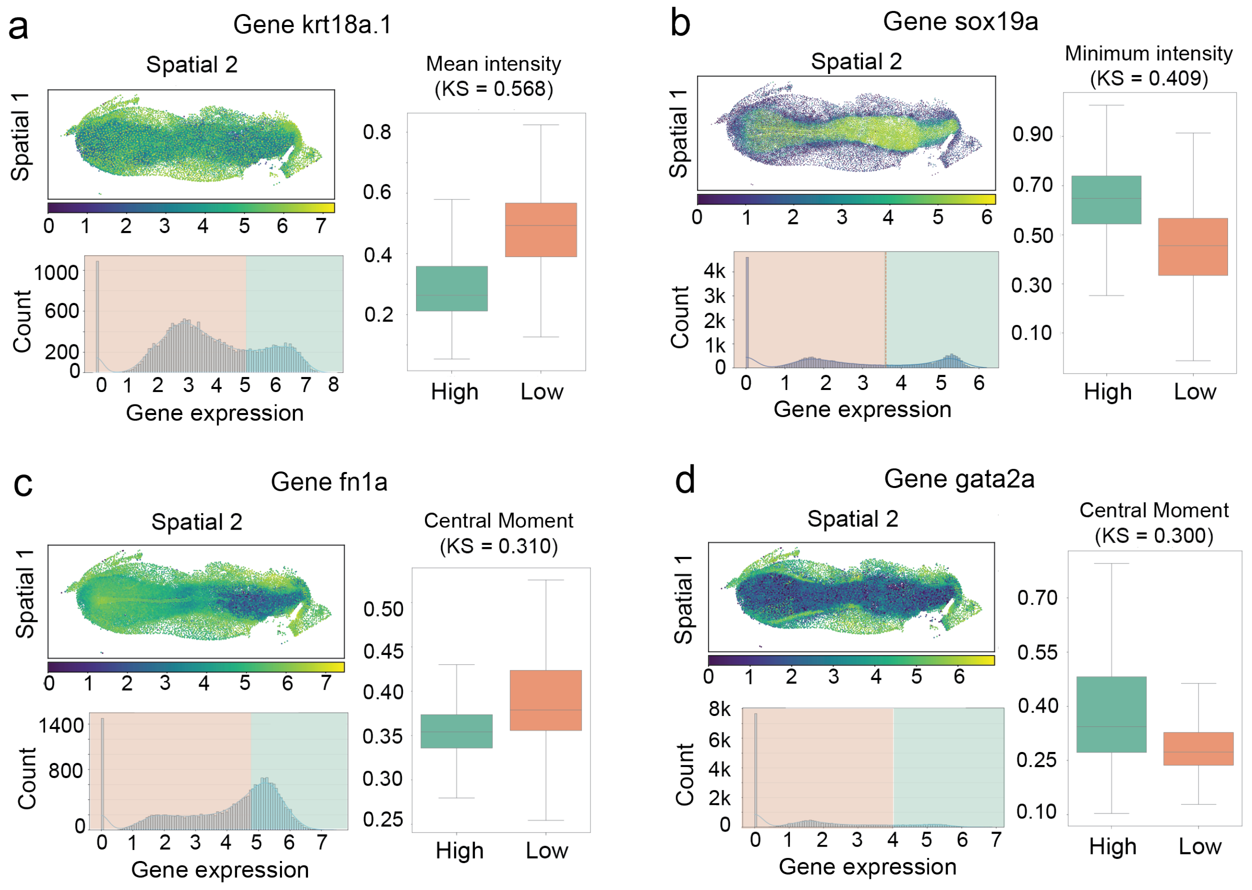


**Supplementary Figure 22:** Additional analysis of COSMIC-identified morphology-associated genes in zebrafish weMERFISH. Additional global morphology-associated genes identified by COSMIC in the zebrafish weMERFISH dataset. Examples include **(a)** *krt18a.1*, **(b)** *sox19a*, **(c)** *fn1a*, and **(d)** *gata2a*. For each gene, spatial expression maps show embryo-wide organization of gene expression (top left), expression histograms indicate the stratification of cells into high- and low-expression groups (bottom left), and CellProfiler-derived feature comparisons show differences in image-derived morphology between these groups (right). Representative morphology features include minimum intensity and central moment. These examples provide additional support that COSMIC-nominated genes are associated with spatially organized and measurable image-feature variation in the embryo.

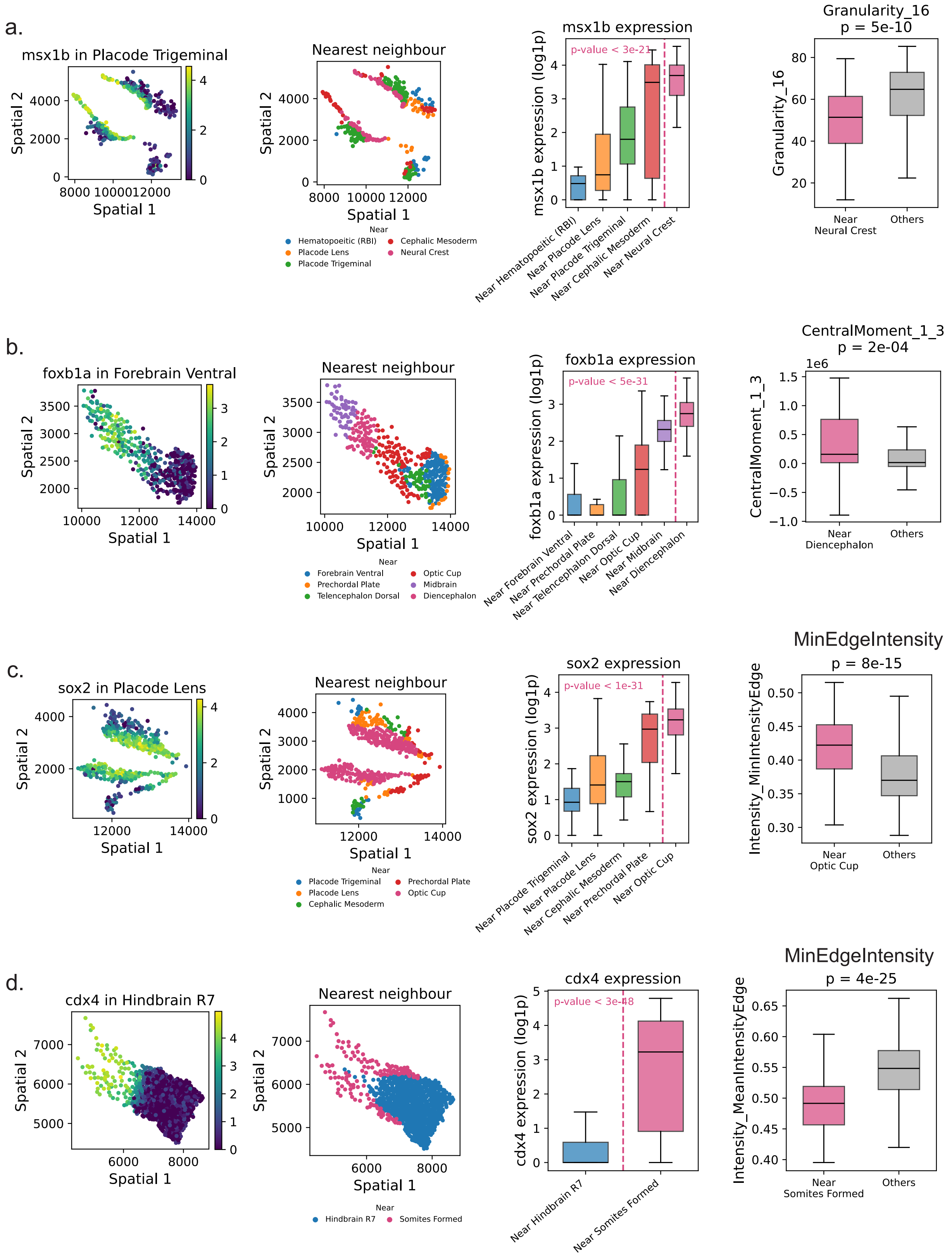

a.
msx1b in Placode Trigeminal
Spatial 2
Spatial 1
Nearest neighbour
Near
Hematopoeitic (RBI)
Placode Lens
Placode Trigeminal
Cephalic Mesoderm
Neural Crest
msx1b expression
p-value < 3e-21
msx1b expression (log1p)
Near Hematopoeitic (RBI)
Near Placode Lens
Near Placode Trigeminal
Near Cephalic Mesoderm
Near Neural Crest
Granularity_16
p = 5e-10
Near Neural Crest
Others
b.
foxb1a in Forebrain Ventral
Nearest neighbour
Near
Forebrain Ventral
Prechordal Plate
Telencephalon Dorsal
Optic Cup
Midbrain
Diencephalon
foxb1a expression
p-value < 5e-31
foxb1a expression (log1p)
Near Forebrain Ventral
Near Prechordal Plate
Near Telencephalon Dorsal
Near Optic Cup
Near Midbrain
Near Diencephalon
CentralMoment_1_3
p = 2e-04
1e6
Near Diencephalon
Others
c.
sox2 in Placode Lens
Nearest neighbour
Near
Placode Trigeminal
Placode Lens
Cephalic Mesoderm
Prechordal Plate
Optic Cup
sox2 expression
p-value < 1e-31
sox2 expression (log1p)
Near Placode Trigeminal
Near Placode Lens
Near Cephalic Mesoderm
Near Prechordal Plate
Near Optic Cup
MinEdgeIntensity
p = 8e-15
Intensity_MinIntensityEdge
Near Optic Cup
Others
d.
cdx4 in Hindbrain R7
Nearest neighbour
Near
Hindbrain R7
Somites Formed
cdx4 expression
p-value < 3e-48
cdx4 expression (log1p)
Near Hindbrain R7
Near Somites Formed
MinEdgeIntensity
p = 4e-25
Intensity_MeanIntensityEdge
Near Somites Formed
Others

**Supplementary Figure 23:** Four examples of genes nominated by COSMIC as morphology-predictable within one tissue type in the zebrafish embryo weMERFISH dataset: **(a)** *msx1b* in placode trigeminal, **(b)** *foxb1a* in ventral, **(c)** *sox2* in placode lens, and **(d)** *cdx4* in hindbrain R7. In each panel, cells of the indicated tissue are grouped by the identity of their neighboring tissue type (majority of the 100 nearest spatial neighbors, restricted to the tissues adjacent to the target tissue). From left to right: cells within a tissue type in spatial coordinates colored by gene expression (log1p) (most left); same cells colored by nearest-neighbor group (middle left); gene expression across neighbor groups, with the highlighted standout group separated by the dashed line (middle right); and the most strongly differing nuclear (DAPI) morphology feature for the standout-neighbor group versus all other cells of the tissue (right). Spatial axes are shown on a common square layout. Expression differences are between the standout-neighbor group and the remaining cells of the tissue (two-sided Mann–Whitney $U$ test); reported means are group means on the log1p scale. **(a)** Trigeminal placode, *msx1b*: cells near Neural Crest show higher *msx1b* than other Trigeminal placode cells (3.50 vs 1.74; $p < 3 \times 10^{-21}$) and differ in nuclear texture (granularity; $p < 5 \times 10^{-10}$), consistent with the role of *msx* genes at the neural-plate border in cranial placode and neural crest development [10,11]. **(b)** Ventral forebrain, *foxb1a*: cells near Diencephalon show higher *foxb1a* (2.66 vs 0.81; $p < 5 \times 10^{-31}$) and differ in nuclear shape (central moment; $p < 2 \times 10^{-4}$), consistent with the restricted expression of *foxb1* in the caudal/ventral diencephalon and midbrain [12]. **(c)** Lens placode, *sox2*: cells near the Optic Cup show higher *sox2* (3.09 vs 1.73; $p < 1 \times 10^{-31}$) and differ in nuclear edge intensity ($p < 8 \times 10^{-15}$), consistent with induction of *Sox2* in head ectoderm contacted by the optic vesicle during lens placode formation [13,25]. **(d)** Posterior hindbrain (R7), *cdx4*: cells near formed somites show higher *cdx4* (2.56 vs 0.40; $p < 3 \times 10^{-48}$) and differ in nuclear edge intensity ($p < 4 \times 10^{-25}$), consistent with the role of Cdx4 in posterior positional identity and spinal cord specification at the hindbrain–spinal cord transition [14,15].

| Cell line | This work (expIDs) | From IRIS work [26] (expIDs) |
|---|---|---|
| *Human* | | |
| DU145 | JP240 | - |
| PBMC | JP247 | NG037, NG039, NG050, NG042, JP250, JP268, NG045, NG027 |
| RPE | JP270, NG046 | - |
| C4-2B | JP273, NG047 | - |
| *Mouse* | | |
| 3T3 | - | JP241, JP277, NG015, NG026, NG029 |
| Naive CD8+ | JP263 | - |
| CAR-A20 | JP271 | - |
| RAW | NG033 | - |

**Supplementary Table 1:** Batches by source for each cell line.

| Cell type | Top genes (well-studied) | Top genes (expected by function) | Top genes (non-obvious, known function) | Top genes (poorly characterized) |
|---|---|---|---|---|
| Fibroblast | *Prc1*, *Top2a*, *Stag2*, *Hist1h4b*, *Kif11*, *Nusap1*, *Cenpe*, *Matr3*, *Cenpf*, *Hist1h4c*, *Kif20b*, *Spc25* | *Rasl2-9*, *Arl6ip1*, *Tuba3b*, *Dsp*, *Arhgap24*, *Rab7b*, *Rtn4*, *Mcm7*, *Tubb2b*, *Tubg1* | *Acacb*, *Ndufaf3*, *Ctss*, *Chchd10*, *Ldhb*, *Hnrnph3*, *Caprin2*, *Sox2ot*, *Wdr89*, *Cd80* | *Gm18119*, *Gm45499*, *Gm5305*, *AC129085.5*, *Gm6218*, *Gm16073*, *Gm11810*, *Gm13641*, *Gm7424*, *Gm8805*, |
| Macrophage | *Hmgn3*, *Hist4h4*, *Hist1h4c*, *Hist1h2ac*, *Hist1h4b*, *Cdkn3*, *Hist1h1c*, *Kif11*, *Hist1h3a*, *Kpnb1* | *Marcks*, *Capg*, *Myh10*, *Rasl2-9*, *Enah*, *Rnd3*, *Krt10*, *Rad51c*, *Chek1*, *Iqgap2*, *Tubgcp2*, *Cald1*, *Hdac2* | *Sema5a*, *Hydin*, *Mzt2*, *Gstp3*, *Cluap1*, *Il1rapl1*, *Chchd10*, *Smoc1*, *Ldhb*, *Pcdh9*, *Caprin2* | *Gm6682*, *Gm6905*, *Gm8420*, *Gm7434*, *Gm10320*, *Gm6341*, *Gm13641*, *Gm5384*, *Gm13194*, *Gm12739*, |
| naive $CD8^+$ | *Hist2h4*, *Hist1h4b*, *Hist1h4h*, *Ran*, *H2afz*, *Npm1*, *Ncl*, *Hmgn5*, *Hmgn1*, *Hist1h4c* | *Tubb6*, *Acta2*, *Actg2*, *Tubb4a*, *Tuba3b*, *Sar1a*, *Tmed4*, *Acta1*, *Tubb2b* | *Mir6236*, *Cmss1*, *Ubb*, *Hspa2*, *Il31ra*, *Eno3*, *Lars2*, *Ppif*, *Elp3*, *Ywhag* | *Gm5905*, *AC129085.5*, *Gm8407*, *Gm42418*, *Gm5735*, *Gm10689*, *Gm6436*, *Gm19620*, *Gm10320*, *CT010467.1* |
| B-cell lymphoma | *Hist1h4f*, *H2afx*, *Birc5*, *Ehmt1*, *Nipbl*, *Emd*, *Nup50*, *Nup98*, *Smc3*, *Nup85* | *Acta1*, *Acta2*, *Tubb2b*, *Tubb4a*, *Reep5*, *Rpn2*, *Snx29*, *Rab8a*, *Dynlt3*, *Dync1li1* | *Lars2*, *Ptk2*, *Peak1*, *Mef2b*, *Rasgrp3*, *Slc15a2*, *Pkig*, *Nrg1*, *Epha7*, *Ubb* | *Gm42418*, *Gm15564*, *Gm26917*, *Gm10039*, *Gm15772*, *Gm14149*, *Gm26870*, *Gm3448*, *Gm15459*, *Gm8730* |

**Supplementary Table 2:** Mouse cell-type-specific morphology-associated genes grouped by prior biological expectedness. The categories were assigned using an LLM-assisted literature annotation procedure. GPT-5.6 Thinking [27] and Claude Opus 4.8 [28] independently classified each top-ranked gene into one of four categories: well-studied morphology-related genes, genes expected to be morphology-related based on known function, non-obvious genes with characterized functions, and poorly characterized genes. Each model was asked to provide supporting literature and identify the specific evidence underlying its assignment. The two independent annotation sets were then provided to both models for cross-review and reconciliation, and the resulting consensus assignments are reported here.

| Cell type | Top genes (well-studied) | Top genes (expected by function) | Top genes (non-obvious, known function) | Top genes (poorly characterized) |
|---|---|---|---|---|
| PCa-LN | *HIST1H4A*, *HIST1H4I*, *AURKB*, *NCAPD2*, *SMC3*, *NIPBL*, *HIST1H4D*, *SGO1*, *KNL1*, *CBX5* | *TUBB4A*, *TUBA4A*, *ACTA1*, *ACTA2*, *ARHGEF38*, *CYLC1*, *TUBA3C*, *TUBA1A*, *KIF15*, *KIF18B* | *ANKRD30BL*, *ENO3*, *SAMD3*, *HSPA2*, *UBC*, *UBA52*, *ANAPC11*, *ATP5H*, *ANKRD28*, *SYN3* | *AL080243.2*, *AC068228.1*, *AL050331.1*, *AL355472.2*, *AL451074.5*, *AC087672.3*, *LINC01505*, *AC099535.1*, *AC010234.1*, *AC073861.1* |
| PCa-Br | *AURKA*, *DLGAP5*, *BUB1*, *KNSTRN*, *CDCA8*, *HJURP*, *KIF23*, *NDC80*, *PSRC1*, *PLK1*, *KIF14* | *CKS2*, *CCNB1*, *ARHGEF39*, *ARHGAP11A*, *CCNF*, *RAB8A*, *CCNA2*, *CEP55*, *MARCKS*, *KIF4A*, *RAD51B* | *BRD8*, *FXR1*, *PRR11*, *SNHG19*, *CASC9*, *ZNF804A*, *HES1*, *GPC6*, *SOGA1*, *CANT1* | *FP671120.1*, *AC134511.1*, *LINC01036*, *AC024230.1*, *AC099850.2*, *LINC00470*, *LINC01037*, *FP236383.1*, *AC084033.3*, *GABPB1-AS1* |
| Lymphocytes | *HIST1H4A*, *HIST1H4I*, *HIST1H3F*, *HIST1H3E*, *HIST1H4D*, *HIST1H3J*, *HIST1H3B*, *SYNE1*, *SYNE2*, *NAP1L1* | *TUBB4A*, *TUBB4B*, *TUBA4A*, *ACTA2*, *ACTA1*, *TUBA3D*, *TMED4*, *TUBA3C*, *RHOB*, *AP2S1* | *YWHAG*, *ANKRD30BL*, *ENO3*, *PPIF*, *PPIC*, *YWHAH*, *NME3*, *HSPA2*, *SUMO3*, *CALML3* | *AC068228.1*, *AL355472.2*, *AL080243.2*, *AC010234.1*, *LINC00486*, *AL451074.5*, *AL050331.1*, *AC090589.1*, *AC007969.1*, *AC073861.1* |
| Myeloids | *SYNE2*, *NCAPD3*, *NDC1*, *CKAP2*, *PRC1*, *CKAP5*, *HIST1H4A*, *SAFB*, *NUP98*, *GLE1* | *ABLIM1*, *MAP2K4*, *SEC24A*, *CHD3*, *SSR1*, *RAB29*, *DAAM1*, *BAZ1A*, *ACTN4*, *DOCK4* | *NSUN6*, *TWSG1*, *TCF7*, *IL32*, *PRKCG*, *ARL4C*, *YTHDC2*, *REXO4*, *KAT8*, *SF3B2* | *ASMTL-AS1*, *THUMPD3-AS1*, *AL050403.2*, *AL590867.2*, *AL356599.1*, *AL365273.2*, *AL445524.1*, *AC092164.1*, *AL669831.5*, *AC027018.1* |
| RPE | *HIST1H4A*, *HIST1H4I*, *HMGB1*, *CIT*, *CKAP2L*, *HIST1H3J*, *TMPO*, *CKAP2*, *KPNA2*, *ASPM* | *TUBB4A*, *ACTA1*, *CYLC1*, *TUBA4A*, *ARHGEF38*, *TUBA3C*, *ACTA2*, *CDK1*, *CCNA2*, *KRT8* | *CNTN5*, *ANKRD30BL*, *NME3*, *ZMAT3*, *PAPSS2*, *UBC*, *HSPA2*, *CLSTN2*, *LIMA1*, *TFCP2L1* | *AC068228.1*, *AL080243.2*, *AC099535.1*, *AC087672.3*, *LINC00486*, *AL050331.1*, *AL355472.2*, *AC010234.1*, *AL451074.5*, *LINC01505* |

**Supplementary Table 3:** Human cell-type-specific morphology-associated genes grouped by prior biological expectedness. The categories were assigned using an LLM-assisted literature annotation procedure. GPT-5.6 Thinking [27] and Claude Opus 4.8 [28] independently classified each top-ranked gene into one of four categories: well-studied morphology-related genes, genes expected to be morphology-related based on known function, non-obvious genes with characterized functions, and poorly characterized genes. Each model was asked to provide supporting literature and identify the specific evidence underlying its assignment. The two independent annotation sets were then provided to both models for cross-review and reconciliation, and the resulting consensus assignments are reported here.

| Rank | `CellProfiler feature` | FDR-adjusted $p$ | Rank | `CellProfiler feature` | FDR-adjusted $p$ |
|---|---|---|---|---|---|
| 1 | `Intensity_IntegratedIntensityEdge` | $2.31 \times 10^{-72}$ | 26 | `SpatialMoment_0_0` | $1.50 \times 10^{-63}$ |
| 2 | `Intensity_IntegratedIntensity` | $1.51 \times 10^{-68}$ | 27 | `FilledArea` | $1.50 \times 10^{-63}$ |
| 3 | `SpatialMoment_2_0` | $4.64 \times 10^{-68}$ | 28 | `CentralMoment_0_0` | $1.50 \times 10^{-63}$ |
| 4 | `Perimeter` | $4.69 \times 10^{-68}$ | 29 | `Area` | $1.50 \times 10^{-63}$ |
| 5 | `BoundingBoxArea` | $3.00 \times 10^{-67}$ | 30 | `EquivalentDiameter` | $1.50 \times 10^{-63}$ |
| 6 | `Entropy_3_02_256` | $3.00 \times 10^{-67}$ | 31 | `SpatialMoment_1_3` | $2.23 \times 10^{-63}$ |
| 7 | `PerimeterCrofton` | $3.18 \times 10^{-67}$ | 32 | `MeanRadius` | $3.02 \times 10^{-63}$ |
| 8 | `Entropy_3_03_256` | $3.24 \times 10^{-67}$ | 33 | `InertiaTensor_1_1` | $7.69 \times 10^{-63}$ |
| 9 | `CentralMoment_2_0` | $3.48 \times 10^{-67}$ | 34 | `MedianRadius` | $2.00 \times 10^{-62}$ |
| 10 | `AngularSecondMoment_3_03_256` | $3.75 \times 10^{-67}$ | 35 | `AngularSecondMoment_3_00_256` | $4.76 \times 10^{-62}$ |
| 11 | `SpatialMoment_2_1` | $3.84 \times 10^{-67}$ | 36 | `MinorAxisLength` | $6.68 \times 10^{-62}$ |
| 12 | `MaxFeretDiameter` | $4.74 \times 10^{-67}$ | 37 | `InertiaTensorEigenvalues_1` | $6.68 \times 10^{-62}$ |
| 13 | `Entropy_3_01_256` | $5.99 \times 10^{-67}$ | 38 | `MinFeretDiameter` | $4.76 \times 10^{-61}$ |
| 14 | `SpatialMoment_2_2` | $9.55 \times 10^{-67}$ | 39 | `SpatialMoment_0_1` | $1.04 \times 10^{-60}$ |
| 15 | `SpatialMoment_1_0` | $9.55 \times 10^{-67}$ | 40 | `CentralMoment_0_2` | $4.00 \times 10^{-60}$ |
| 16 | `Entropy_3_00_256` | $3.77 \times 10^{-66}$ | 41 | `MaximumRadius` | $6.02 \times 10^{-60}$ |
| 17 | `AngularSecondMoment_3_02_256` | $4.05 \times 10^{-66}$ | 42 | `SpatialMoment_0_2` | $1.04 \times 10^{-58}$ |
| 18 | `MajorAxisLength` | $8.44 \times 10^{-66}$ | 43 | `SpatialMoment_0_3` | $1.48 \times 10^{-58}$ |
| 19 | `InertiaTensorEigenvalues_0` | $8.44 \times 10^{-66}$ | 44 | `InertiaTensor_0_0` | $1.29 \times 10^{-54}$ |
| 20 | `SpatialMoment_1_1` | $1.09 \times 10^{-65}$ | 45 | `Granularity_7` | $2.12 \times 10^{-48}$ |
| 21 | `ConvexArea` | $4.67 \times 10^{-65}$ | 46 | `Granularity_16` | $7.96 \times 10^{-48}$ |
| 22 | `SpatialMoment_2_3` | $7.88 \times 10^{-65}$ | 47 | `InfoMeas1_3_03_256` | $3.76 \times 10^{-47}$ |
| 23 | `CentralMoment_2_2` | $1.83 \times 10^{-64}$ | 48 | `Granularity_15` | $4.22 \times 10^{-47}$ |
| 24 | `AngularSecondMoment_3_01_256` | $1.83 \times 10^{-64}$ | 49 | `Granularity_6` | $5.61 \times 10^{-46}$ |
| 25 | `SpatialMoment_1_2` | $1.50 \times 10^{-63}$ | 50 | `InfoMeas1_3_01_256` | $5.78 \times 10^{-45}$ |

**Supplementary Table 4:** Top 50 CellProfiler features distinguishing cells with high versus low *Marcks* expression. Features are ranked by the two-sided Kolmogorov-Smirnov statistic, and the corresponding FDR-adjusted $p$-values are reported.

| Shared feature | Shared feature | Shared feature |
|---|---|---|
| AngularSecondMoment_3_00_256 | EquivalentDiameter | InverseDifferenceMoment_3_02_256 |
| AngularSecondMoment_3_01_256 | Extent | InverseDifferenceMoment_3_03_256 |
| AngularSecondMoment_3_02_256 | FilledArea | MajorAxisLength |
| AngularSecondMoment_3_03_256 | Granularity_10 | MaxFeretDiameter |
| Area | Granularity_11 | MaximumRadius |
| BoundingBoxArea | Granularity_12 | MeanRadius |
| BoundingBoxMaximum_X | Granularity_13 | MedianRadius |
| BoundingBoxMaximum_Y | Granularity_14 | MinFeretDiameter |
| BoundingBoxMinimum_X | Granularity_15 | MinorAxisLength |
| BoundingBoxMinimum_Y | Granularity_16 | Perimeter |
| CentralMoment_0_0 | Granularity_2 | PerimeterCrofton |
| CentralMoment_0_1 | Granularity_3 | RadialDistribution_MeanFrac_2of4 |
| CentralMoment_0_2 | Granularity_4 | RadialDistribution_MeanFrac_4of4 |
| CentralMoment_0_3 | Granularity_5 | RadialDistribution_ZernikeMagnitude_0_0 |
| CentralMoment_1_0 | Granularity_6 | RadialDistribution_ZernikeMagnitude_7_3 |
| CentralMoment_1_1 | Granularity_7 | RadialDistribution_ZernikeMagnitude_8_2 |
| CentralMoment_1_2 | Granularity_8 | Solidity |
| CentralMoment_1_3 | Granularity_9 | SpatialMoment_0_0 |
| CentralMoment_2_0 | InertiaTensorEigenvalues_0 | SpatialMoment_0_1 |
| CentralMoment_2_1 | InertiaTensorEigenvalues_1 | SpatialMoment_0_2 |
| CentralMoment_2_2 | InertiaTensor_0_0 | SpatialMoment_0_3 |
| CentralMoment_2_3 | InertiaTensor_1_1 | SpatialMoment_1_0 |
| Contrast_3_00_256 | InfoMeas1_3_00_256 | SpatialMoment_1_1 |
| Contrast_3_01_256 | InfoMeas1_3_01_256 | SpatialMoment_1_2 |
| Contrast_3_02_256 | InfoMeas1_3_02_256 | SpatialMoment_1_3 |
| Contrast_3_03_256 | InfoMeas1_3_03_256 | SpatialMoment_2_0 |
| ConvexArea | InfoMeas2_3_00_256 | SpatialMoment_2_1 |
| Correlation_3_00_256 | InfoMeas2_3_01_256 | SpatialMoment_2_2 |
| Correlation_3_01_256 | InfoMeas2_3_02_256 | SpatialMoment_2_3 |
| Correlation_3_02_256 | InfoMeas2_3_03_256 | SumAverage_3_00_256 |
| Correlation_3_03_256 | Intensity_IntegratedIntensity | SumAverage_3_01_256 |
| DifferenceEntropy_3_00_256 | Intensity_IntegratedIntensityEdge | SumAverage_3_02_256 |
| DifferenceEntropy_3_01_256 | Intensity_LowerQuartileIntensity | SumAverage_3_03_256 |
| DifferenceEntropy_3_02_256 | Intensity_MaxIntensityEdge | SumEntropy_3_00_256 |
| DifferenceEntropy_3_03_256 | Intensity_MeanIntensity | SumEntropy_3_01_256 |
| DifferenceVariance_3_00_256 | Intensity_MeanIntensityEdge | SumEntropy_3_02_256 |
| DifferenceVariance_3_01_256 | Intensity_MedianIntensity | SumEntropy_3_03_256 |
| DifferenceVariance_3_02_256 | Intensity_MinIntensity | Variance_3_00_256 |
| DifferenceVariance_3_03_256 | Intensity_MinIntensityEdge | Variance_3_01_256 |
| Entropy_3_00_256 | Intensity_UpperQuartileIntensity | Variance_3_02_256 |
| Entropy_3_01_256 | InverseDifferenceMoment_3_00_256 | Variance_3_03_256 |
| Entropy_3_02_256 | InverseDifferenceMoment_3_01_256 | Zernike_7_3 |
| Entropy_3_03_256 | | |

**Supplementary Table 5:** Significant CellProfiler features shared by *Sema5a*, *Capg*, and *Myh10*.
